# Thermodynamic modeling of Csr/Rsm- RNA interactions capture novel, direct binding interactions across the *Pseudomonas aeruginosa* transcriptome

**DOI:** 10.1101/2024.08.01.606018

**Authors:** Alexandra J Lukasiewicz, Abigail N Leistra, Lily Hoefner, Erika Monzon, Cindy J Gode, Bryan T Zorn, Kayley H Janssen, Timothy L Yahr, Matthew C Wolfgang, Lydia M Contreras

**Affiliations:** Department of Molecular Biosciences, The University of Texas at Austin, Austin, TX; McKetta Department of Chemical Engineering, The University of Texas at Austin; Department of Biology, The University of Texas at Austin, Austin, TX; Marsico Lung Institute, University of North Carolina at Chapel Hill, Chapel Hill, NC; Department of Microbiology and Immunology, University of Iowa, Iowa City, Iowa, USA; Bellin College, Green Bay, WI; Department of Microbiology and Immunology, University of North Carolina at Chapel Hill, Chapel Hill, NC

**Keywords:** Post-transcriptional regulation, Computational modeling, RNA regulation, RNA-binding proteins, Regulatory networks, Transcriptional control, Systems biology, RNA-protein interactions, RNA secondary structure, Pseudomonas aeruginosa

## Abstract

**Background:** *Pseudomonas aeruginosa* (PA) is a ubiquitous, Gram-negative, bacteria that can attribute its survivability to numerous sensing and signaling pathways; conferring fitness due to speed of response. Post-transcriptional regulation is an energy efficient approach to quickly shift gene expression in response to the environment. The conserved post-transcriptional regulator RsmA is involved in regulating translation of genes involved in pathways that contribute to virulence, metabolism, and antibiotic resistance. Prior high-throughput approaches to map the full regulatory landscape of RsmA have estimated a target pool of approximately 500 genes; however, these approaches have been limited to a narrow range of growth phase, strain, and media conditions. Computational modeling presents a condition-independent approach to generating predictions for binding between the RsmA protein and highest affinity mRNAs. In this study, we draft a two-state thermodynamic model to predict the likelihood of RsmA binding to the 5’ UTR sequence of genes present in the PA genome.

**Results:** Our modeling approach predicts 1043 direct RsmA-mRNA binding interactions, including 457 novel mRNA targets. We then perform GO term enrichment tests on our predictions that reveal significant enrichment for DNA binding transcriptional regulators. In addition, quorum sensing, biofilm formation, and two-component signaling pathways were represented in KEGG enrichment analysis. We confirm binding predictions using *in vitro* binding assays, and regulatory effects using *in vivo* translational reporters. These reveal RsmA binding and regulation of a broader number of genes not previously reported. An important new observation of this work is the direct regulation of several novel mRNA targets encoding for factors involved in Quorum Sensing and the Type IV Secretion system, such as *rsaL* and *mvaT*.

**Conclusions:** Our study demonstrates the utility of thermodynamic modeling for predicting interactions independent of complex and environmentally-sensitive systems, specifically for profiling the post-transcriptional regulator RsmA. Our experimental validation of RsmA binding to novel targets both supports our model and expands upon the pool of characterized target genes in PA. Overall, our findings demonstrate that a modeling approach can differentiate direct from indirect binding interactions and predict specific sites of binding for this global regulatory protein, thus broadening our understanding of the role of RsmA regulation in this relevant pathogen.

## Background

*Pseudomonas aeruginosa* (PA) is a widespread, opportunistic pathogen that contributes to nosocomial infection and mortality in immunocompromised individuals. Critical to pathogenesis is the ability of PA to rapidly alter gene expression to respond to the environment. The post-transcriptional regulator RsmA, a member of the CsrA family of RNA-binding proteins (RBPs), achieves this rapid response via post-transcriptional regulation. RsmA is a 6.9 kDa homodimeric protein whose regulatory influence is of clinical relevance as it regulates the expression of genes involved in motility, cell adhesion [1], biofilm formation [2], and secretion of effector proteins [1].

The mechanism by which Rsm/Csr family proteins repress translation is by blocking ribosomal pairing to the Ribosome Binding Site (RBS) present in the 5’ untranslated region (UTR) of an mRNA [3,4]. This can occur through direct binding to and occlusion of the RBS sequence, or through binding in adjacent regions that result in structural rearrangement that reduces [2] or increases [5] accessibility of the RBS. In PA, the RsmA protein exerts tight control of pathways associated with planktonic colonization and sessile biofilm forming states [6]. In addition, the CsrA paralog RsmF/N[7,8] also binds and regulates overlapping [9] and exclusive [10] genes relative to RsmA. The regulatory activity of RsmA itself is sensitive to control by the GacA/GacS two-component signaling (TCS) pathway, which activates expression of antagonistic sRNA sponges RsmY and RsmZ [11] that sequester the RsmA protein. Upon sequestration by these sRNA sponges, the regulatory effect of RsmA is inhibited and produces an inverse effect on translation of directly bound mRNAs. RsmA binds and regulates genes globally throughout the transcriptome. RsmA knockout results in large phenotypic changes to the cell including decreased infection phenotypes [12], impedes active colonization, and promotion of chronic infection states [13].

Full characterization of the binding repertoire of a post-transcriptional regulator, such as RsmA, is difficult to adequately capture using a single high throughput approach [14]. Wide variety in gene expression and regulatory effects have been observed for Csr/Rsm family proteins due to various stresses or infectious states [15,16]. This is partially due to the fact that the pathways that govern the cellular transition from active colonization to chronic biofilm forming states are complex, deeply interlinked, and sensitive to the experimental contexts they are studied in [17].

Efforts to experimentally map the regulatory influence of RsmA range from broad, high throughput sequencing screens to individual *in vitro* biochemical assays. Overall, these high throughout approaches have estimated a target pool of approximately 500 genes that are either directly or indirectly regulated by RsmA [1,9,10,18,19]. Direct binding has been biochemically confirmed *in vitro* for fewer than 2% of this estimated pool of 500 genes. To date, confirmed direct bound mRNA targets of RsmA include *tssA1, fha1, magA* [1]*, psl* [2]*, rahU, algU, pqsR, hxuI* [20]*, mucA* [9], and *retS* [21].

While sequencing approaches have been valuable for understanding the breadth of regulation influenced by the Gac/Rsm pathway, they may not capture potential targets due to low gene expression, strain to strain variation, condition dependent expression, heterogenous expression, sample manipulation, or high limits of detection. For example, microarray, RNA-seq, and proteomic screens fall short when assessing whether post-transcriptional regulation is occurring in a direct (i.e. direct binding of RBP to transcript) or indirect (i.e. network) manner. RNA-seq based approaches can also lose detection of transcripts that are not always degraded when bound by a post-transcriptional regulator, which convolute differential expression-based analyses; thus, missing potential targets of the protein [22]. In contrast, cross-linking immunoprecipitation (CLIP) and RNA immunoprecipitation (RIP) sequencing approaches can identify more direct binding interactions; however, data resulting from these techniques lose positional resolution for mRNA binding sites for small proteins like RsmA. In addition, cross-linking can introduce false positives due to nonspecific linkages between the protein of interest and nearby RNA. Finally, many available high throughput datasets are limited to a narrow range of growth phase, strain, and media conditions that do not capture the full diversity of conditions the organism experiences natively. This presents a bottleneck in discovery, as gene expression varies widely across experimental conditions [23] and can be influenced by extensive strain diversity [24,25].

Computational modeling offers a condition-independent method for predicting binding partners of globally binding proteins. Thermodynamic models of protein-RNA interactions have demonstrated high predictive capabilities, such as that for the PUF4 protein interactions in *Saccharomyces* cerevisiae [26]. Similarly, thermodynamic models to predict binding and translation rates for ribosomes [27,28] have been used for both prediction of native translation and forward design of effective RBS sequences [29]. Although the small handful of confirmed, direct RsmA targets limits the ability to generate accurate models of binding using learning algorithms, much more data of direct targets has been collected for its closely related protein CsrA as genome wide screens have been performed to predict binding sites of the CsrA protein. In 2014, a sequence-based model was crafted for the Csr/Rsm family proteins to identify potential targets within transcriptomes of *E. coli, P. aeruginosa, L. pneumatophilia,* and *S. enterocolitica* [20]. In this work, we improve upon this approach by crafting a biophysical model of interaction built upon additional molecular features that influence binding which yields an energetic prediction for the probability of an interaction between RsmA and an mRNA in *P. aeruginosa*

The *Escherichia coli* CsrA protein has been shown to be well suited for construction of a biophysical model of protein-RNA binding with characterized, empirically-derived, parameters [30], as core elements of binding mechanism that governs its post-transcriptional regulatory effect have been biochemically assessed. These principal rules of interaction include (1) the clear definition of a core ANGGA binding motif [31], (2) the energetic contribution of individual nucleotides within the core motif, (3) establishing a minimal distance between binding sites to reduce steric hindrance within the homodimer [4], and (4) position of binding within stem loop structures of the bound RNA [31] (**Fig. 1a).** Previously, these core rules were leveraged to craft a biophysical model to observe binding patterns of the CsrA protein in *E. coli* [30] which yielded insights in the various molecular features that influence CsrA binding to 236 mRNAs [30]

**Figure 1:**
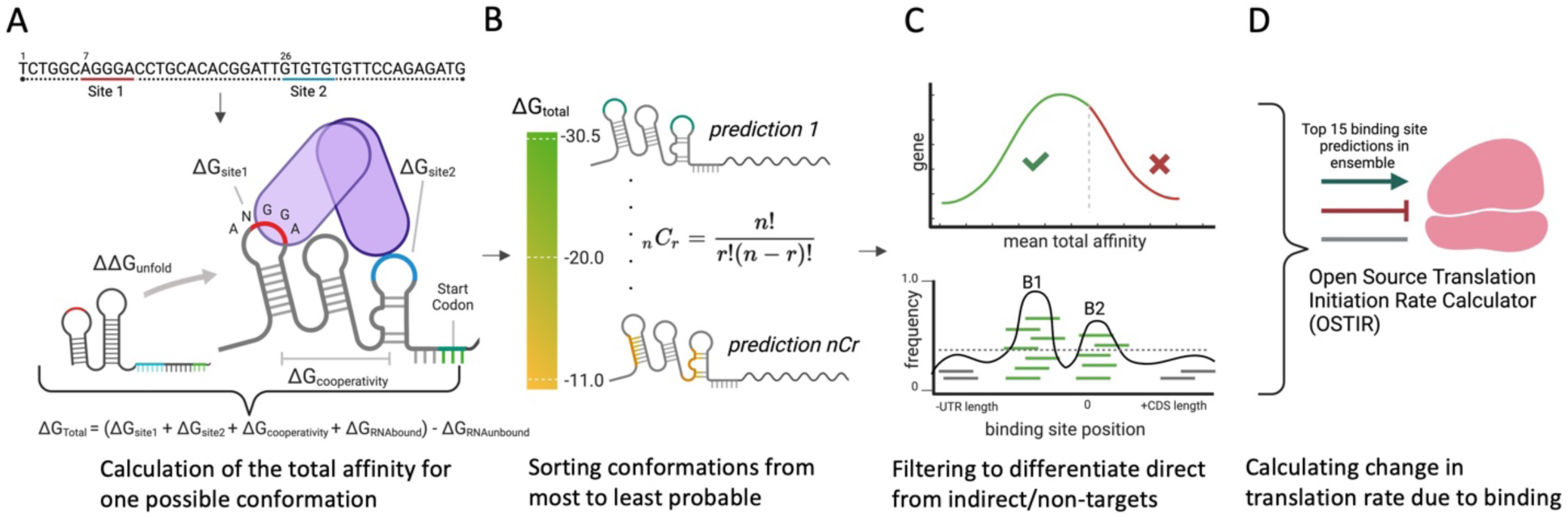
Overview of energy parameters and procedure of the RsmA biophysical model. A) Core energy terms define the energetic parameters of RsmA binding to a specific RNA sequence and model the change in free energy (ΔG_total_) of the system from an unbound to a bound state. B) Each pair of binding site predictions are evaluated across the entire sequence space per gene, and sorted from most to lease probable given the free energy of binding ΔG_total_. C) Predictions are filtered given favorability of the change in free energy (ΔGtotal) and frequency of binding sites at a given location. D) The top 15 ranked predictions are then used to calculate the change in Translation Initiation Rate due to binding.

Given this established prior framework we hypothesized that we could craft a model to capture RsmA binding and regulation of genes in *P. aeruginosa*. Homologs of CsrA are found widely across the γ-proteobacteria [32,33]. Within the *Pseudomonas* genus, homologs such as RsmA and RsmE share high sequence and structural similarity with CsrA [34]; the protein sequence of *P. aeruginosa* RsmA is 85% identical to its ortholog CsrA in *E coli* [35]. Furthermore, similar binding mechanisms. SELEX studies have also shown that the RsmA protein shares high affinity for the same binding motif ANGGA [36], and NMR structural studies in the P. fluorescens homolog RsmE also recapitulated affinity for this core motif [34]. In addition, the crystal structures of Csr/Rsm family proteins in complex with RNA are available in the Protein Data Bank for *Escherichia coli* [1Y00], *Yersinia enterocolitica* [2BTI]*, Pseudomonas protegens pf-5* [2MFO], *Pseudomonas fluorescens* [2JPP], and *Pseudomonas aeruginosa* [7YR7]. In tandem with models that leverage data from crystal structures [37] these data can be used to computationally predict changes in free energy for a given motif.

Here, we modify, tune, validate, and improve upon a prior model constructed for the *E. coli* CsrA protein [30] to accurately predict breadth of binding and regulation by the RsmA protein across the entire *Pseudomonas aeruginosa PA14* transcriptome. This approach allows us to probe the entire sequence space computationally, thus lifting the constraints presented by prior experimental approaches. In an improvement upon our prior model, we consider alternative motifs given the generation of a crystal-structure derived, RsmA-specific, position weight matrix. Unlike GGA motif-based screens, our model also yields predictions regarding the mechanism of binding to a given target including: the approximation of binding strength, diversity of binding peak frequencies, and predicting the effect binding has on translation. We also leverage several publicly available high throughput sequencing datasets to statistically verify the accuracy of our predictions. In doing so, we predict 1043 genes to be bound by RsmA and identify 457 genes with no prior binding evidence. Our pool of filtered predictions is enriched in transcriptional regulators and virulence associated pathways. An important resulting observation of this work is the experimental characterization of two novel transcriptional regulators *rsaL* and *mvaT,* mRNA encoding for factors involved in Quorum Sensing and the Type IV Secretion System, among others. In this work, we use model predictions to confirm binding, binding site pockets, and regulation of these mRNAs *in vitro* and *in vivo*. This characterization both validates the predictive capabilities of the model and expand upon our understanding of RsmA regulation. Overall, our constructed model opens up new avenues for differentiating direct from indirect targets of RsmA and aids in generating hypotheses for the varying regulatory mechanisms governing complex signaling networks in PA.

## Materials and Methods

### Construction of model and definition of energy terms

A free energy model constructed for describing binding by the CsrA protein from *Escherichia coli* was described in [30]. In our current approach, we have modified the model to include the nucleotide contributions of bases other than the core ANGGA. This was also tuned to capture RsmA-mRNA interactions using the structure of the *P. fluorescens* RsmE in complex with *hcnA* (PDB: 2JPP). The thermodynamic model relies upon the sum of energetic contributions of 3 key parameters: 1-the position weight matrix of individual nucleotide contributions to binding (ΔG_site1_ & ΔG_site2_), 2-the change in free energy from the unbound to bound state of the mRNA (ΔG_mRNA_), 3-the distance between binding sites to reflect steric effects of dimer binding (ΔG_cooperativity_) (**Fig. 1a**). Total free energy ΔG_total_ is calculated using the following two-state thermodynamic equation, previously defined in [30]:

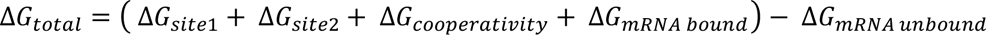

After sorting the summed ΔG_total_ values for each pair of binding sites across the sequence space (**Fig. 1b**) and the position of binding sites for the top 15, highest affinity, predictions were converted into structural constraints within the open source translation rate calculator, OSTIR (**Fig. 1c**) [38]. This yielded a measure of the translation initiation rate for the bound (TIR_RsmA bound i_) and unbound (TIR_unbound_) states for each prediction of binding positions. Effects of binding on translation were calculated as follows:

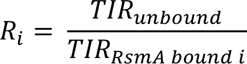

TIR ratios were used to predict the effect that RsmA binding would have on translation, and binned into three categories: repressed (R_i_ > 1.2), activated (R_i_ < 0.8), or no impact (0.8 < R_i_ < 1.2) based on boundaries defined in [30].

### Calculation of the per-nucleotide contributions to binding

The protein sequence of *P. aeruginosa* RsmA is 85% identical to its ortholog CsrA in *E coli*. Key residues for RNA recognition, such as the arginine present at position 44 are conserved. The Rosetta-Vienna RNP ΔΔG tool [37] was used to measure the relative change in binding affinities between a wild-type *hcnA* sequence GGGCUUC**ACGGA**UGAAGCCC (motif in bold) and all possible mutants within the 5-nt binding motif at positions 8-12. The solution NMR structure of *Pseudomonas fluorescens* RsmE in complex with the *hcnA* mRNA [39](PDB: 2JPP) was used as the scaffold of the model. This approach incorporates the RNAfold command within the Vienna RNA package 2.0 [40] to calculate the minimum free energy of each unbound mutant

(**Supplementary table 2**). The position weight matrix of per-nucleotide contributions to binding was calculated as follows:

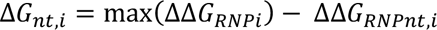

Wherein *i* is the position of the nucleotide within the 5 nt binding motif and *nt* is the specific nucleotide mutation (ATGU) at that position. To generate an energetic measure of the individual nucleotide contribution, each ΔG value was subtracted from the maximum affinity found across all 4 nucleotides at a given position. The ΔG_nt,i_ was then converted from kcal/mol to RT units given the gas constant at 37° C (R = 0.616).

### Generation and modeling of UTR sequences from the PA14 genome

The 5’ Untranslated Region (UTR) of an mRNA transcript is the primary region where the Csr/Rsm family proteins enact their regulatory function by influencing ribosome binding. We selected the 5’ UTR plus the first 100 bases of coding sequence (CDS) to generate predictions via modeling. Prior RNA sequencing in [41] defined the transcription start sites (TSS) across the *P. aeruginosa* PA14 transcriptome at 28° C and 37° C. Where the primary TSS was defined, we selected nucleotides from the TSS site to 100 bases into the CDS. If no TSS was known, we selected -100 bases from the start site to encompass the RBS region. Sequences were extracted from the *Pseudomonas aeruginosa UCBPP-PA14* reference genome assembly GCF_000014625.1. This yielded 5285 UTR sequences which are summarized in **Supplementary table 2**. Predictions of all combinations of 2 binding sites were performed for each of the modeled 4861 sequences in parallel on the Stampede2 compute cluster at the Texas Advanced Computing Center (TACC) at The University of Texas at Austin. Associated python scripts used to run the model on the Stampede2 compute cluster can be found at https://github.com/ajlukasiewicz/rsm_biophysical_model

### Ensemble analysis of predicted binding sites and peak calling

All possible combinations of binding pairs are evaluated across the entire sequence space, and sorted by affinity. This yields an ensemble of predictions per gene with varying degrees of free energies. We then transform the overall affinity score ΔG_total_ into a measure of the likelihood of binding via the Boltzmann probability distribution:

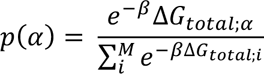

Wherein the probability of a particular binding conformation (p(α)) is a function of the ΔG_total_ for an individual prediction given the distribution of all possible conformations for a gene. β (0.45) is a scaling factor based on thermodynamic predictions of RNA-RNA interactions [30]. Here we alter the scaling factor for calculating this probability using predicted energy and affinity values from our prior model [30] and affinities derived from literature. Measured binding affinities were converted into free energy using the following equation: Δ(G) = RT ln(kD) wherein the gas constant RT at 37° C (-0.616). Dissociation constants were found via prior EMSA experiments for CsrA binding to *glgC, nhaR, cstA, pgaA,* and *rpoE* [42–46]. The Bolzmann probability was used to weigh predicted ΔG_total_ affinity scores in calculating an overall average. We selected a range of β values from 0.35 to 0.45. β= 0.4 was determined to generate the highest linear correlation between the predicted ΔG value and the measured affinity (adjusted R2 = 0.98, p-value = 0.0009527). Linear regression tests were performed in R.

Out of all predictions per gene, the 300 top predictions were used to calculate the Bolzmann probability given the inflection point of energy predictions observed per gene in [30]. The frequency of binding site position predictions was calculated as a function of the Bolzmann probability of binding to that position. These frequencies were then used to calculate densities of binding interactions across the UTR itself, yielding peaks which we interpret as footprints or binding sites of RsmA. Using the *lolB* sequence (**Fig. 2b; supplementary table 3**) as a negative control, we established the peak height threshold for binding to be the maximum height for *lolB* binding site frequencies, 0.0064.

**Figure 2:**
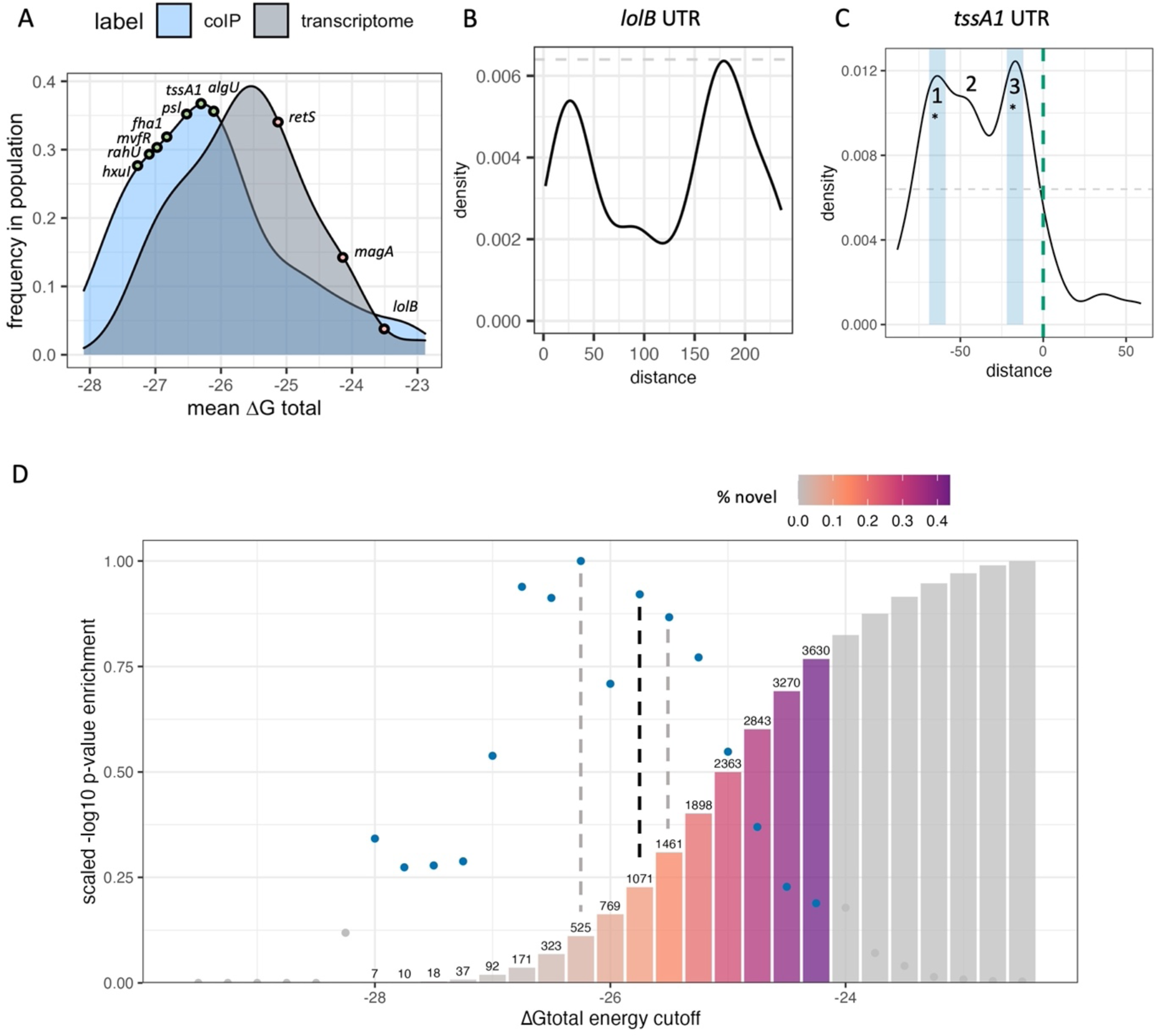
Model parameters are refined and validated using experimental datasets. A) Overall affinity scores from genes identified to be bound by RsmA in prior RNA immunoprecipitation studies (coIP) are a distinct population relative to a random sample from the rest of the transcriptome. Positive and negative control RNA tssA1 and lolB fall at opposite sites in these populations wherein more negative ΔGtotal values represent higher affinity scores. B) Frequency of binding site predictions along the lolB mRNA sequence. Predictions along the sequence space of this gene are very disperse and have low affinity. C) Frequency of binding site predictions across the tssA1 UTR sequence. Binding site frequencies across the space of this sequence pass our threshold at three main sites. *Two of which were confirmed binding sites of RsmA given past mutational studies (Schulmeyer et. al., 2016). D) Hypergeometric enrichment testing reveals that the peak energy cutoff that enriches for known targets of RsmA, while excluding non-targets, is -25.75 kcal/mol (black line). The number of predictions that pass this filter are shown in text, and the % of novel predictions are shown in color. Non colored bars and points represent energy thresholds where predicted targets were not significantly enriched (p-value > 0.05) relative to random chance.

Peaks in binding site density data were called using the signal function within SciPy 1.0 [10] with the following parameters: the peak width was set from 5 to 15 to represent the range between the minimum base pairing footprint and the maximum number of possible predictions for a single site. The minimum height for a peak was set at 0.0064, which was determined to be the maximum height for a negative control UTR, *lolB*. The script for parsing and calling peaks can be found in the rsm_biophysical_model GitHub repository as peak_calling.py. Analysis and generation of footprint density plots was performed in R (Version 4.3.1).

### RNA Co-immunoprecipitation

Strain PA14Δ*rsmAF* carrying an empty vector control (pJN105), pRsmA_His6_, pRsmF_His6_, or the RNA binding mutant expressing plasmids pRsmA(R44A)_His6_ and pRsmF(R62A)_His6_ were grown at 37C with shaking at 300 RPM in 200 ml Tryptic Soy Broth (TSB) supplemented with 20 mM MgCl_2_, 5 mM EGTA, 15 μg/ml gentamicin, and 0.1% arabinose to mid-log phase, and pelleted at 4C. Cells pellets were immediately resuspended and lysed in Qiagen native purification lysis buffer (50 mM NaH_2_PO_4_, 300 mM NaCl, 10 mM imidazole, pH 8.0) supplemented with 2.5 mM vanadyl ribonucleoside complex (NEB) (to inhibit RNase activity), 1 mg/ml lysozyme, and 0.1% Triton X-100. Lysis was completed by three freeze-thaw cycles. Lysates were treated with 10 μl RQ-1 RNase-free DNase and cleared by centrifugation. An aliquot was removed from the cleared lysate for total RNA isolation and preserved in TRIzol (Thermo Fisher), and the remaining lysate was incubated with nickel-nitrilotriacetic acid (Ni-NTA)–agarose at 4°C for 1 h under nondenaturing binding conditions. Ni-NTA–agarose was then loaded into a column and washed 3 times with nondenaturing binding buffer containing 10 mM imidazole. Protein and associated RNAs were eluted in 4 fractions with 250 mM imidazole and 4 fractions with 500 mM imidazole. An aliquot of each fraction was analyzed by western blot, and fractions containing RsmA_His6_, RsmF_His6_ or the respective RNA binding mutant version of the proteins were individually pooled as were the equivalent fractions from the vector control strain. Each pool was treated with TRIzol and RNA was extracted according to the manufacturer’s protocol. RNA was treated with RQ1 RNase-free DNase and concentrated using RNA Clean and Concentrator kit (Zymo).

### Library preparation and Next-Generation Sequencing Analysis

Purified total RNA and co-IP enriched RNA was treated with Ribo-Zero (Illumina) according to the manufacture and purified and concentrated with Zymo Clean and Concentrator 5. First strand cDNA was generated using Superscript II RT (Invitrogen) and Random Primer 9 (NEB) and converted to double stranded cDNA using Second Strand cDNA Synthesis Kit (NEB) according to the manufacturer’s protocols. cDNA was purified using Zymo RNA Clean and Concentrator Kit modified for cDNA recovery. Libraries were prepared using the Nextera XT DNA Library Kit (Illumina, San Diego, CA) according to the manufacture’s protocol including tagment of cDNA, amplicons indexation/barcoding through PCR amplification using Nextera master mix, clean-up, and pooling. Finally, pooled and barcoded amplicons were single end sequenced on an Illumina NextSeq500 System. Sequencing reads were trimmed using Trimmomatic to remove library adapters. Trimmed reads were aligned to a Pseudomonas aeruginosa PA14 reference genome using bowtie2 [47]. Aligned reads were then transformed into binary alignment maps (BAM files) using samtools [48]. Finally, files were analyzed in Geneious software to obtain count tables containing transcripts per million read counts for each gene. Raw sequencing outputs were uploaded to the publicly available Sequence Read Archive (SRA) under the Bioproject ID PRJNA1131461.

Analysis of gene expression was performed using the DEseq2 package [49] in R. To determine enriched genes, we first calculated the differential expression between the total RNA and the overexpressed RsmA-his pulldown genes. Genes with L2FC >1 and p adj < 0.005 were considered enriched in our dataset (**Supplementary table 6**).

### Proteomic sample preparation and analysis

Overnight cultures of WT *P. aeruginosa* PA103 and Δ*rsmA*, Δ*rsmF*, and Δ*rsmAF* mutants were diluted to an optical density of 0.1 at 600 nm (OD_600_) in tryptic soy broth supplemented with 1% glycerol, 100 mM monosodium glutamate, and 2 mM EGTA. Cultures were incubated at 37°C with shaking until the OD_600_ reached 1.0. Cells (1 ml) were harvested by centrifugation (10 min, 4°C, 12,500 x g). Cell pellets were washed with 1 ml PBS and then stored at -80°C. Proteomic sample preparation and analyses were performed by the VIB Proteomics Core, Gent, Belgium. Differentially expressed proteins were identified using the DEseq2 package [49] in R. Proteins with L2FC >1 and p adj < 0.005 were considered differentially expressed in our dataset (**Supplementary table 7**).

### Filter binding assay for testing binding interactions in vitro

Assessment of binding interactions between RsmA and several candidate genes were evaluated using an *in vitro* nitrocellulose filter binding assay. Sequences generated with efficient T7 promoter design and synthesized (IDT). Sequences for these targets can be found in **Supplementary table 3**. RNA was produced via *in vitro* transcription (Thermo T7 megascript kit) with supplemented 3.75 mM guanosine for efficient radiolabeling. P^32^ labeled ATP was integrated to the 5’ end of purified RNA with PNK and cleaned up using silica filter spin column extraction (NEB Monarch).

His-tagged RsmA was purified using nickel chromatography. Briefly, BL21 *E. coli* cells were transformed with an arabinose-inducible, his-tagged RsmA encoding plasmid. These were grown in overnight cultures and seeded into large shaker flasks until reaching exponential phase (OD600 = 0.6).

Binding strengths between purified RsmA and various radiolabeled RNA sequences were assessed using nitrocellulose filter binding. Serially diluted RsmA was incubated with 0.5 nM p32 radiolabeled RNA in an optimized binding buffer (10 mM Tris-HCl pH 7.5, 100 mM KCl, 10 mM MgCl2, 10 mM DTT, 10 ug/mL heparin, Murine RNase inhibitor) at 37 C for 30 minutes. Following incubation, reactions were loaded into the Bio-Dot microfiltration apparatus (Bio-Rad) and light suction was applied to pass the reactions through sandwiched 0.45 mM nitrocellulose and N+ (Cytiva Amersham™ Hybond™-N+) membranes. Signal intensities were captured via phosphorimaging on the Amersham Typhoon 5, and measured using Bio-Rad Image Lab software. Dissociation constants were calculated using the modified hill equation described in [50] with a Hill Constant of 2 to reflect cooperative binding of the homodimeric form of the RsmA protein.

### Construction of translational reporters for assessing effects on regulation in PA10

The effects of RsmA binding on translation were assayed using a translational GFP reporter system. The *E. coli* and *P. aeruginosa* compatible plasmid, pJN105, encodes for a arabinose inducible RsmA expression and was modified as follows: The constitutive lacUV5 promoter upstream of the 5’ UTR of our gene of interest was inserted into pJN105 along with the first 99 bases of coding sequence. This leader was fused to the GFPmut3 sequence with a trailing SRA degradation tag (M0051, sequence from IGEM database). Sequences for our genes of interest, along with positive and negative controls were amplified with compatible primers and inserted through Gibson assembly (NEB HiFi Gibson assembly kit). pJN105 was encoded with an inducible RsmA region via the pBAD promoter and constitutive araC expression. All plasmids and primers used in this study can be found in **Supplementary table 3**.

Following assembly, plasmids were transformed using heat shock into chemically competent *DH5α E. coli* and plated on 15 ug/mL Gentamycin supplemented (Sigma-Aldrich) LB plates. Plasmids were extracted from overnight cultures using the Zymo zippy miniprep kit and submitted to Plasmidsaurus for sequence confirmation. Following extraction, plasmids were then transformed into chemically competent PA103 ΔRsmA/RsmF strains and plated on LB-agar media supplemented with 80 ug/mL Gentamycin antibiotic. Transformed strains were grown overnight in LB broth supplemented with 80 ug/mL Gentamycin (Sigma) and then seeded into 30 mL of supplemented LB culture at a 1:100 dilution. Upon reaching OD 0.02, cultures were split into two flasks and half were induced with 0.5% L-arabinose. Induced and uninduced cultures were monitored for fluorescence intensity on the Cytation3 plate reader at 484 and 513 excitation and emission wavelengths. Fluorescence and OD600 measurements were taken at 0, 1, 2, 4, and 6 hours post induction. Fluorescence values were normalized by OD600 measurement and analyzed in R.

### Generating mutations for rsaL and mvaT

Mutations were made for all combinations of predicted binding sites on *rsaL* and *mvaT* while minimizing the change to overall structure for the folded mRNA. Minimum Free Energy calculations were performed using ViennaRNA RNAfold secondary structure prediction tool (version 2.4.18). All scripts were written and executed in Python 3.7. For binding sites within the coding region, mutations were made to exclude stop codons while still maintaining overall structure. Motif mutations were generated using all combinations of low scoring residues present in our prior PWM. The full list of mutant sequences can be found in **Supplemental table 3**.

## Results

### Using crystallized RsmA-RNA binding structures to generate a biophysical framework that captures different energetic contributions of various RNA sequences to binding

The *P. aeruginosa* RsmA and *E. coli* CsrA protein sequences share 85% amino acid identity (BLAST alignment: Camacho et al., 2009), however slight differences in the primary and secondary binding motifs have been reported for the Csr/Rsm family across organisms [20,51]. To construct an energetic matrix that captures interactions between RsmA and specific motifs in *P. aeruginosa*, we selected the scaffold structure of RsmE-*hcnA* available in the Protein Data Bank (PDB: 2JPP) as representative of the overall protein structure in complex with mRNA. Changes in free energy due to single positional mutations were captured using the Rosetta-Vienna RNP *ΔΔG to*ol [37] as described in (Methods). This generated a Position Weight Matrix (PWM) of per-nucleotide contributions of binding based on their position within a 5-nucleotide window (Table 1).

**Table 1:** Rosetta modeling derived Position Weight Matrix of the free energy contributions for each nucleotide present in a 5 nt window. An example of this calculation would be as follows: high affinity motifs such as AUGGA would contribute the maximum possible score to the overall free energy calculation, whereas low affinity sequences such as UCCUU would not contribute to the overall score at all.

| pos/nt | A | G | C | U |
| --- | --- | --- | --- | --- |
| 1 | -1.97 | -0.49 | -0.27 | 0 |
| 2 | -0.02 | -0.04 | 0 | -0.16 |
| 3 | -2.18 | -3.42 | 0 | -0.09 |
| 4 | -1.76 | -5.10 | -0.54 | 0 |
| 5 | -2.11 | -1.90 | -0.08 | 0 |

The highest affinity motif produced by a 5-nt window using this crafted PWM (Table 1) would therefore be AUGGA, which is consistent with the binding motif observed for RsmA [34]. Prior models crafted for the *E. coli* CsrA protein confer the highest energetic contribution when a strict AAGGA motif is found [30]. A comparison of the two matrices can be found in Supplemental table 1. The Rosetta-crafted PWM presented here confers an additional benefit to the model, wherein non-canonical motifs may contribute to the overall energy calculation and thus considers alternative sequences that RsmA can bind. Using this PWM we can then calculate the free energy contributions of a motif within sliding 5 nt windows (ΔG_site1_ and ΔG_site2_), which we sum with additional biophysical parameters (Equation 1) to generate a prediction of overall affinity, or the change in free energy (ΔG_total_) due to RsmA binding to an mRNA of interest.

To briefly summarize the contributions of this PWM to our two-step thermodynamic equation, we calculate the ΔG_total_ as the change in free energy from the unbound (ΔG_mRNA unbound_) to a bound state. These biophysical parameters are defined as follows: The energies of the bound state are calculated given the matrix-derived free energy of each motif bound by the homodimeric form of RsmA (ΔG_site1_ and ΔG_site2_) and added to a penalty for steric hindrance for binding sites in close proximity (ΔG_cooperativity_) and the minimum free energy of RNA folding given bound folding constraints (ΔG_mRNA bound_)(**Fig. 1a**, **Equation 1**). These calculations are performed for all possible combinations of binding sites along each transcript modeled (**Fig. 1b**) and the positions are sorted by the predicted highest affinity. Given the empirically-derived nature of these energy terms, we hypothesize that the *in-silico* predictions of high energetic affinity (ΔG_total_) can be used to predict binding interactions *in-vivo*.

### Genes enriched in RNA co-immunoprecipitation and proteomics establish positive control population for model tuning

To tune model filtering terms, we established a positive control population using RNA co-immunoprecipitation sequencing (RIP-seq) and proteomics. For the RIP-seq experiments Total RNA and pulled down fractions were sequenced in PA14 Δ*rsmAF* carrying plasmids encoding His-tagged RsmA, RsmF, the respective inactive mutants (RsmA R44A, RsmF R62A) or an empty vector control (pJN105). PCA analysis (**Supplementary** Figure 2a) of RNA sequencing performed for the pulldown study suggests that the difference in RNA in total and enriched fractions contributed to 33% of the observed variance in the dataset. 18% of the variance could be attributed to an inactivating mutation present in the overexpressed RsmF protein. Conditions lacking vector expressing RsmA/RsmF and the presence of empty vector encoding no protein both clustered closely and therefore the presence of the plasmid did not alter gene expression.

358 genes were identified to be significantly enriched (L2FC >1 and p-adj < 0.005; **Supplementary** Figure 3b**, Supplementary Table 6**) in RsmA pulldown relative to the total RNA. These targets were considered to have a high likelihood of being bound partners of RsmA and were used to define the positive control population to tune the cutoff term for our model. This enriched population included positive controls such as *algU*, *rahU,* and *magA,* however, other well characterized direct targets of RsmA (“positive control genes”) such as *tssA1* were not enriched in the RsmA pulldown pool. Interestingly, more genes were significantly enriched in the RsmF pulldown relative RsmA (**Supplementary** Figure 3b). This pool of 565 mRNAs included positive control genes such as *tssA1, fha1, rahU,* and *mucA.* 228/565 genes overlap with the pool of enriched mRNAs pulled down by RsmA.

The proteomics experiments identified an additional 261 proteins (**Supplementary Table 7**) found to be significantly differentially expressed (L2FC >1, p-adj < 0.005) in PA103 ΔrsmA strain relative to WT (interpreted as repressed in native conditions).

### Predicted total affinity can be used to differentiate bound from unbound targets

The predicted overall affinity score, ΔG_total_, can be interpreted as a probability for binding occurring when RsmA and the target mRNA are present. To evaluate the predictive capabilities of the model, we sought to determine whether the calculated total affinity score could be used as a metric to differentiate direct binding interactions from indirect or unbound gene targets. Predictions were generated for 5861 UTR sequences extracted from the PA14-UCBB transcriptome (NCBI:txid 208963, **Supplementary table 4**). As of this publication, PA14 has a total of 5893 identified genes but we were unable to generate predictions for all due to their lack of inclusion in prior TSS profiling [41]. To evaluate our predictions, we sought to compare the model predictions to experimental results. A combination of prior RNA co-immunoprecipitation sequencing [9] and the RNA co-immunoprecipitation and proteomics performed in this work were used to experimentally identify 780 genes potentially regulated by RsmA. This pool of genes was used to define a positive control population for binding. A random selection of 780 additional UTR sequences were collected from the rest of the modeled PA14 transcriptome to generate a control population. For each gene within the positive and background populations, the average ΔG_total_ affinity score was calculated given the 300 most favorable predicted energies in the ensemble. These first 300 predictions represent the most probable conformations of binding between RsmA and the RNA target. A significant difference (p <0.05) was observed between the average total affinity scores of 780 randomly selected sequences and those from Co-IP enriched genes (**Fig. 2a**). We identified several control genes to validate our results.The *tssA1* (positive) and *lolB,* (negative) genes are outlined (**Fig. 2a**) due to their extensive binding characterization. These fall at expected values within each population. The average total affinity score for *tssA1* was determined to be highly favorable (ΔG_total_: -27.75 RT), and fell within the energy range for our positive control population (**Fig. 2a**). The average total affinity for the negative control, *lolB,* was calculated to be -23.80 RT which fell within the population range for our randomly selected “non-targets” population. This indicated to us that we could use the ΔG_total_ metric as a cutoff for filtering true from false targets in our pool of predictions.

To further refine the exact ΔG_total_ cutoff that differentiates direct bound targets from indirect non-targets, we performed hypergeometric enrichment testing for the pool of predictions that would enrich for genes pulled down in prior RIP-seq studies, while also minimizing those included by random chance. We evaluated cutoff values within a ΔG_total_ range of -27.50 RT to -24.0 RT (**Fig. 2d**). The cutoff value that conferred the highest significant enrichment for immunoprecipitated genes was found at a ΔG_total_ threshold of -26.25 (p = 7.08e-08), and the second highest at ΔG_total_ -25.75 (p = 2.61e-07). In addition, genes with no prior evidence of binding by RsmA were selected to performed exclusion testing of non-targets for each energy cutoff. This determined that the depletion of non-targets reached its maximum at the cutoff value of -25.50 (p = 6.36e-07). Given these results, the optimal cutoff used was -25.75 which yielded 1071 predictions of putative targets for RsmA. This observation validated that the ΔG_total_ can be used as a predictor of overall affinity.

### Peak analysis of predicted binding sites for enriched targets validate the predictive capabilities of the model

In addition to predicting an overall affinity, our model also has the capability to determine the position of RsmA binding sites along the modeled mRNA leader sequence. The Boltzmann probability of binding was calculated given the ΔG_total_ per prediction presented in Equation 4, and described in Methods. Calculation of the frequency of binding interactions at a specific site was extrapolated from this predetermined probability and used to weigh highest affinity predictions relative to the expanded set of those per gene. Then, peak calling was performed on all genes with a baseline cutoff established from the negative control sequence of the *lolB* mRNA leader sequence (Methods). The application of this cutoff filtered our list of predictions to 1043 possible targets of RsmA, 457 of which are genes for which no prior experimental evidence was found.

The specific binding sites of *P. aeruginosa* RsmA on its established targetome has been experimentally validated on *tssA1*[36]. To evaluate the capabilities of the model for predicting bound regions, we compared peak predictions on the 5’ UTR of *tssA1* which has been experimentally verified binding sites that fall at -15 and -67 nt from the start codon [36]. Predicted binding site peaks not only fall within those two regions (**Fig. 2b**), but also identify a third region where RsmA may potentially bind to repress translation of *tssA1.* Confirmation of more than two binding sites that confer flexible binding of the protein to a given mRNA target has been identified for CsrA [52]. Due to the lack of footprinting data available for other mRNAs within PA, binding site predictions were also performed on experimentally footprinted targets of Rsm/Csr family proteins in closely related organisms, such as *E. coli* (CsrA-*glgC*) and *P. fluorescens* (RsmE-*hcnA*). These produced high positive predictive values on those binding partners (**Supplementary Fig. 2**). Peak predictions for all modeled genes can be found in the supplementary binding packet. Overall, the capturing multiple experimentally characterized binding site across a range of well-studied RsmA/CsrA targets that we selected provided confidence in the ability of the model to identify RsmA binding sites across different potential mRNA targets.

### Enrichment of quorum sensing and biofilm pathway transcription factors in predicted RsmA targets

Given our pool of 1043 predicted targets, we next sought to determine whether new pathways that were regulated by RsmA (but not yet identified) were enriched in our filtered pool. Encouragingly, pathways with prior experimental evidence of regulation by the GacA/S TCS pathway were identified in our analyses. GO term and KEGG pathway enrichment analyses of our pool of 1043 putative mRNA targets show significant (EASE score < 0.1) representation of genes involved in key virulence pathways (**Fig. 3a,b**). Molecular features enriched in our predicted targets include those with DNA-binding transcriptional activator (GO:0001216), metal ion binding (GO: 0046872) and cytochrome-c oxidase (GO:0004129) activities (**Fig. 3a**). Roughly 60 transcriptional regulators were predicted to be bound by RsmA in our model, including key QS regulators LasR, MvfR, and the orphan regulator, QscR. Key pathways enriched by our predictions include quorum sensing (pae02024), biofilm formation (pae02025), valine, leucine, and isoleucine degradation (pae00280), and peptidoglycan biosynthesis (pae00550)(**Fig. 3b**). Although many of these processes have already been shown to be regulated by the Gac/Rsm pathway [53,54], several novel predictions were generated within each feature (**Fig. 3a,b**). This suggests modeling allows us to expand upon the total number of genes that RsmA may regulate across complex and condition-sensitive pathways.

**Figure 3.**
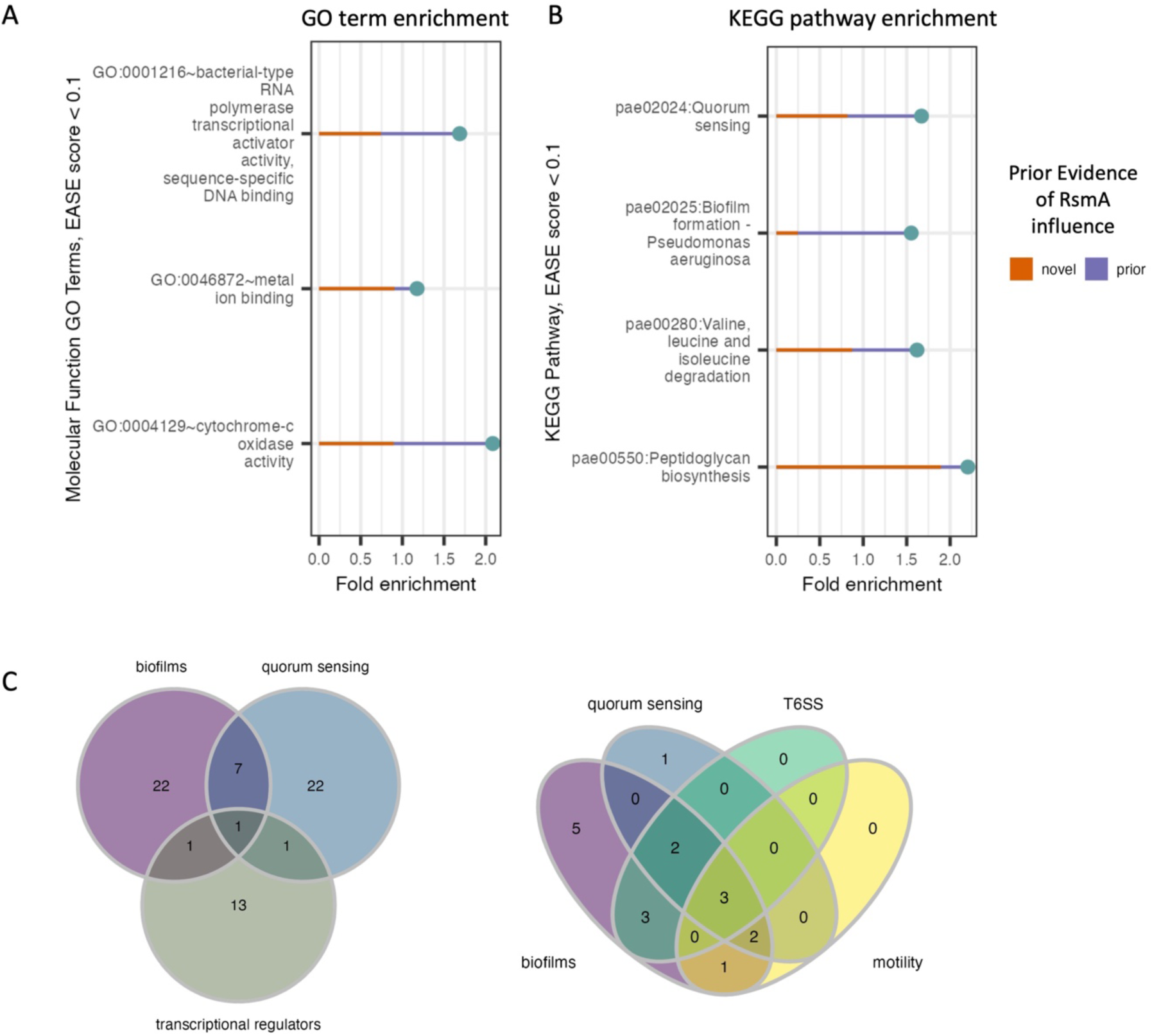
Distribution of enriched molecular functions and pathways in pool of predicted targets of RsmA. A&B) DAVID enrichment analysis for molecular function GO terms and KEGG pathways, sorted by increasing p-value (< 0.1). Along with the reported fold change of enrichment, the lines display the proportion of genes within each category that are novel predictions yielded by the model (red line) and the proportion of genes with some prior evidence of association with RsmA (purple line). C) Predictions that fall within key virulence pathways such as quorum sensing and biofilm formation also have shared transcription factor regulation. Left: one transcriptional regulator, LasR (PA14_45960) is associated with both quorum sensing and biofilms. Right: Model predictions identify several newly profiled transcription factors (Wang, et. al., 2020) that are also associated with virulence pathways. Three transcriptional regulators associated with all four processes include PA1437, a two-component response regulator, PA4184, SouR regulator of Phezanine biosynthesis, and PA1431, rsaL, a novel target and regulator involved in quorum sensing.

The full profiling of transcriptional regulatory network in PA is yet incomplete, but recent efforts to characterize binding specificities *in vitro* [55] has expanded upon our understanding of TF interaction with known, key virulence pathways. Transcriptional regulators were significantly enriched in our predicted pool of genes bound by RsmA (**Fig. 3a**); therefore, we sought to identify which of these transcriptional regulators were associated with KEGG enriched pathways. Of note is the identification of *lasR* (PA14_45960) is shared by both QS and biofilm forming processes (**Fig. 3c**). Out of 86 total transcription factors mapped to biofilm, quorum sensing, the Type 6 Secretion System (T6SS) and motility pathways in [55], 17 were identified by our model to be bound by RsmA. Of these 17, 3 were found to be associated with all four pathways (**Fig. 3c**), which were identified as PA1431 (*rsaL*), PA4184 (*souR*), and PA1437, a two-component response regulator. Only PA1437 was previously predicted to be a potential target via a prior motif search approach [20], whereas PA1431 (*rsaL*) and PA4184 (*souR)* are entirely novel mRNA predictions.

### Meta-analysis of aggregated RNA-seq datasets reveal that novel targets identified in our model are lowly expressed in standard media types used for binding/pulldown studies

The influence of RsmA on regulating the aforementioned pathways has been well demonstrated by prior studies [53,54]. Therefore, we sought to determine how many of our predicted genes were also found in other high-throughput characterizations of RsmA regulation in *P. aeruginosa.* We compared predictions to all those found in previous modeling [20], microarray analysis [18], RNA-seq studies [1,19], RIP-seq studies [9], CLIP-seq studies [10], and recent nascent chain profling methods such as ChiPPar-seq [19]. Comparisons across these studies revealed that 586 of our predictions had some level of prior evidence of binding or direct/indirect regulation by RsmA, and 457 were entirely novel predictions.

Prior experimental approaches have estimated RsmA has some regulatory effect (including direct and indirect) on approximately 500 genes, yet our number of predictions (1043) is double that estimate. In an effort to understand why our pool of predictions is larger than prior approximations, we hypothesized that many predictions were dependent on conditions not tested in prior experimental screens. To investigate this hypothesis we leveraged the aggregated, publicly available, RNA sequencing data from a meta-analysis of gene expression across various conditions in *P. aeruginosa* [23]. This dataset included values of normalized gene expression in transcripts per million (log TPM) from 411 sequencing datasets, including data from a RsmA pulldown study [9]. These datasets measure gene expression in a wide variety of experimental conditions including various strain types, growth phases, media, antibiotic supplementation, clinical isolates, and lifestyles and demonstrates that gene expression is highly variable and condition-specific [23]. In our analysis, we interpreted a gene to be expressed if the log TPM value was greater than 0. The expression data was filtered and subsequently binned into 10 ranges and then labeled given their prior evidence for regulation by RsmA. Overall, genes with some prior experimental evidence of binding to RsmA were more represented in higher expression bins, whereas those that had no evidence, or were novel predictions by our model, aggregated towards lower expression bins (**Fig. 4a**). This observation suggests that the novel predictions generated by the model were not identified as RsmA targets in prior experimental screens due to low expression levels in the conditions tested.

**Figure 4:**
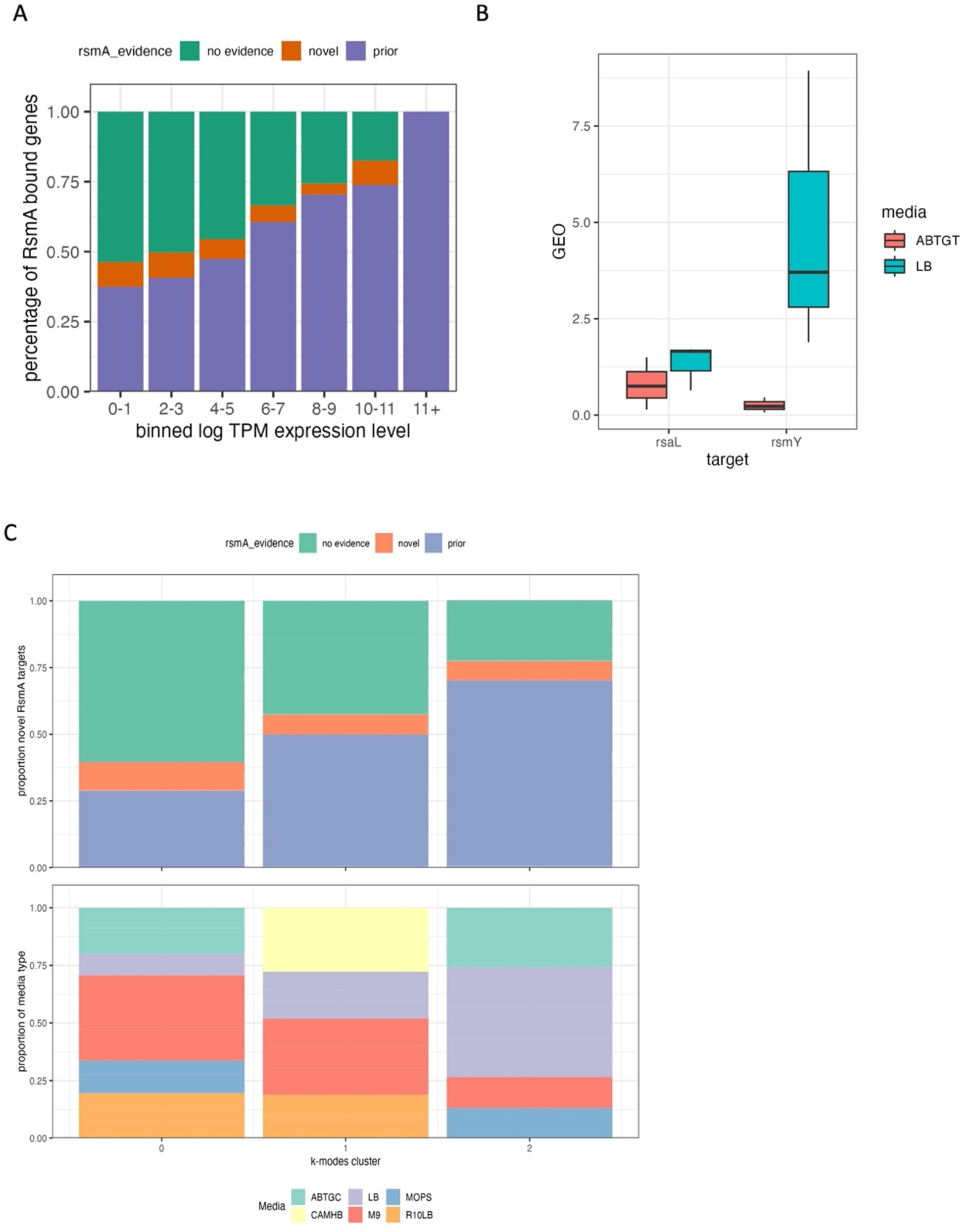
Comparison of results with transcriptomic data suggests novel transcripts are found at lower concentrations. 483 genes were predicted by the model that are not represented in any prior modeling, microarray, RNA-seq or pulldown studies of RsmA. A recent publication aggregated 411 expression datasets for Pseudomonas aeruginosa grown in various experimental conditions. A) Bar chart of the proportion of predictions with no evidence of RNA binding, prior evidence of binding, and entirely novel predictions, binned by log TPM expression level in that experiment. The proportion of genes with prior evidence increases as the log TPM levels of expression increases, suggesting that expression influences detection. B) RT-qPCR data of rsaL and RsmY expression in minimal and LB media at early stationary phase suggests that rsaL is lowly expressed in both media types. C) K-modal clustering of all categories in the aggregated experimental conditions from 411 expression datasets (KO, media, growth phase, stress) to observe whether the presence of novel or predicted targets cluster within specific conditions, knockouts, or media types, overlaid with the proportion of genes that fall within each cluster. Predictably, most of the genes that have some prior association with RsmA are expressed in conditions cultured in LB media, whereas more novel targets were expressed in minimal media such as M9 or ABTHC.

To assess where novel predictions were clustering across these varied conditions, we used k-modal clustering of experimental condition categories as described in Methods. Overall, a higher proportion of genes with some prior evidence of RsmA interaction were found in experiments performed in LB media (cluster 3), whereas nutrient-limited media types like M9 and ABTGT exhibited a higher proportion of novel predictions and genes with no RsmA regulatory evidence (**Fig. 4c, cluster 1**). This recapitulates observations that media type has a large impact on gene expression, and therefore the availability of certain genes for high throughput profiling. As example of note is *rsaL*, a novel target encoding for a quorum sensing transcriptional regulator, that we identify to be bound by RsmA computationally but, when assessed across datasets, appears rarely expressed. We define high expression in this case as a log TPM value greater than that of the *rimM* housekeeping gene (average log TPM = 1.95). *RsaL* reaches a log TPM expression level above 1.95 in only 3 of the 411 RNA-seq experiments (SRA accession numbers SRS605141, SRR6018047, and ERS530377) aggregated in [23]; indicating that sufficient levels of *rsaL* expression may only occur in certain experimental conditions.

To assess whether expression of *rsaL* could be detected if media and growth conditions were optimized, we evaluated expression levels of the gene via RT-qPCR. To mimic the planktonic conditions where *rsaL* expression was detected [56], we cultured PA103 WT strains in either minimal ABTGT or LB media and sampled for *rsaL* expression at late-exponential phase. After normalization to the *rimM* housekeeping gene and to internal primer efficiency E scores, expression of *rsaL* was not significantly different between media types (**Fig. 4b**). In addition, expression of RsmY was significantly increased in LB media relative to ABTGT, but showed no significant change in ABTGT relative to *rsaL*

### *RsmA binds and regulates several predicted mRNA targets* encoding for key transcriptional regulators as assessed by *in vitro* binding and *in vivo* translational reporter assays

Given the concordance of our computational predictions with previously published experimental results, we sought to test RsmA binding to our novel predictions *in vitro*. Therefore, we selected 8 genes that were representative of the core quorum sensing regulatory cascade (**Fig. 5a**, **Supplementary table 3**) to assess binding *in vitro*. These were quorum sensing regulatory genes *lasR/lasI, rhlR/rhlI, mvfR,* and a novel prediction *rsaL.* Secretion system regulators included the *mvaT* and *aprD* leader sequences. These targets have varied support in the literature for RsmA interactions, the majority lacking evidence of either *in vitro* binding or regulatory impact. Finally, the *tssA1* and *loB* sequences were included as positive and negative controls. Filter binding assays were performed with the [α-^32^P] ATP radiolabeled mRNA and purified RsmA protein. *aprD* binding was evaluated via Electrophoretic Mobility Shift Assay (EMSA) (**Supplemental Fig. 4**). Each of these genes had varying degrees of prior RsmA regulatory characterization as summarized in **Fig. 5a**. Importantly, we observed strong *in vitro* binding interactions between RsmA and *mvaT, lasR, rhlI* and *tssA1* leader sequences. These observations are consistent with the predicted overall affinity (ΔG_total_) scores for each gene, which were predicted to be -26.29, -26.54, -26.37, and -26.34 respectively (**Fig. 5a,b**). Weaker interactions were seen for *rsaL, mvfR,* and *lasI.* These each had average predicted affinities of - 25.82, -26.55, and -24.79 (**Fig. 5a,b**). Disassociation constants (kDs) from this biochemical characterization correlate well with the predicted total affinity (R2= 0.92, **Fig. 5c**). It is worth noting that although we initially excluded genes such as *rhlR* from our true target predictions (in accordance with the -25.75 energy threshold), we tested them experimentally for binding given the observation that we predicted two other mRNA targets *(lasR* and *lasI)* in our final candidate pool that encode for two closely functionally related proteins to RhlR in the quorum sensing pathway. We did not observe binding between RsmA and *rhlR* in our *in vitro* filter binding assays (**Fig. 5b**) or via EMSA (**Supplementary Fig. 4**) experiments, which recapitulates the negative result from the model. Finally, we did not observe binding between RsmA and the *lolB* negative control. Overall, these results indicate that RsmA does bind to targets predicted by the model, and that relative binding affinity predicted via the ΔG_total_ affinity score is correlated with affinities measured *in vitro*.

**Figure 5:**
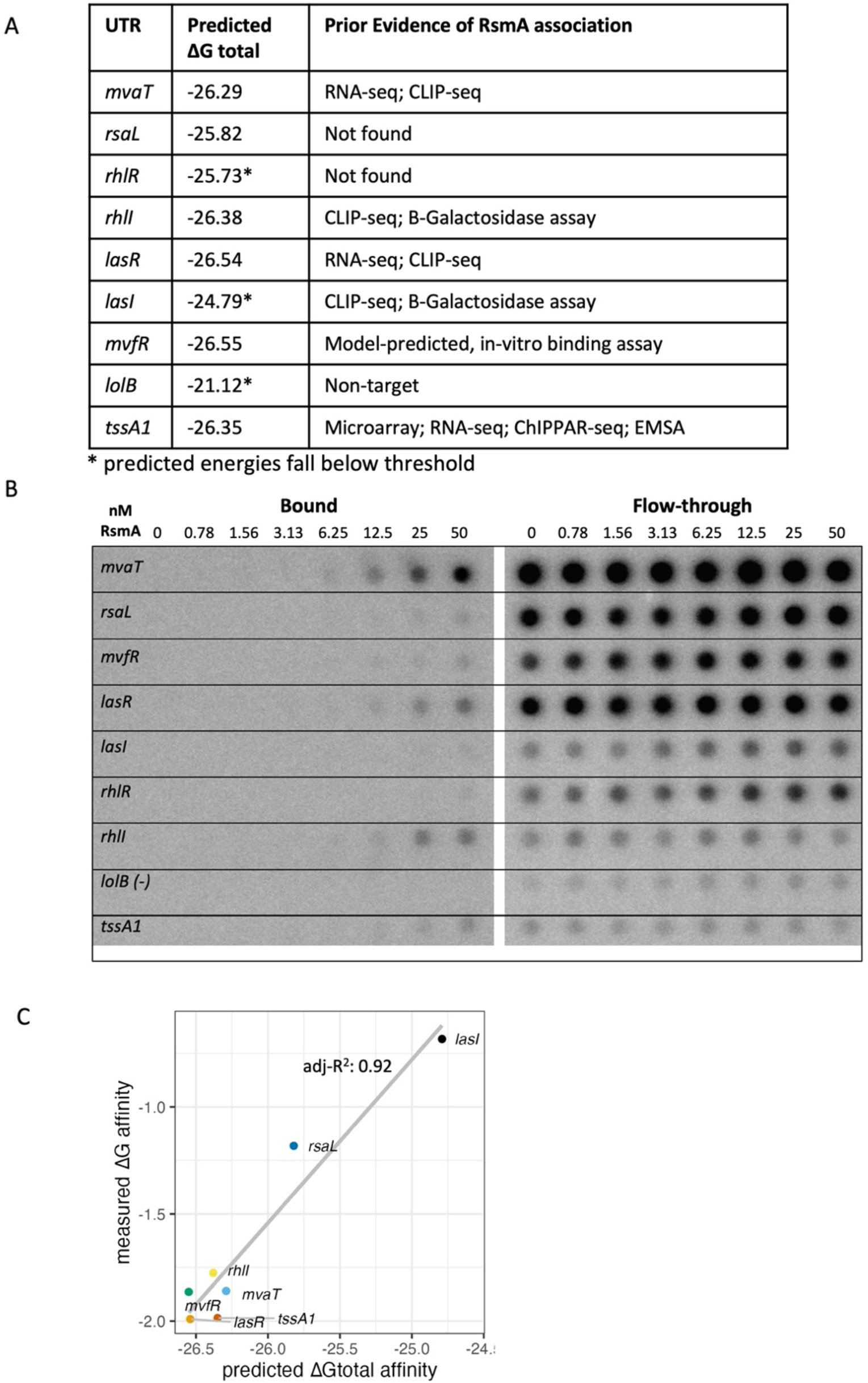
in vitro filter binding assay demonstrates binding interactions between RsmA and predicted targets. A) Summary table of the genes tested for in vitro filter binding which are representative of a variety of predicted energies and prior levels of characterization. B) Phosphoscreen of bound and unbound radiolabeled intensities for the UTRs presented in table A. C) A linear correlation exists between predicted and measured disassociation constants generated from fitting filter binding assay.

As a post-transcriptional regulator, RsmA is able to repress or activate gene expression by blocking or enhancing ribosomal binding to the 5’ UTR region of an mRNA. To evaluate the effects of binding on translation, we performed plasmid-based *in vivo* translational reporter assays (summarized in **Fig. 6a**). Sequences from the same pool of 8 genes selected for *in vitro* characterization were fused to the GFPmut3 coding sequence, and fluorescence values were measured following RsmA induction in a PA103 ΔRsmA/RsmF strain (**Supplementary table 3**). *lolB* was not used in these assays due to the observation that the established sequence used in prior mobility shift experiments [1] is not the leader sequence, but falls within a portion of the coding region and therefore does not contain a ribosome binding site (see supplemental table 3). Specifically, BLAST search revealed the *lolB* sequence used in prior experiments falls between nucleotides 5236896 and 5237178 in the PA01 genome. Given the lack of binding observed between RsmA and *rhlR* in our *in vitro* binding assays (**Fig. 5b, Supplementary Figure 4**) we selected this target to use as a suitable negative control for this assay. No significant difference in fluorescence is observed for *rhlR* (**Fig. 6b**). The *tssA1* 5’ UTR was used as a positive control for repression and showed a significant (p<0.05) reduction in normalized fluorescence values following induction of RsmA (Fig. 4b). We also observed significant reduction of fluorescent signal for the HSL synthetase genes *lasI* and *rhlI* (p < 0.001, and p < 0.05, respectively) (**Fig. 6c**). Given results for our positive and negative regulatory controls, we then performed the assay on *mvaT, lasR* and *rsaL*. Each of these genes have some lacking prior evidence of direct RsmA binding and/or regulation from the literature (**Fig. 5a**). These targets yielded reduced fluorescent values following RsmA induction (**Fig. 6c)** and we interpret these results to suggest these genes are repressed by RsmA *in vivo*.

**Figure 6:**
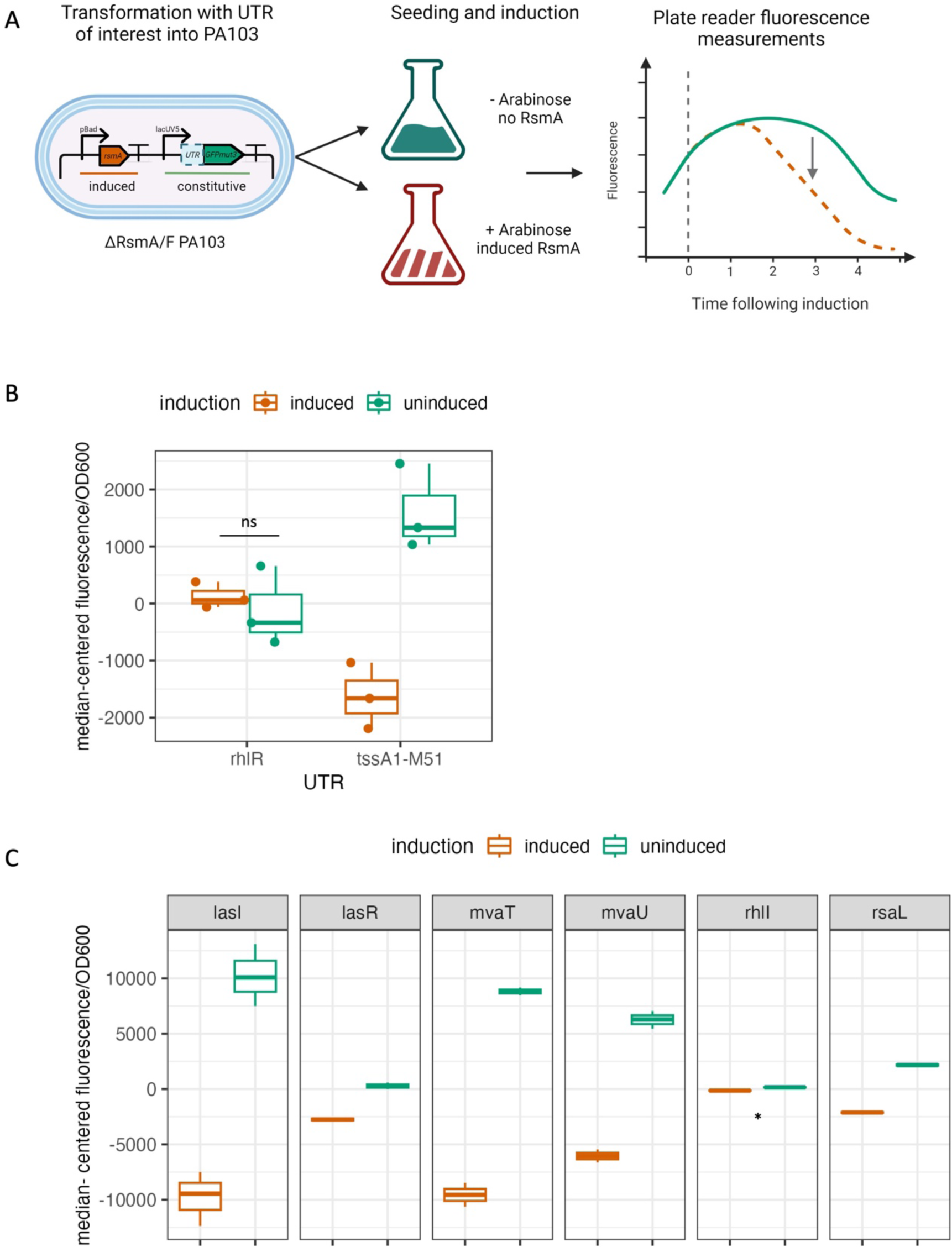
in vivo repression assay. A) Experimental overview of in vivo translational repression assay. UTRs were fused to GFPmut3 and expressed off of the lacUV5 constitutive promoter. Plasmids were transformed into PA103 ΔRsmA/RsmF strains and seeded into +/- 0.5% arabinose LB media. Fluorescence was monitored up to 6 hours following induction. B) rhlR and tssA1 UTR sequences were used as negative and positive controls for our assay. No significant change in fluorescence was measured for rhlR, which is consistent with our prediction and in vitro experimental results. A significant reduction in fluorescence values was observed for the positive control tssA1. C) A significant reduction in fluorescence was also detected for our pool of additional tested genes, including lasI and rhlI. Fluorescence values are plotted median centered to account for changes in translation rates due to the native RBS encoded in each individual UTR.

### RsmA binds to model-predicted binding sites in novel targets rsaL and mvaT in vitro

The model identifies several binding sites along the sequence space of each gene. Given our observation that two novel targets *rsaL* and *mvaT* were bound by RsmA *in vitro,* we sought to assess binding to the specific predicted locations produced by the model. The top three binding sites for each gene (**Fig.s 7a,b and Fig.s 8a,b**) were mutated individually, and for all combinations of 2 binding sites along the sequence. Binding to each mutant was evaluated via *in vitro* filter binding assay.

**Figure 7:**
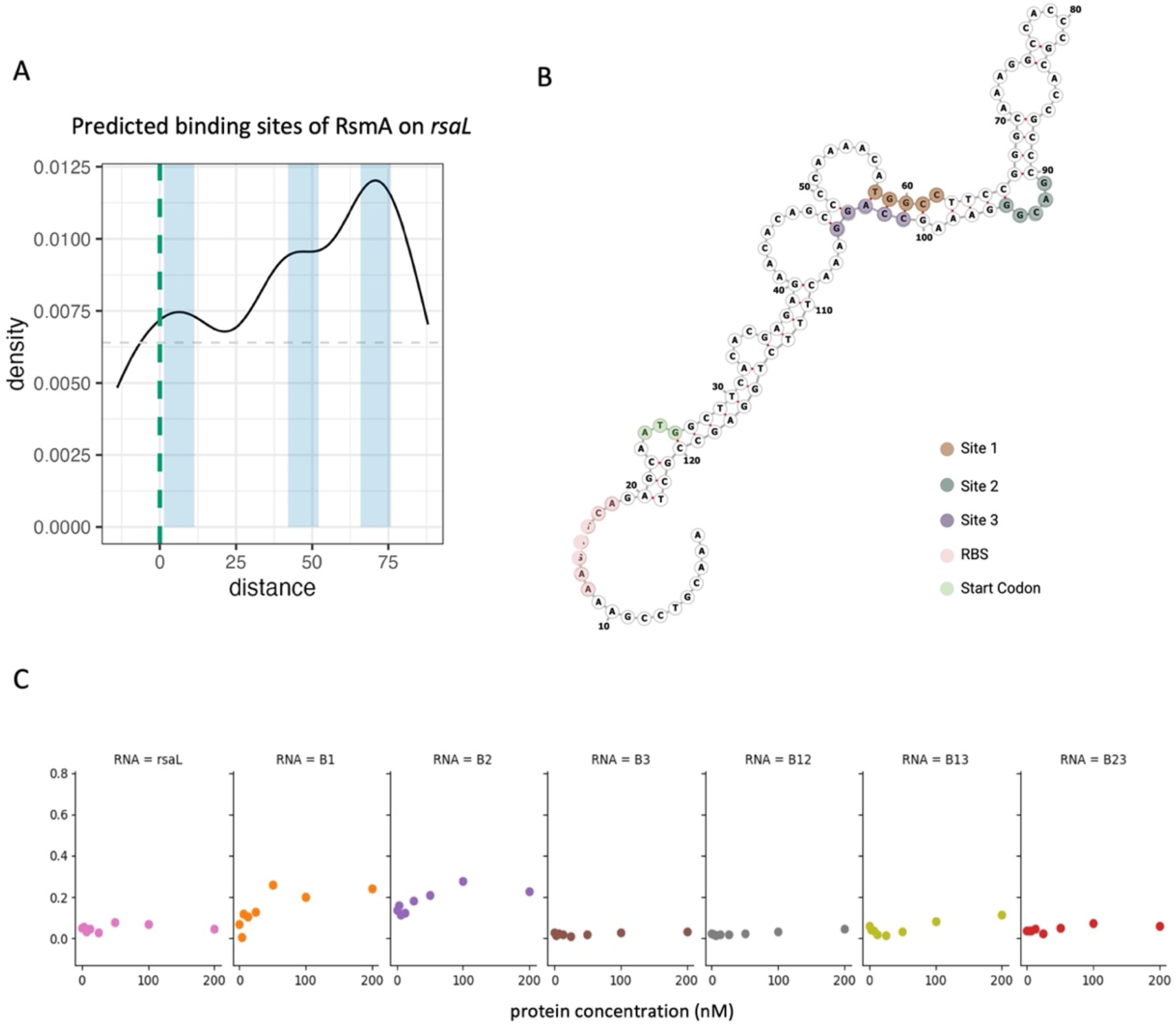
Mutational evaluation of predicted RsmA binding sites on rsaL. A) Density plot of predicted binding pockets along the modeled region of the rsaL leader sequence + 100 bases of CDS. Blue boxes represent the highest frequency regions for the ensemble of predictions along the sequence space. Light grey dashed line represents the minimum peak threshold for considering a binding pocket. Green dashed line is the start codon. B) Structural diagram of the rsaL leader sequence with labeled binding pockets (brown, green, and purple) as well as key functional regions such as the start codon (green) and predicted RBS (pink). C) Filter binding generated binding curves for RsmA in complex with WT rsaL (pink) and individual mutations (orange through brown) or mutations in combination (grey through red).

**Figure 8:**
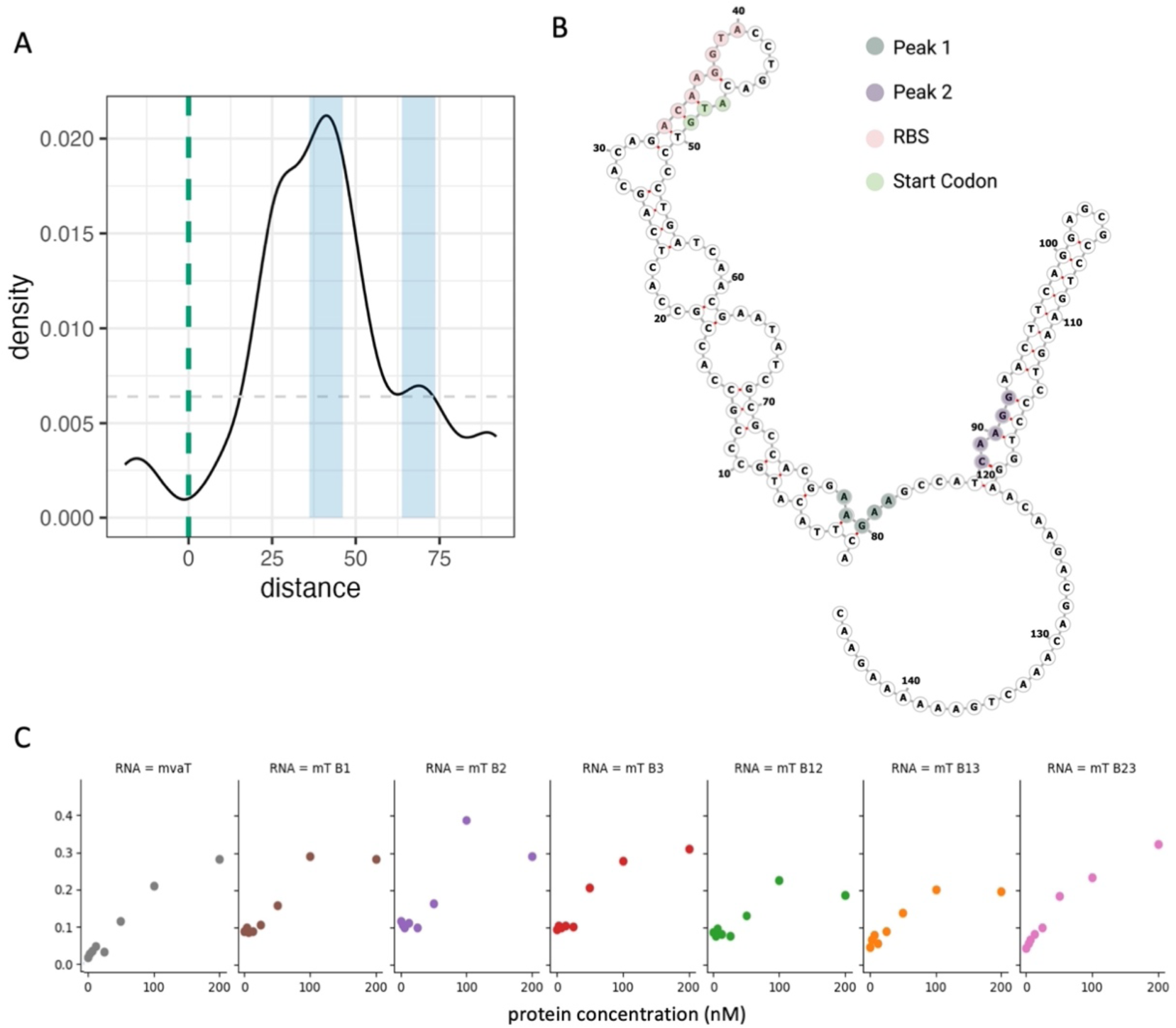
Mutational evaluation of predicted RsmA binding sites on mvaT. A) Density plot of predicted binding pockets along the modeled region of the mvaT leader sequence + 100 bases of CDS. Blue boxes represent the highest frequency regions for the ensemble of predictions along the sequence space. Light grey dashed line represents the minimum peak threshold for considering a binding pocket. Green dashed line is the start codon. B) Structural diagram of the mvaT leader sequence with labeled binding pockets (green, and purple) as well as key functional regions such as the start codon (green) and predicted RBS (pink). C) Filter binding generated binding curves for RsmA in complex with WT mvaT (grey) and individual site mutations (brown through red) or mutations in combination (green through pink).

The three predicted binding sites (termed BS1, BS2, and BS3) on the *rsaL* transcript fall within the coding region at +12, +67, and +76 nt from the start codon (**Fig. 7a,** BS1, BS2, and BS3), with the highest frequency of binding predictions falling peaks 67 and 76 nt (**Fig. 7a**). Guided by the strict peaks (i.e. specific binding sites) predicted by the model in this case, we selected these three specific binding sites to test. Evaluating these mutations via *in vitro* binding reveals that mutation of BS3 significantly reduces binding affinity of RsmA to the *rsaL* transcript (**Fig. 7c**). Mutating BS1 and BS2 individually did not alter affinity to the transcript however, tandem mutations at sites BS1 and BS2 as well as sites BS2 and BS3 hinder binding interactions from occurring. Overall these results suggest that BS3 is the main anchor of binding interactions with the transcript, with BS2 as the site with second highest affinity. Mutation of BS1, which falls below our peak threshold, did not impact binding as strongly and is therefore a less likely site for RsmA-*rsaL* interactions.

Relative to the distinct peaks observed on *rsaL*, binding site predictions on the *mvaT* leader sequence fall in a wider range, as evidenced by a single peak in the within the coding sequence of the gene (**Fig. 8b**). Predicted binding sites on the *mvaT* leader sequence were mutated at positions +26, +41, and +68 nt from the start codon (**Fig. 8a,b**, BS1, BS2, and BS3). Given the lack of distinct peaks, and therefore a broader selection of potential binding sites, RsmA-*mvaT* binding interactions were not disrupted as expected. Specifically, in our *in vitro* binding assays, no change in affinity was observed by mutating BS1, BS2, or BS3 individually. A slight decrease in affinity was observed when mutating BS1 and BS2, or BS1 and BS3 in tandem (**Fig. 8c**). It is interesting to note that predicted RsmA binding sites along the *mvaT* sequence cluster in a wide region within the CDS (**Fig. 8a**), suggesting that there may be a multitude of conformations by which RsmA binds to this transcript.

## Discussion

In this work, we expand beyond motif-based screens to computationally profile binding and regulation by the RsmA protein across the entire *P. aeruginosa* transcriptome. Modeling and subsequent filtering yielded 1043 potential targets, of which 457 were not identified in prior experimental screens. We deem these as novel putative targets of RsmA. These putative novel targets were found to have variable media and condition-specific expression when investigated in context of publicly available sequencing data, which we posit explains earlier inability to detect them. Within each prediction we identify key molecular features that influence binding, and used these to effectively differentiate direct from indirect binding. Overall, this effort demonstrates the utility in using empirically derived binding parameters to computationally interrogate expansive sequence spaces.

### Metrics such as the energy terms and binding sites correlate with experimental evidence, which demonstrate utility of model in predicting true vs false targets of RsmA

Given empirically derived binding parameters, our free energy model of RsmA binding was able to differentiate direct from indirect or unbound targets. Our predictions of overall affinity (ΔG_total_) and the position of binding sites were identified as the key parameters that allowed us to interrogate binding to mRNA leader sequences across the transcriptome. Molecular features on the RNA sequence are key for enabling regulatory function, and also provide information on the mechanism by which RsmA is able to bind. In comparing our model predictions to publicly available pulldown sequencing data, we demonstrate that the calculation of the overall affinity term ΔG_total_ can be used as a metric to differentiate true from false targets of RsmA (**Fig. 2**) which allowed us to effectively filter predictions made across the entire transcriptome. This was facilitated by improvements made to tailor our model for the *P. aeruginosa* RsmA protein. One such improvement was the generation of a RsmA-specific PWM (**Table 1**). This PWM allows for the contribution of non-canonical bases to the overall energy score, and prioritizes an AUGGA motif. Although not drastically different from the canonical A(N)GGA CsrA consensus, the AUGGA motif was independently observed in prior crystal structure [34], SELEX [36], and CLIP-seq [10] studies to be favored by RsmA. This also demonstrates the utility in using solved crystal structures to generate models of protein-RNA interactions. Overall, considering slight changes in the protein sequence allowed for our approach to be better tailored for assessing interactions occurring within *P aeruginosa*.

Our model appears to be able to accurately capture binding interactions between RsmA and candidate targets, as evidenced by the correlation between the measured *in vitro* binding affinities and the predicted ΔG_total_ values that we performed in a small selection of predicted mRNA targets (**Fig. 5c**). More qualitatively, genes that did not pass our energetic threshold (such as *rhlR*) were not observed to bind *in vitro* (**Fig. 5b**), and showed no significant change in translation *in vivo* (**Fig. 6c**). This suggests that the model has utility in predicting relative binding affinity and can aid in further exploration of network regulation, particularly as it relates to lowly expressed or condition-dependent genes. Interestingly, of the 1043 genes predicted to be bound by RsmA, several previously characterized genes did not pass our energy cutoff. These included *magA,* and *mucA*, for which binding was previously experimentally confirmed *in vitro* [1,9]. Each of these predictions yielded less favorable mean ΔG_total_ scores, with only a handful of the suite of binding conformations scoring with high favorability. It is possible then, that other sequences that exhibit strict site ranges may have been lost to filtering. Other genes that did not pass our energetic cutoff included those regulated in tandem with other post-transcriptional regulators, or require multiple copies of RsmA. This is possible as it has been demonstrated that RsmA is not always the sole repressor and can bind genes in tandem with other regulatory factors; this has been shown to occur with two transcriptional regulators, AmrZ and Vfr, wherein RsmA is only able to bind these transcripts in the presence of an additional global post-transcriptional regulator Hfq. [12,19]. Neither *amrZ* nor *vfr* were predicted to be bound by RsmA in our model, therefore our pool of predicted targets is limited to those regulated by RsmA alone.

Future iterations of our model can improve upon capturing the influence of multimerization on binding. RsmA binding can cause structural changes along an RNA transcript and promote multimerization via subsequent folding of higher affinity sites. This phenomenon has been best demonstrated via loading of multiple copies of RsmE on the RsmZ sRNA sponge [39] Our model only considers binding interactions between a single RsmA protein and transcript; therefore, the structural influence of multiple proteins is missed by the model. To address these limitations, future improvements could include structural constraints due to partner binding, however, the footprint and position of the cooperative partner must be known. In addition, changes can be made to have RNA sequences “inherit” structural constraints from a primary iteration of predictions, and measure changes in total affinity due to the addition of secondary or tertiary elements. This can also prove useful in modeling RsmA-mRNA interactions in other Pseudomonas species that encode multiple paralogs of RsmA, such as RsmA and RsmE in *P. putida* and *P. syringae* [32]

Global trends in our binding site predictions agree with patterns observed in prior high throughput screens. Distances between the top binding sites and the start codon were plotted for all genes that passed our total affinity and peak filtering (**Supplementary Fig. 4**). Overall, binding sites for RsmA were localized to three main regions: RBS region (between -30 and 0 relative to the start codon), the start codon, and a broad distribution of sites within the first 100 bases of the coding sequence. This is consistent with binding site frequencies observed in CLIP-seq studies of RsmA in *P. aeruginosa* [10] and CsrA in *E. coli* [57]. These observations suggest that, in addition to predicting an overall affinity score, our model can also predict specific binding sites on the mRNA which provides additional information on the exact mechanism by which the protein interacts with its target.

More globally, binding site distributions vary across transcripts. To investigate this, we used custom peak calling scripts with parameters defined in Methods. A peak is therefore a region with a sufficiently high frequency of predicted sites that passes some minimum threshold set by negative controls. Approximately 30% of genes modeled have wide, overlapping, pockets of binding sites that span 30 + nucleobases across of the mRNA. An example of this is shown in predictions on the *mvaT* transcript (**Fig. 8a).** 70% contain narrower, distinct, peaks that are less than 30 nucleotides wide, which is also seen for predictions across *rsaL* (**Fig. 7a**). Analysis of peak count distributions for our predictions (shown in **Supplementary Fig. 2d**) reveals that the majority of genes have an average number of 1.25 peaks in their distribution of binding site peaks, and a smaller population of genes contain an average of 2.5 peaks where RsmA is predicted to bind. This indicates that the majority of genes contain 1-2 distinct binding peaks, whereas a smaller population contain 2 or more distinct peaks. This recapitulates prior observations that Rsm/Csr proteins facultatively interact with targets at a single binding site, or at double binding sites [4,8]. The divergent patterns of binding also suggest “anchoring” at single high affinity site along the gene, prior to binding to lower affinity positions. This phenomenon was recently characterized for CsrA-*acnA* and *evgA* sequences in *E. coli* [52]

Further, the location of predicted binding peaks appeared to correlate well with *in vitro* experimental evidence. Our initial observation was the concordance of predicted peak location on the well-studied RsmA binding partner *tssA1*. These predictions fell within characterized binding sites on the mRNA sequence (**Fig. 2c**) [36]. The model also accurately predicted high affinity binding sites on the *rsaL* mRNA sequence which had no prior binding or foot-printing evidence. Using *in vitro* filter binding, we experimentally confirmed these predictions by disrupting interactions via mutation of the highest affinity motif (BS3) (**Fig. 7c**), and a further disruption of binding strength was observed upon mutating the second strongest motif (BS2) in tandem with BS3 (**Fig. 7c)**. This is consistent with the theory that Csr/Rsm family proteins may anchor to lower affinity sites on the nascent transcript [19], before binding more strongly to downstream high affinity sites [30]. In contrast, mutating predicted sites along the *mvaT* leader sequence did not result in a change in affinity (**Fig. 8c**). Predicted RsmA binding sites along the *mvaT* sequence cluster in a wide region within the CDS (**Fig. 8a**), and suggest that there may be a multitude of conformations by which RsmA binds to this transcript. This mechanism of binding has been theorized previously [30] as a strategy CsrA to ensure binding to a dynamic structured RNA.

### Loss of target discovery can be attributed to widely varying expression profiles across study conditions

Perhaps the most exciting element of the model results is demonstrating the ability of computational predictions to capture interactions for mRNAs that are expressed transiently or in a condition-dependent manner. Our evaluation of target predictions across 411 gene expression datasets revealed that the majority of novel genes predicted by our model are lowly expressed (**Fig. 4a, b**) or condition specific (**Fig. 4c**). Indeed, K-modal clustering showed a higher ratio of these novel genes to cluster with nonstandard media types like ABTGT or M9 minimal media (**Fig. 4c**). This highlights the importance of considering multiple approaches to profile the effects of a post-transcriptional regulator, as condition dependent gene expression can cause a bottleneck in discovery. This is the case for sRNA discovery, especially, as many are expressed in specific nutrient [58] or infection contexts [59].

### Model identifies that RsmA exerts regulatory control of Quorum Sensing and Biofilm forming pathways through binding and regulation of redundant TF nodes

RsmA is a major global regulator of a variety of pathways that contribute to survival and pathogenicity of *P. aeruginosa*. These include indirect activation of pathways critical for epithelial colonization such as the Type 3 Secretion System (T3SS) [60], Type IV Pili, and flagellar biosynthesis processes[1]. RsmA also has been shown to directly repress pathways that contribute to chronic infection states, such as the formation of biofilms, Quorum Sensing (QS) [53], and the Type 6 Secretion System (T6SS)[6]. Tight control of these processes is advantageous for fitness and survival of PA as it responds to rapid changes in the environment. Direct forms of post-transcriptional regulation typically have a stronger and more immediate effect on gene expression. It is therefore important to effectively differentiate between indirect and direct forms of regulation by RsmA to better understand the influence on dynamic signaling networks. In this study, we used our tuned model to predict the likelihood of a direct interaction occurring between RsmA and an mRNA leader sequence, and found predictions to be enriched for transcriptional regulators and core virulence pathways (**Fig. 3a**). Here, we discuss noteworthy predictions generated for genes in quorum sensing and biofilm forming pathways.

Quorum Sensing (QS) in PA are complex, interconnected, context-dependent signaling cascades that facilitate group control and survival. Gene expression in these pathways is stochastic and sensitive to environmental conditions including fluctuations in nutrients, pH, and cellular density[61,62]. QS expression can also vary from cell to cell in a population, and it is thought that this heterogeneity is a survival strategy that ensures proper division of labor and resource conservations within biofilms[63]. It has also been observed that post-transcriptional regulation by sRNAs and RBPs allows for fine tuning of signal production [64]. These factors present challenges in fully characterizing how these pathways are regulated experimentally, and efforts have been made to understand dynamics using computational modeling [65]. The activation of the hierarchical and interconnected quorum sensing pathways in PA has been shown to directly influence the lifestyle switch towards sessile biofilm forming states. The Gac/Rsm regulatory pathway has been identified as a key influencer of the QS cascade [53]. Our model identified several transcriptional regulators in the QS pathway as potential regulatory targets of RsmA (**Fig. 3a,c**). This included *lasR* and *mvfR* transcriptional activators as well as the *lasI* and *rhlI* homo-serine lactone synthetases. The hierarchical cascade of QS signaling is initiated when transcriptional activator, LasR, is becomes active upon sensing 3-oxo-C12-HSL. This event sets off a signaling cascade and activates expression of subsequent transcriptional regulators RhlR, and MvfR (**Fig. 9**; [66]). There exists an interplay between the RhlR and MvfR, wherein RhlR represses MvfR expression [67]. Interestingly, RsmA binding to *rhlR* was neither predicted nor observed (**Fig. 5a, b, 6b, Supplementary Fig. 3**) which, given the repressive effect RhlR has on *mvfR* transcription, suggests a redundant mechanism by which RsmA regulates expression of this pathway along multiple nodes. Additional QS associated regulators were also evaluated *in vitro* given results of our model, including transcriptional repressors *rsaL* and *mvaT.* Both *rsaL* and *mvaT r*epress elements of the LasR/I QS cascade (**Fig. 9**). *mvaT* has been observed to repress additional transcription factors including *mvfR* [68] and represses *rsaL* in *P. fluorescens* [69].

**Figure 9:**
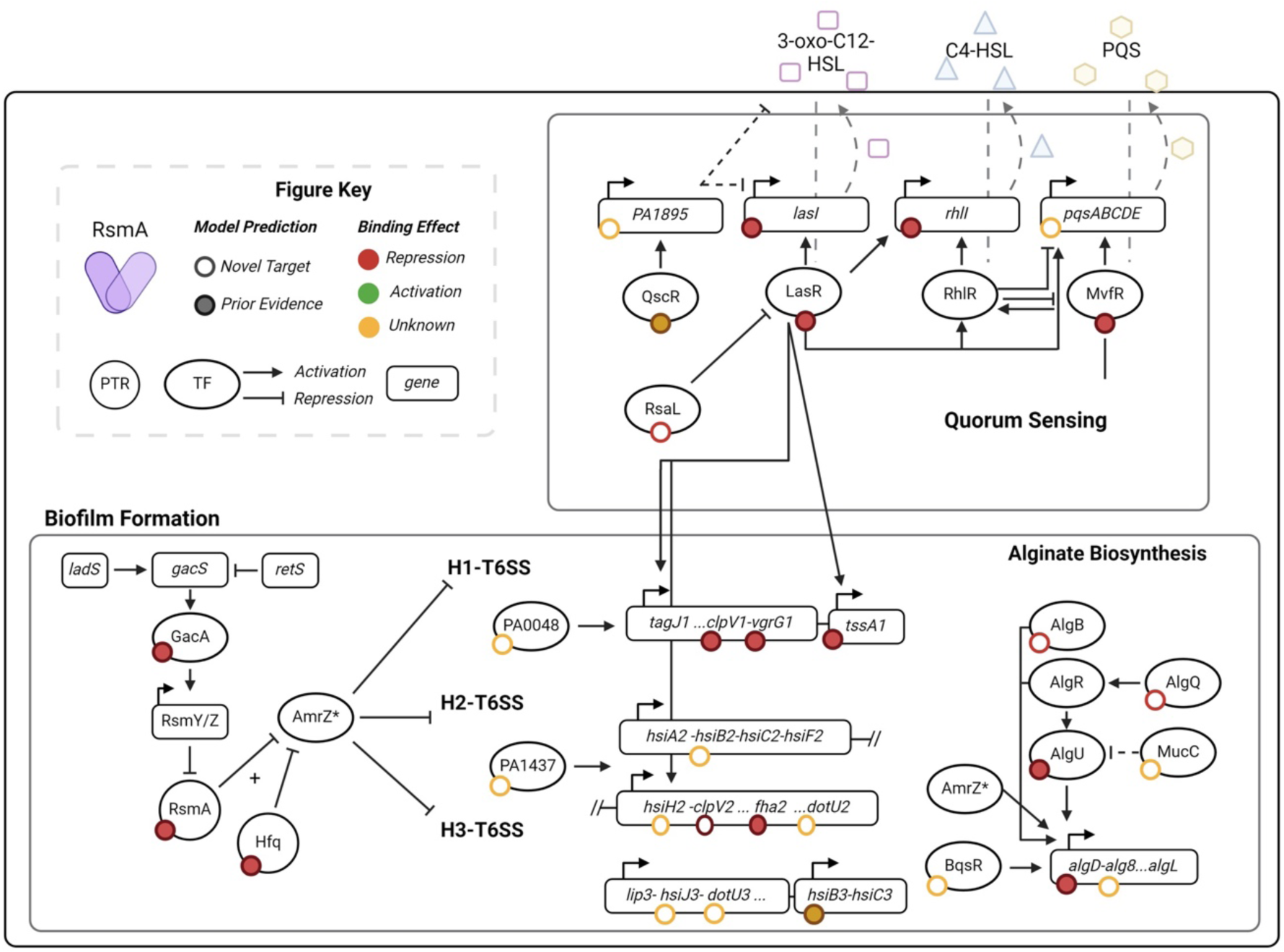
Virulence associated pathways enriched in target predictions included key regulatory transcription factors. Pathway diagrams shown here represent RsmA targets identified by our model in context of their cellular contribution to virulence. Circles represent predictions that passed our filter and are shown in solid or hollow based on whether there is prior experimental evidence of direct or indirect regulation of that gene. In addition, these circles are colored by their predicted regulatory effect: repression (red), activation (green), or an unknown effect (yellow). Genes are shown as boxes, and key transcriptional regulators are present as ovals. Finally post transcriptional regulators such as RsmA and Hfq are shown as circles. As shown in the box describing Quorum Sensing, several key TF regulators are targeted by RsmA, as well as their cognate synthetases that contribute to the autoregulatory feedback loop. Key TFs such as LasR are also directly involved in influencing biofilms, and here we illustrate the activation effect on several pathogenicity islands that make up the T6SS in PA. * Our model did not identify AmrZ as a potential direct target, and this is likely due to the cooperative effect that Hfq binding has on loading RsmA to this gene. We also show predictions for several transcriptional regulators present in the Alginate biosynthesis pathway, providing further clarity on the level of control over this pathway.

Several genes predicted by our model are part of the extensive biofilm formation pathway. Our observation that our model and experimental results confirm binding and repression of LasR led us to further investigate whether RsmA also regulated additional targets of LasR activated genes involved in the T6SS. Inter-operonic binding was observed for genes in the H1, H2, and H3-T6SS (**Fig. 2c**). The GacA/S TCS has been observed to regulate key genes in the H1-T6SS and H3-TCSS, including the well-characterized target *tssA1.* In PA14, the H2-T6SS is more essential than H1[70], and is activated by the QS transcriptional regulator MvfR [71]. The prediction that RsmA regulates of several genes within this locus (**Fig. 3c**), as well as repressing *mvfR,* reflects a shift towards redundant regulatory control of that crucial region. Overall, this outlines the utility of the model in capturing inter-operonic binding events that regulate the assembly of large, multi-component structures in PA.

In our study we further evaluated the strength and regulatory nature of binding between RsmA and the *rsaL* and *mvaT* transcriptional regulators. RsaL was identified as a regulator of four major virulence-associated pathways, including QS (**Fig. 3**,[55]), exhibits low levels of expression across an aggregate of publicly available sequencing data [23], and is an entirely novel prediction generated by our model. In this study, we demonstrate that RsmA binds to this mRNA *in vitro* (**Fig. 5b**) at positions +67 and +76 nucleotides from the start codon (**Fig. 6**).

Binding results in repression of translation of this protein (**Fig. 8c**). We also theorize that this gene evaded prior high throughput screens because of low (**Fig. 4b**), or context dependent expression during planktonic growth phase. The observation that RsmA represses translation of *rsaL* suggests a surprising mechanism of indirect activation, as RsaL negatively regulates *lasI* expression by blocking LasR transcriptional activation [72]. Perhaps this is a mechanism by which RsmA can initiate the autoregulatory feedback loop for the LasR/I signaling cascade at intermediate points during the motile – sessile lifestyle switch.

The second transcript we characterized further was that encoding the MvaT transcriptional repressor. There exists prior evidence of RsmA causing changes in expression[19] or binding directly to this transcript [10], however no prior evidence exists of direct binding *in vitro* or repression *in vivo.* Interestingly, MvaT has also been shown to regulate the Gac/Rsm regulatory pathway through repression of the RsmY and RsmZ sRNA sponges [73]. MvaT is also a regulator of QS, and its influence the system is thought to be through repression of *mvfR* and *rsaL*. In this study, we find that RsmA binds *mvaT* within the coding sequence (**Fig. 7**) and represses expression of *mvaT* as well as its paralog *mvaU* (**Fig. 8c**). Although mutations at model-predicted binding sites did not result in full loss of binding, the width of predicted binding sites on this transcript (**Fig. 7a**) suggests that RsmA may bind in multiple conformations.

In this study, we confirm RsmA binds and represses translation of *lasR, lasI, rhlI, mvfR, rsaL* and *mvaT* (**Fig.s 5-8**). We hypothesize that this mechanism of redundant regulatory control across quorum sensing and biofilm formation allows for tight regulation of energetically costly pathways that can become rapidly de-repressed upon sequestration by the RsmY and RsmZ small RNAs, and could also fine tune production of signaling molecules at intermediate steps along the planktonic to biofilm forming lifestyle switch.

## Supporting information

Supplemental Binding Packet

## Conclusions

This study demonstrates the utility in using thermodynamic modeling for differentiating direct from indirect regulatory interactions between the RsmA protein and the entirety of the transcriptome within PA. Our computational approach yielded novel genes not yet reported to be bound or regulated by the RsmA, likely due to lack of expression in standard laboratory growth conditions. We also affirm the conserved nature of Rsm/Csr regulation across gammaproteobacteria, as known interactions in PA are recapitulated given empirically derived parameters derived from the CsrA protein in *E. coli*. The further biochemical characterization of binding to two transcriptional regulatory targets *mvaT* and *rsaL* reveal that RsmA has a far more extensive influence on quorum sensing pathways. We anticipate that the predictions presented in this dataset will aid in further characterization RsmA regulatory influence upon the complex and interconnected networks within this widespread pathogen.

## Availability of Data and Materials

Scripts and associated files can be found at https://github.com/ajlukasiewicz/rsm_biophysical_model. Raw sequencing data for the RNA Co-Immunoprecipitation Sequencing experiments can be found in the Sequence Read Archive (SRA) under the Bioproject ID PRJNA1131461

## Acknowledgements

We would like to thank Ryan Buchser, Kobe Grismore, Trevor Simmons, and Philip Sweet for their feedback on the manuscript. Figures were generated for this work using Biorender (BioRender.com)

## Author Contributions

Designed research: A.J.L., A.N.L., and L.M.C.; performed experiments: A.J.L. L.H., E.M., C.J.G., B.T.Z., K.H.J.; wrote scripts: A.J.L analyzed data: A.J.L., A.N.L., and L.M.C.; wrote paper: A.J.L., M.J.W., T.L.Y., and L.M.C.

## Description of Additional Files

- Supplementary Table 1: Energy predictions used to generate PWM
- Supplementary Table 2: Sequences Modeled from the PA14 Transcriptome
- Supplementary Table 3: Primers, Plasmids, and Strains used in this work
- Supplementary Table 4: All binding site and translation rate predictions made
- Supplementary Table 5: RsmA targets determined by prior HTS approaches
- Supplementary Table 6: DEseq2 analysis of RNA Co-immunoprecipitation Seq in PA14
- Supplementary Table 7: DEseq2 summary of proteomics dataset
- Supplementary Figures: PDF containing supplemental figures 1-7
- Supplemental Binding Packet: HTML file containing visualizations of binding sites, assigned KEGG pathways, mean predicted affinities, and predicted translational effect for all 1043 targets that passed filtering

