## Supplemental Figures for "Thermodynamic modeling of Csr/Rsm- RNA interactions capture novel, direct binding interactions across the *Pseudomonas aeruginosa* transcriptome"

- **Figure 1 – Predicted effects of binding on translation**
- **Figure 2 – Validation of binding site predictions using previously footprinted mRNAs from closely related organisms**
- **Figure 3 – Summary of RNA Co-Immunoprecipitation Sequencing and Proteomics Results**
- **Figure 4 – RsmA EMSA results for *rhlR, rhlI, aprD,* and *aprX***
- **Figure 5 – UMAP clustering of aggregate sequencing data**
- **Figure 6 – Filter binding images for *rsaL* binding site mutations**
- **Figure 7 - Filter binding images for *mvaT* binding site mutations**

Supplemental Figure 1: Predicted effects of binding on translation

A

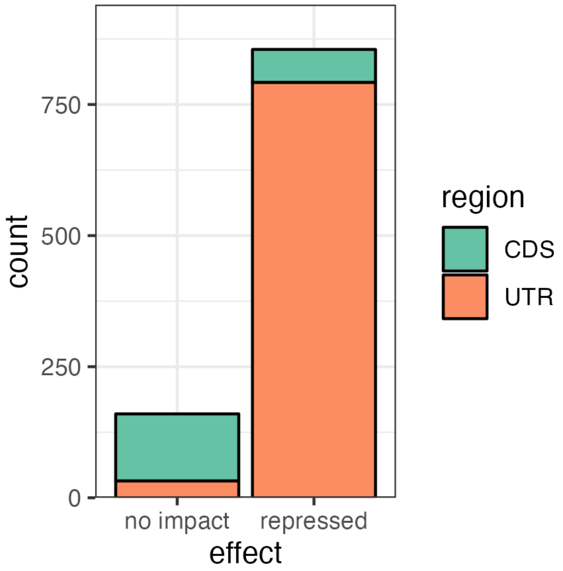


**Figure 1: RsmA effects on translation across the transcriptome.** The location of the binding sites influenced the predicted outcome of binding on translation using the OSTIR translation rate prediction tool. The majority of targets that passed energetic and peak filtering were predicted to be repressed upon binding by RsmA. The minority of these repressed outcomes are due to predictions present in the coding sequence (CDS) as opposed to the untranslated region (UTR). The majority of targets for which no regulatory outcome was predicted due to binding has predicted binding sites in the CDS relative to the UTR.

### Supplemental Figure 2: Validation of model predictions using previously characterized binding sites

**A** ***Escherichia coli* CsrA-*glgC***

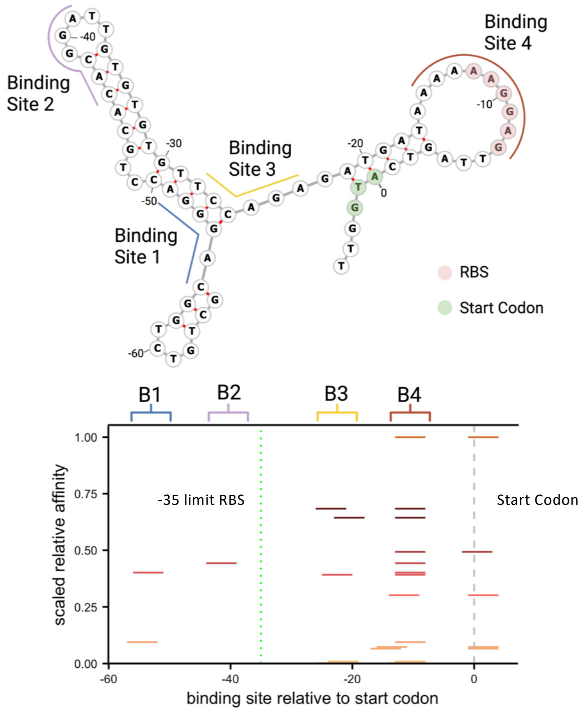


**C** ***Pseudomonas aeruginosa* RsmA-*tssA1***

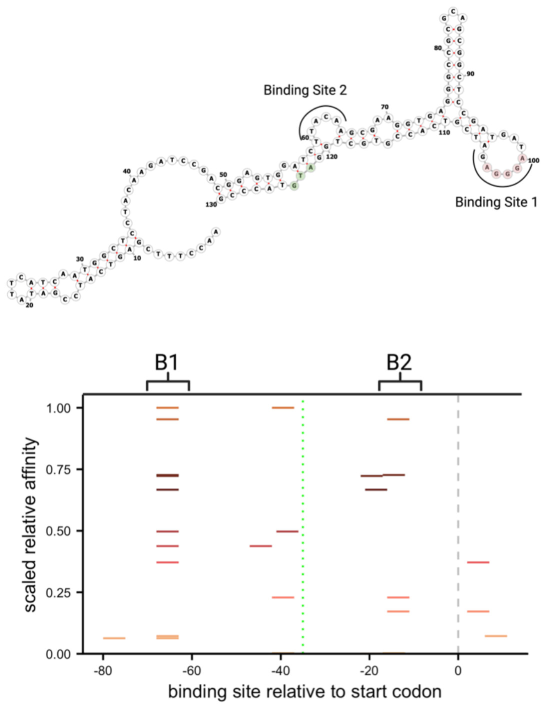


**B** ***Pseudomonas protegens* RsmE-*hcnA***

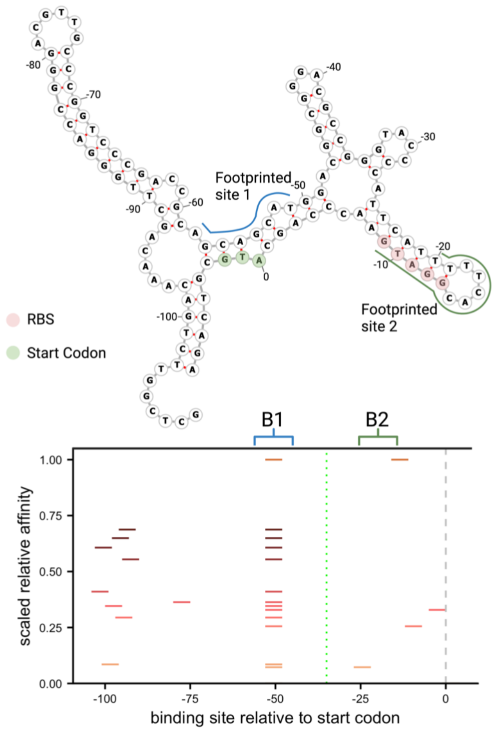


**D** **Binding peak frequencies across all filtered genes**

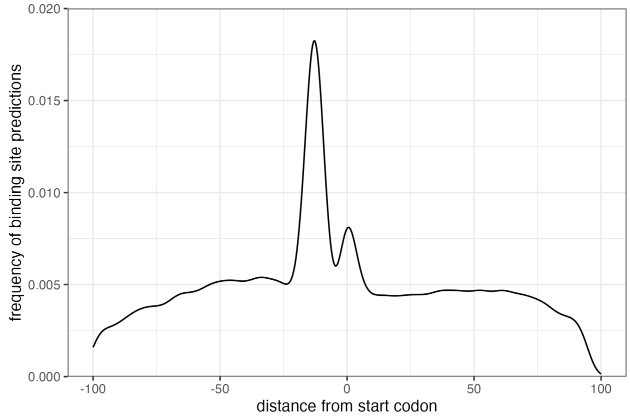


**Figure 2: Applying our biophysical model to previously characterized Rsm-UTR interactions across three organisms validated broad applicability across Rsm/Csr encoding bacteria**

**A.** 4 sites on the *glgC* UTR were previously found to be bound by CsrA in *E. coli* (Mercante et. al., 2009)**.** Applying our CsrA based model to this target correctly captured each binding site, witch a positive predictive value PPV of 0.77. **B.** Similarly as in A, we applied the same model to the *hcnA* UTR present in *P. protegens.* Binding sites of the CsrA homolog RsmE on the *hcnA* UTR were captured in (Lapouge et. al., 2007). Top predicted conformations centered around footprinted site 1. **C.** Within *P. aeruginosa*, the *tssA1* UTR is a well characterized target of the RsmA protein and two putative binding sites characterized in (Schulmeyer et. al., 2016) were accurately predicted by our model. **D.** Density plot of all binding peak frequencies plotted for all genes that pass the energetic and binding peak calling filters.

### Supplemental Figure 3: RNA Co-Immunoprecipitation Sequencing and Proteomics Results

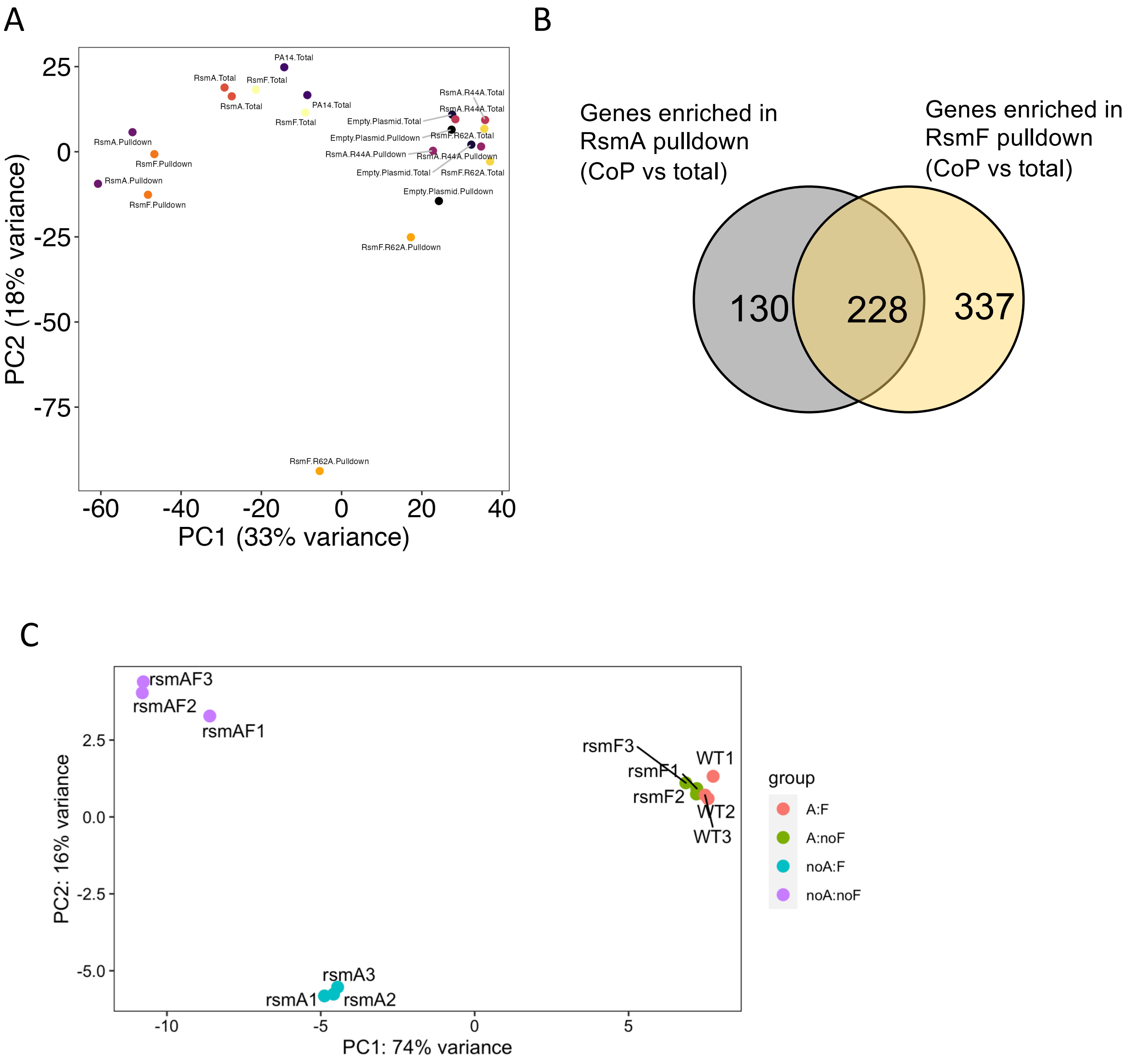


**Figure 3:** PCA analysis of counts data for RNA co-immunoprecipitation sequencing and proteomics on PA14 and PA103 respectively. A) PCA plot of RNA Co-Immunoprecipitation Sequencing read counts for pulled down and total RNA fractions. B) Venn diagram of exclusive and overlapping RNAs pulled down by either his-tagged RsmA or RsmF. C) PCA of protein counts in WT, delA, delF, and delAF PA103

Supplemental Figure 4: *rhlR, rhlI, aprD* and *aprX* EMSA results

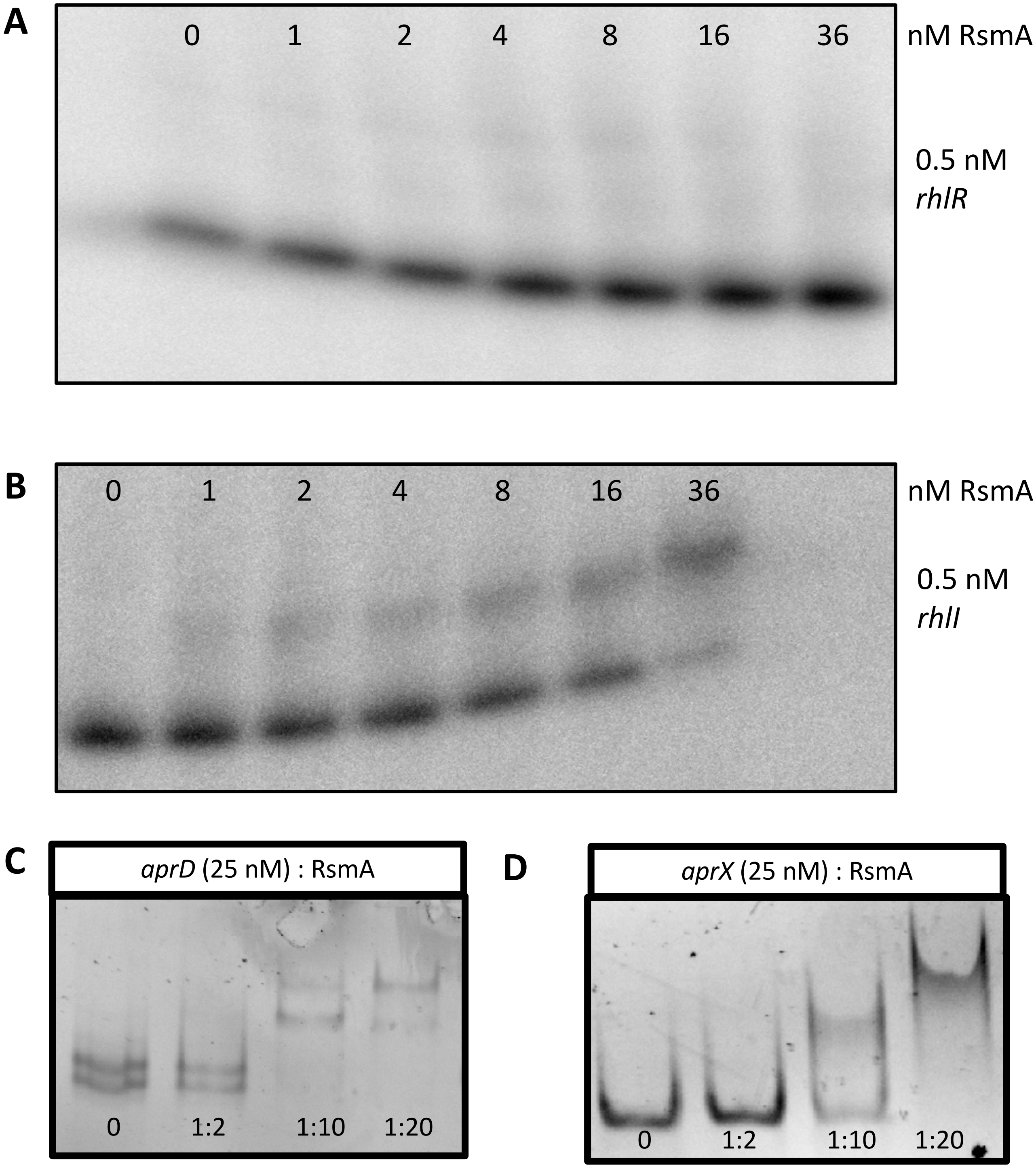


**Figure 4:** Electrophoretic Mobility Shift Assay to assess binding of RsmA to the 5' leader sequence and first 100 bases of A) *rhlR* (no binding observed) and B) *rhlI* (kD = 20.88 $\pm$ 7.5 nM). Un-radiolabeled EMSA was performed between the 5' UTR sequence and first 100 bases of C) *aprD* and D) *aprX* RNAs

Supplemental Figure 5: UMAP clustering of sequencing data aggregated in [4]

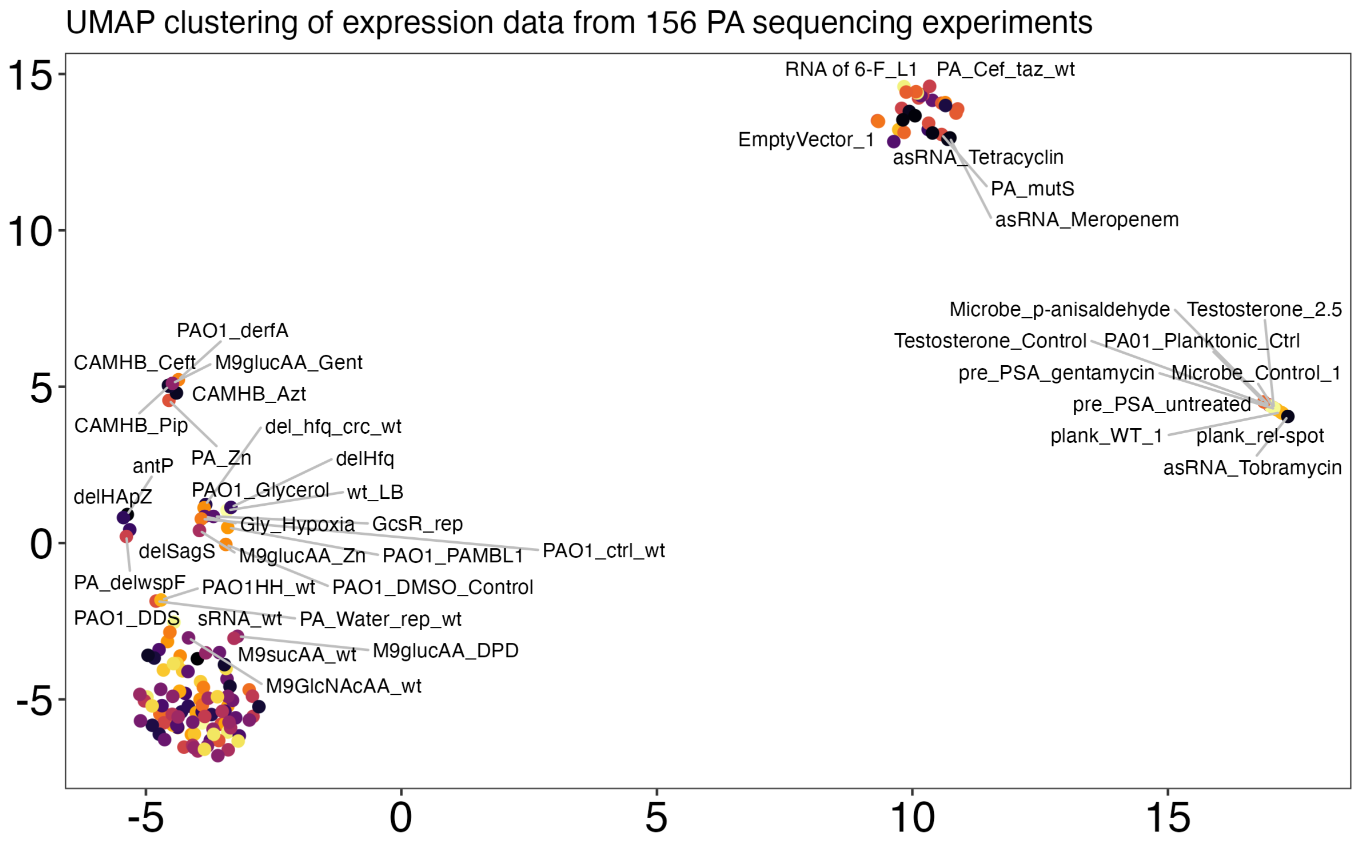

### Supplemental Figure 6

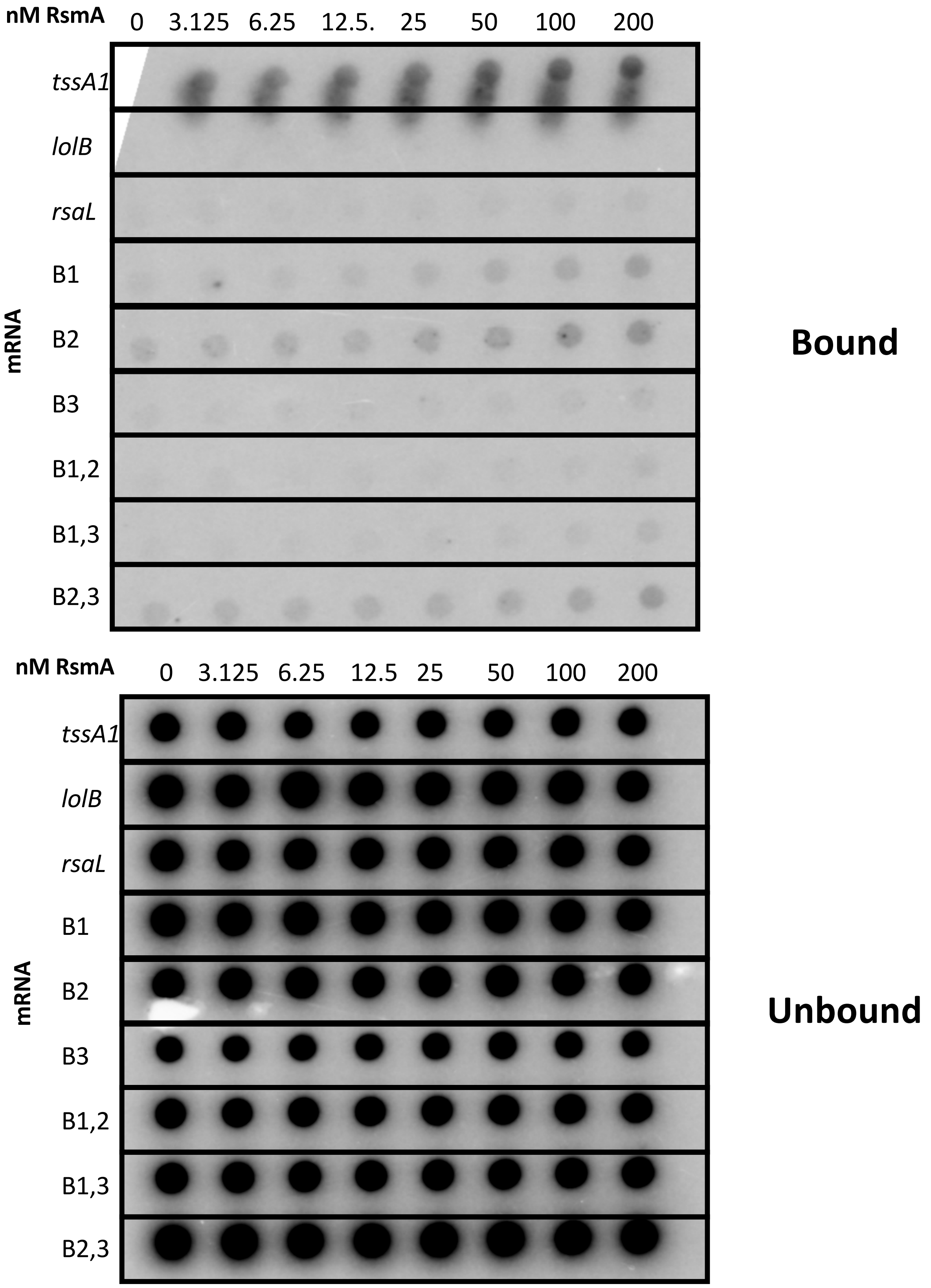


**Figure 6:** Filter-binding assay results for evaluating RsmA binding to *rsaL* and individual mutants at binding sites BS1, BS2, and BS3. Positive and negative controls *tssA1* and *lolB* are shown in the first two rows of the membrane. Bound fractions are shown on the top nitrocellulose membrane, and the unbound RNA flow through is shown on the bottom N+ membrane.

### Supplemental Figure 7

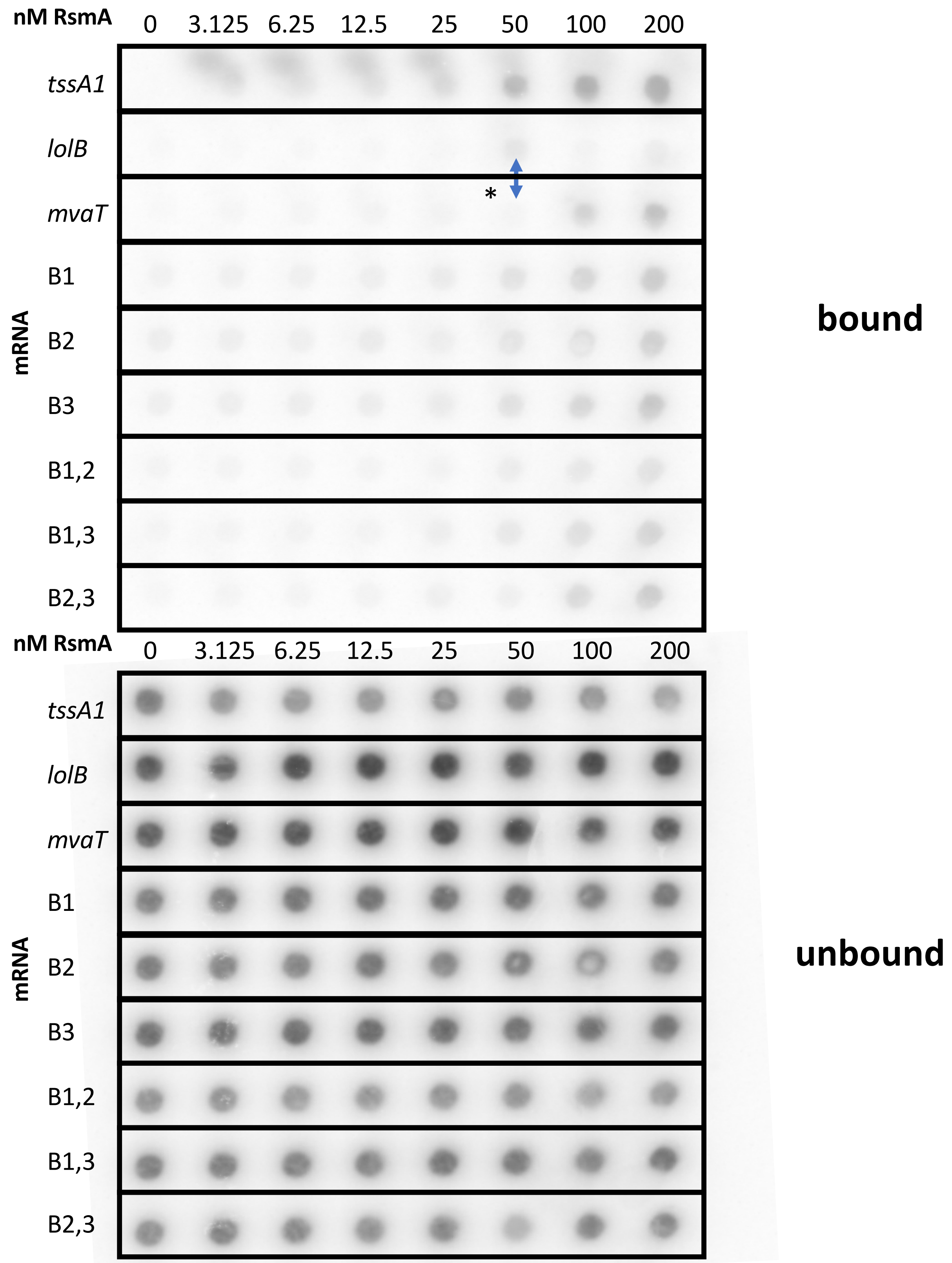


**Figure 7:** Filter-binding assay results for evaluating RsmA binding to *mvaT* and individual mutants at binding sites BS1, BS2, and BS3. Positive and negative controls *tssA1* and *lolB* are shown in the first two rows of the membrane. Bound fractions are shown on the top nitrocellulose membrane, and the unbound RNA flow through is shown on the bottom N+ membrane.
