## Supplemental Binding Packet for "Thermodynamic modeling of Csr/Rsm- RNA interactions capture novel, direct binding interactions across the *Pseudomonas aeruginosa* transcriptome": Supplemental_binding_packet.html

|  |  |  |  |  |  |  |  |
| --- | --- | --- | --- | --- | --- | --- | --- |
| Modeled binding peaks for 1043 predicted targets of RsmA | | | | | | | |
| PA14 gene ID | gene name | description | GO terms | KEGG pathways | overall affinity score in RT | predicted effect on translation | binding site predictions |
| PA14\_00050 | gyrB | DNA gyrase subunit B | DNA binding, DNA topoisomerase type II (ATP-hydrolyzing) activity, ATP binding, DNA topological change, chromosome |  | -26.28105 | repression |  |
| PA14\_00090 | glyS | glycyl-tRNA synthetase subunit beta | arginine-tRNA ligase activity, ATP binding, arginyl-tRNA aminoacylation, nucleotide binding, glycine-tRNA ligase activity, cytoplasm, glycyl-tRNA aminoacylation | Aminoacyl-tRNA biosynthesis | -25.86171 | repression |  |
| PA14\_00160 | NA | hypothetical protein | protein binding |  | -25.96560 | no impact |  |
| PA14\_00210 | NA | lysin domain-containing protein |  |  | -26.23488 | repression |  |
| PA14\_00230 | NA | Rossmann fold nucleotide-binding protein | DNA mediated transformation |  | -26.91108 | repression |  |
| PA14\_00250 | qor | quinone oxidoreductase | oxidation-reduction process, oxidoreductase activity, zinc ion binding |  | -25.87822 | repression |  |
| PA14\_00290 | aroE | shikimate 5-dehydrogenase | shikimate 3-dehydrogenase (NADP+) activity, shikimate metabolic process, NADP binding, oxidation-reduction process | Biosynthesis of amino acids, Biosynthesis of antibiotics, Biosynthesis of secondary metabolites, Metabolic pathways, Phenylalanine, tyrosine and tryptophan biosynthesis | -26.42002 | repression |  |
| PA14\_00340 | NA | sulfate transporter | sulfate transport, sulfate transmembrane transporter activity, integral component of membrane |  | -25.84300 | repression |  |
| PA14\_00530 | NA | hypothetical protein |  |  | -26.77829 | repression |  |
| PA14\_00580 | NA | lipoprotein |  |  | -26.67633 | repression |  |
| PA14\_00600 | NA | transcriptional regulator | DNA binding |  | -26.14631 | repression |  |
| PA14\_00630 | NA | hypothetical protein |  |  | -26.95979 | repression |  |
| PA14\_00660 | NA | RNA 2'-phosphotransferase-like protein | phosphotransferase activity, alcohol group as acceptor, tRNA splicing, via endonucleolytic cleavage and ligation, transferase activity, transferring phosphorus-containing groups |  | -27.18575 | repression |  |
| PA14\_00730 | NA | hypothetical protein |  |  | -25.87712 | repression |  |
| PA14\_00740 | NA | lipoprotein |  |  | -26.44604 | repression |  |
| PA14\_00820 | tagQ1 | TagQ1 |  |  | -26.29270 | repression |  |
| PA14\_00875 | ppkA | serine/threonine protein kinase PpkA | protein kinase activity, ATP binding, protein phosphorylation, positive regulation of protein secretion, protein serine/threonine kinase activity | Bacterial secretion system, Biofilm formation - Pseudomonas aeruginosa | -26.70750 | repression |  |
| PA14\_00990 | tssA1 | TssA1 |  |  | -26.34988 | repression |  |
| PA14\_01010 | hsiB1 | HsiB1 |  |  | -26.38080 | repression |  |
| PA14\_01030 | hcp1 | Hcp1 |  |  | -26.72812 | no impact |  |
| PA14\_01100 | clpV1 | ClpV1 | protein metabolic process, ATP binding | Bacterial secretion system, Biofilm formation - Pseudomonas aeruginosa | -25.80157 | repression |  |
| PA14\_01110 | vgrG1a | VgrG1a |  |  | -26.97112 | repression |  |
| PA14\_01120 | tsi6 | Tsi6 |  |  | -26.44927 | repression |  |
| PA14\_01170 | NA | hypothetical protein |  |  | -26.34650 | repression |  |
| PA14\_01180 | NA | hypothetical protein |  |  | -26.19782 | no impact |  |
| PA14\_01200 | NA | hypothetical protein |  |  | -26.15251 | repression |  |
| PA14\_01240 | NA | carbonic anhydrase | carbon utilization, carbonate dehydratase activity, zinc ion binding | Nitrogen metabolism | -26.50859 | no impact |  |
| PA14\_01290 | coxB | cytochrome c oxidase subunit II | cytochrome-c oxidase activity, copper ion binding, membrane, electron transfer activity, heme binding, integral component of membrane, oxidoreductase activity, electron transport chain | Metabolic pathways, Oxidative phosphorylation | -26.30912 | repression |  |
| PA14\_01300 | coxA | cytochrome c oxidase subunit I | oxidation-reduction process, cytochrome-c oxidase activity, respiratory chain complex IV, aerobic respiration, integral component of membrane, heme binding, electron transport coupled proton transport | Metabolic pathways, Oxidative phosphorylation | -26.12748 | no impact |  |
| PA14\_01340 | NA | hypothetical protein | membrane |  | -26.61273 | repression |  |
| PA14\_01380 | NA | protoheme IX farnesyltransferase | integral component of membrane, transferase activity, transferring alkyl or aryl (other than methyl) groups, protoheme IX farnesyltransferase activity, heme O biosynthetic process | Biosynthesis of secondary metabolites, Metabolic pathways, Oxidative phosphorylation, Porphyrin and chlorophyll metabolism | -26.83773 | repression |  |
| PA14\_01490 | NA | hemolysin | hemolysis by symbiont of host erythrocytes |  | -26.91096 | repression |  |
| PA14\_01500 | NA | transcriptional regulator | DNA-binding transcription factor activity, regulation of transcription, DNA-templated |  | -27.00465 | repression |  |
| PA14\_01600 | NA | aldehyde dehydrogenase | oxidoreductase activity, oxidoreductase activity, acting on the aldehyde or oxo group of donors, NAD or NADP as acceptor, oxidation-reduction process, methylmalonate-semialdehyde dehydrogenase (acylating) activity, beta-alanine biosynthetic process | beta-Alanine metabolism, Carbon metabolism, Inositol phosphate metabolism, Metabolic pathways, Propanoate metabolism, Valine, leucine and isoleucine degradation | -27.26365 | repression |  |
| PA14\_01660 | NA | guanine deaminase | guanine catabolic process, zinc ion binding, guanine deaminase activity, hydrolase activity, hydrolase activity, acting on carbon-nitrogen (but not peptide) bonds | drosopterin and aurodrosopterin biosynthesis, guanosine nucleotides degradation II, guanosine nucleotides degradation III, Metabolic pathways, Purine metabolism, Purine metabolism, purine nucleobases degradation II (anaerobic) | -26.51067 | repression |  |
| PA14\_01810 | NA | oxidoreductase | oxidoreductase activity, oxidation-reduction process |  | -25.84014 | repression |  |
| PA14\_01940 | NA | RND efflux membrane fusion protein | membrane, transmembrane transporter activity, transmembrane transport |  | -26.67288 | repression |  |
| PA14\_01970 | NA | RND efflux transporter | membrane, transmembrane transporter activity, transmembrane transport |  | -25.82186 | repression |  |
| PA14\_02020 | NA | outer membrane porin | integral component of membrane |  | -26.21154 | no impact |  |
| PA14\_02140 | NA | hypothetical protein |  |  | -26.04698 | repression |  |
| PA14\_02150 | NA | hypothetical protein | signal transduction, integral component of membrane, catalytic activity |  | -26.27045 | repression |  |
| PA14\_02200 | NA | chemotaxis protein methyltransferase | S-adenosylmethionine-dependent methyltransferase activity | Bacterial chemotaxis, Two-component system | -26.32595 | no impact |  |
| PA14\_02220 | NA | chemotaxis transducer | signal transduction, membrane, transmembrane signaling receptor activity, chemotaxis, integral component of membrane | Bacterial chemotaxis, Two-component system | -27.05187 | repression |  |
| PA14\_02260 | NA | two-component response regulator | phosphorelay signal transduction system | Bacterial chemotaxis, Two-component system | -25.75452 | repression |  |
| PA14\_02300 | fabG | 3-ketoacyl-ACP reductase |  |  | -25.92959 | repression |  |
| PA14\_02310 | atsA | arylsulfatase | catalytic activity, sulfuric ester hydrolase activity, arylsulfatase activity, phosphoric diester hydrolase activity | Sphingolipid metabolism | -26.04159 | repression |  |
| PA14\_02370 | NA | porin | integral component of membrane |  | -26.88642 | no impact |  |
| PA14\_02435 | NA | hypothetical protein | oxidoreductase activity, oxidation-reduction process |  | -26.10635 | repression |  |
| PA14\_02460 | NA | NAD(P) transhydrogenase subunit alpha part 2 |  |  | -25.75933 | repression |  |
| PA14\_02570 | mdcC | malonate decarboxylase subunit delta |  |  | -25.77812 | repression |  |
| PA14\_02590 | mdcE | malonate decarboxylase subunit gamma | cellular carbohydrate metabolic process, ligase activity | 3-hydroxypropanoate cycle, 3-hydroxypropanoate/4-hydroxybutanate cycle, candicidin biosynthesis, fatty acid biosynthesis initiation I, glyoxylate assimilation, malonate degradation II (biotin-dependent) | -25.89917 | repression |  |
| PA14\_02740 | NA | class II aldolase/adducin domain-containing protein |  |  | -25.84622 | repression |  |
| PA14\_02790 | pcaF | beta-ketoadipyl CoA thiolase | transferase activity, transferring acyl groups other than amino-acyl groups, catalytic activity, 3,4-dihydroxybenzoate catabolic process | Benzoate degradation, Microbial metabolism in diverse environments | -25.76400 | repression |  |
| PA14\_02810 | pcaT | dicarboxylic acid transporter PcaT | integral component of membrane, transmembrane transporter activity, transmembrane transport, integral component of plasma membrane |  | -25.91422 | repression |  |
| PA14\_02890 | NA | hypothetical protein | cell outer membrane |  | -26.00684 | no impact |  |
| PA14\_03170 | NA | hypothetical protein | DNA integration, nucleic acid binding |  | -25.76743 | repression |  |
| PA14\_03200 | tle3 | Type VI effector Tle3 |  |  | -26.28507 | repression |  |
| PA14\_03220 | vgrG2b | VgrG2b |  |  | -27.48331 | repression |  |
| PA14\_03510 | NA | hypothetical protein |  |  | -25.93118 | repression |  |
| PA14\_03530 | NA | transcriptional regulator | regulation of transcription, DNA-templated, DNA-binding transcription factor activity |  | -26.15826 | repression |  |
| PA14\_03610 | NA | Zn-dependent protease with chaperone function | metalloendopeptidase activity, proteolysis |  | -26.17998 | repression |  |
| PA14\_03680 | cysT | sulfate transport protein CysT | sulfate transport, ATPase-coupled sulfate transmembrane transporter activity, membrane, plasma membrane, transmembrane transport | ABC transporters, Sulfur metabolism | -26.41424 | repression |  |
| PA14\_03760 | NA | sodium:solute symporter | membrane, transmembrane transporter activity, transmembrane transport |  | -26.22386 | no impact |  |
| PA14\_03930 | spuE | polyamine transport protein | polyamine transport, polyamine binding, periplasmic space | ABC transporters | -26.04000 | repression |  |
| PA14\_04080 | NA | ABC transporter permease | membrane, transmembrane transport, integral component of membrane, transmembrane transporter activity, ATP-binding cassette (ABC) transporter complex, nitrogen compound transport | ABC transporters | -26.71677 | repression |  |
| PA14\_04100 | NA | hypothetical protein |  |  | -26.01195 | repression |  |
| PA14\_04250 | NA | ABC transporter ATP-binding protein | ATP binding, transmembrane transporter activity, ATP-binding cassette (ABC) transporter complex, transmembrane transport, ATPase activity, ATPase-coupled polyamine transmembrane transporter activity, polyamine transport | ABC transporters | -25.99923 | no impact |  |
| PA14\_04330 | NA | hypothetical protein |  |  | -27.43375 | repression |  |
| PA14\_04410 | ptsP | phosphoenolpyruvate-protein phosphotransferase PtsP | transferase activity, transferring phosphorus-containing groups, protein binding, phosphorylation, phosphoenolpyruvate-dependent sugar phosphotransferase system, catalytic activity | Phosphotransferase system (PTS) | -26.54031 | repression |  |
| PA14\_04440 | NA | permease | integral component of membrane |  | -26.02724 | repression |  |
| PA14\_04490 | NA | hypothetical protein |  |  | -26.00842 | no impact |  |
| PA14\_04560 | NA | hypothetical protein |  |  | -26.42844 | repression |  |
| PA14\_04700 | NA | hypothetical protein | protein binding |  | -26.15406 | repression |  |
| PA14\_04950 | mtgA | monofunctional biosynthetic peptidoglycan transglycosylase | peptidoglycan biosynthetic process, peptidoglycan-based cell wall, integral component of membrane, transferase activity, transferring pentosyl groups | Peptidoglycan biosynthesis | -26.61698 | repression |  |
| PA14\_04960 | NA | hypothetical protein |  |  | -26.63033 | repression |  |
| PA14\_05030 | NA | hypothetical protein |  |  | -25.96998 | repression |  |
| PA14\_05050 | NA | deoxyribonucleotide triphosphate pyrophosphatase | nucleoside triphosphate catabolic process, nucleoside-triphosphate diphosphatase activity, nucleoside-triphosphatase activity | Purine metabolism | -25.98232 | repression |  |
| PA14\_05060 | NA | hypothetical protein |  |  | -25.75795 | repression |  |
| PA14\_05110 | NA | hypothetical protein |  |  | -25.76702 | repression |  |
| PA14\_05180 | pilT | twitching motility protein PilT | ATP binding, type IV pilus, pilus retraction, type IV pilus-dependent motility |  | -25.86196 | repression |  |
| PA14\_05310 | gshB | glutathione synthetase | glutathione synthase activity, ATP binding, glutathione biosynthetic process, metal ion binding | Cysteine and methionine metabolism, Glutathione metabolism, Metabolic pathways | -25.79507 | no impact |  |
| PA14\_05320 | pilG | twitching motility protein PilG | phosphorelay signal transduction system | Biofilm formation - Pseudomonas aeruginosa, Two-component system | -26.10482 | repression |  |
| PA14\_05390 | chpA | ChpA | phosphorelay signal transduction system, chemotaxis, signal transduction, phosphorelay sensor kinase activity, protein histidine kinase activity, cytoplasm, phosphorylation, transferase activity, transferring phosphorus-containing groups | Biofilm formation - Pseudomonas aeruginosa, Two-component system | -27.23070 | repression |  |
| PA14\_05540 | mexB | RND multidrug efflux transporter MexB | membrane, transmembrane transporter activity, transmembrane transport, efflux transmembrane transporter activity, integral component of membrane, xenobiotic transport | beta-Lactam resistance, Cationic antimicrobial peptide (CAMP) resistance | -26.63600 | repression |  |
| PA14\_05550 | oprM | major intrinsic multiple antibiotic resistance efflux outer membrane protein OprM precursor | membrane, transmembrane transporter activity, transmembrane transport, efflux transmembrane transporter activity, drug transmembrane transporter activity | beta-Lactam resistance, Quorum sensing | -25.93586 | repression |  |
| PA14\_05620 | sahH | S-adenosyl-L-homocysteine hydrolase | adenosylhomocysteinase activity | <i>S</i>-adenosyl-L-methionine cycle II, Cysteine and methionine metabolism, Cysteine and methionine metabolism, Metabolic pathways | -25.77400 | repression |  |
| PA14\_05890 | NA | stomatin-like protein | membrane |  | -26.62783 | repression |  |
| PA14\_06090 | NA | hypothetical protein |  |  | -26.56362 | no impact |  |
| PA14\_06150 | NA | hypothetical protein |  |  | -25.93607 | repression |  |
| PA14\_06280 | NA | hypothetical protein |  |  | -26.29357 | no impact |  |
| PA14\_06390 | NA | hypothetical protein |  |  | -27.06072 | repression |  |
| PA14\_06400 | NA | LysR family transcriptional regulator | DNA-binding transcription factor activity, regulation of transcription, DNA-templated |  | -25.95936 | repression |  |
| PA14\_06420 | NA | hypothetical protein | catalytic activity, carbohydrate metabolic process | &gamma;-glutamyl cycle, 5-oxo-L-proline metabolism, Glutathione metabolism | -26.03131 | no impact |  |
| PA14\_06620 | NA | acyl-CoA dehydrogenase | oxidoreductase activity, acting on the CH-CH group of donors, flavin adenine dinucleotide binding, oxidation-reduction process |  | -26.01378 | repression |  |
| PA14\_06750 | nirS | nitrite reductase | electron transfer activity, heme binding | Microbial metabolism in diverse environments, Nitrogen metabolism | -26.54933 | repression |  |
| PA14\_06890 | NA | hypothetical protein | catalytic activity, molybdenum ion binding, pyridoxal phosphate binding |  | -25.83427 | repression |  |
| PA14\_06920 | NA | class III pyridoxal phosphate-dependent aminotransferase | transaminase activity, pyridoxal phosphate binding, catalytic activity | 2-Oxocarboxylic acid metabolism, Arginine biosynthesis, Biosynthesis of amino acids, Biosynthesis of antibiotics, Biosynthesis of secondary metabolites, Lysine biosynthesis, Metabolic pathways, Microbial metabolism in diverse environments | -26.20679 | repression |  |
| PA14\_06940 | NA | hypothetical protein |  |  | -26.38598 | repression |  |
| PA14\_07010 | NA | hypothetical protein | membrane, integral component of membrane |  | -26.50039 | repression |  |
| PA14\_07070 | NA | hypothetical protein | oxidoreductase activity, oxidation-reduction process |  | -26.43337 | repression |  |
| PA14\_07200 | NA | hypothetical protein |  |  | -26.63206 | repression |  |
| PA14\_07210 | NA | hypothetical protein |  |  | -26.51305 | repression |  |
| PA14\_07250 | NA | hypothetical protein |  |  | -26.45753 | repression |  |
| PA14\_07300 | NA | hypothetical protein |  |  | -26.33994 | repression |  |
| PA14\_07330 | NA | hypothetical protein | integral component of membrane |  | -26.87012 | repression |  |
| PA14\_07370 | NA | hypothetical protein | integral component of membrane |  | -26.04589 | repression |  |
| PA14\_07380 | NA | hypothetical protein |  |  | -26.97839 | repression |  |
| PA14\_07420 | NA | hypothetical protein |  |  | -25.76632 | no impact |  |
| PA14\_07520 | rpoD | RNA polymerase sigma factor RpoD | DNA-binding transcription factor activity, DNA-templated transcription, initiation, regulation of transcription, DNA-templated, DNA binding, sigma factor activity |  | -25.94098 | no impact |  |
| PA14\_07590 | folB | dihydroneopterin aldolase | dihydroneopterin aldolase activity, folic acid-containing compound metabolic process | Folate biosynthesis, Metabolic pathways | -26.55826 | repression |  |
| PA14\_07630 | NA | hypothetical protein |  |  | -25.88896 | no impact |  |
| PA14\_07680 | NA | hypothetical protein | protein kinase activity |  | -27.09370 | repression |  |
| PA14\_07840 | NA | two-component response regulator | phosphorelay signal transduction system, regulation of transcription, DNA-templated, DNA binding |  | -26.09922 | no impact |  |
| PA14\_07850 | NA | ABC transporter substrate-binding protein |  |  | -26.15679 | repression |  |
| PA14\_07900 | NA | ABC transporter permease | membrane, transmembrane transport | Quorum sensing | -26.35649 | repression |  |
| PA14\_08020 | NA | bacteriophage protein |  |  | -26.81742 | repression |  |
| PA14\_08060 | NA | tail fiber assembly protein |  |  | -26.23744 | repression |  |
| PA14\_08160 | NA | lytic enzyme |  |  | -26.58651 | repression |  |
| PA14\_08200 | NA | hypothetical protein |  |  | -26.00534 | repression |  |
| PA14\_08210 | NA | hypothetical protein |  |  | -25.94224 | repression |  |
| PA14\_08240 | NA | hypothetical protein |  |  | -26.80327 | no impact |  |
| PA14\_08250 | NA | hypothetical protein |  |  | -26.25859 | repression |  |
| PA14\_08300 | NA | phage-related protein, tail component | protein binding |  | -26.38628 | repression |  |
| PA14\_08310 | NA | hypothetical protein |  |  | -25.88211 | repression |  |
| PA14\_08320 | NA | hypothetical protein |  |  | -25.76658 | no impact |  |
| PA14\_08350 | trpD | anthranilate phosphoribosyltransferase | tryptophan biosynthetic process, anthranilate phosphoribosyltransferase activity, transferase activity, transferring glycosyl groups | Biosynthesis of amino acids, Biosynthesis of antibiotics, Biosynthesis of secondary metabolites, Metabolic pathways, Phenylalanine, tyrosine and tryptophan biosynthesis | -27.06760 | repression |  |
| PA14\_08420 | NA | HIT family protein | catalytic activity |  | -25.96506 | repression |  |
| PA14\_08480 | argC | N-acetyl-gamma-glutamyl-phosphate reductase | cellular amino acid biosynthetic process, oxidoreductase activity, acting on the aldehyde or oxo group of donors, NAD or NADP as acceptor, protein dimerization activity, N-acetyl-gamma-glutamyl-phosphate reductase activity, arginine biosynthetic process, oxidation-reduction process, NAD binding | 2-Oxocarboxylic acid metabolism, Arginine biosynthesis, Arginine biosynthesis, Biosynthesis of amino acids, Biosynthesis of antibiotics, Biosynthesis of secondary metabolites, L-arginine biosynthesis III (via <i>N</i>-acetyl-L-citrulline), L-arginine biosynthesis IV (archaebacteria), Metabolic pathways | -26.28718 | repression |  |
| PA14\_08630 | NA | pantothenate kinase | pantothenate kinase activity | Metabolic pathways, Pantothenate and CoA biosynthesis | -26.33798 | repression |  |
| PA14\_08680 | tufB | elongation factor Tu | translation elongation factor activity, GTP binding, translational elongation, GTPase activity |  | -26.59689 | repression |  |
| PA14\_08730 | rplA | 50S ribosomal protein L1 | RNA binding, structural constituent of ribosome, translation, large ribosomal subunit | Ribosome | -25.81372 | repression |  |
| PA14\_08760 | rpoB | DNA-directed RNA polymerase subunit beta | DNA-directed 5'-3' RNA polymerase activity, transcription, DNA-templated, ribonucleoside binding, DNA binding | Metabolic pathways, Purine metabolism, Pyrimidine metabolism, RNA polymerase | -25.76213 | no impact |  |
| PA14\_08900 | rplV | 50S ribosomal protein L22 | structural constituent of ribosome, ribosome, translation, large ribosomal subunit | Ribosome | -27.00943 | repression |  |
| PA14\_08910 | rpsC | 30S ribosomal protein S3 | structural constituent of ribosome, translation, small ribosomal subunit, nucleic acid binding, RNA binding | Ribosome | -26.74368 | repression |  |
| PA14\_08950 | rplN | 50S ribosomal protein L14 | structural constituent of ribosome, ribosome, translation, large ribosomal subunit | Ribosome | -25.80334 | repression |  |
| PA14\_08960 | rplX | 50S ribosomal protein L24 | structural constituent of ribosome, ribosome, translation | Ribosome | -25.98982 | repression |  |
| PA14\_08970 | rplE | 50S ribosomal protein L5 | structural constituent of ribosome, ribosome, translation | Ribosome | -26.33277 | no impact |  |
| PA14\_09270 | pchE | dihydroaeruginoic acid synthetase | catalytic activity, phosphopantetheine binding | Biosynthesis of siderophore group nonribosomal peptides, Pyochelin synthesis | -26.00953 | repression |  |
| PA14\_09340 | fptA | Fe(III)-pyochelin outer membrane receptor | cell outer membrane, siderophore uptake transmembrane transporter activity, siderophore transport, signaling receptor activity |  | -26.44751 | repression |  |
| PA14\_09410 | phzG1 | pyrodoxamine 5'-phosphate oxidase | oxidoreductase activity, acting on the CH-NH2 group of donors, oxidation-reduction process, pyridoxamine-phosphate oxidase activity, pyridoxine biosynthetic process, FMN binding, cofactor binding, phenazine biosynthetic process | Phenazine biosynthesis, Quorum sensing | -28.31133 | repression |  |
| PA14\_09420 | phzF1 | phenazine biosynthesis protein | catalytic activity, biosynthetic process | Phenazine biosynthesis, Quorum sensing | -26.44280 | NA |  |
| PA14\_09440 | phzE1 | phenazine biosynthesis protein PhzE | biosynthetic process | Phenazine biosynthesis, Quorum sensing | -25.78680 | NA |  |
| PA14\_09520 | mexI | RND efflux transporter | membrane, transmembrane transporter activity, transmembrane transport | Quorum sensing | -26.04972 | repression |  |
| PA14\_09600 | ddlA | D-alanine--D-alanine ligase | cytoplasm, D-alanine-D-alanine ligase activity, ATP binding, metal ion binding | D-Alanine metabolism, Metabolic pathways, Peptidoglycan biosynthesis, Vancomycin resistance | -26.16340 | repression |  |
| PA14\_09690 | NA | two-component response regulator | regulation of transcription, DNA-templated, phosphorelay signal transduction system, DNA binding |  | -26.03051 | repression |  |
| PA14\_09700 | NA | monooxygenase | FAD binding |  | -25.78646 | repression |  |
| PA14\_09710 | NA | aldehyde dehydrogenase | oxidoreductase activity, oxidoreductase activity, acting on the aldehyde or oxo group of donors, NAD or NADP as acceptor, oxidation-reduction process | Arginine and proline metabolism, Ethanol utilization; second step, Metabolic pathways | -26.08322 | repression |  |
| PA14\_09770 | souR | sarcosine oxidation and utilization regulator SouR | DNA-binding transcription factor activity, regulation of transcription, DNA-templated, sequence-specific DNA binding, DNA binding |  | -27.54743 | no impact |  |
| PA14\_09820 | NA | acetolactate synthase | catalytic activity, thiamine pyrophosphate binding, magnesium ion binding | 2-Oxocarboxylic acid metabolism, Biosynthesis of amino acids, Biosynthesis of antibiotics, Biosynthesis of secondary metabolites, Butanoate metabolism, C5-Branched dibasic acid metabolism, Metabolic pathways, Pantothenate and CoA biosynthesis, Valine, leucine and isoleucine biosynthesis | -26.65378 | repression |  |
| PA14\_09920 | NA | translation initiation inhibitor |  |  | -26.04459 | repression |  |
| PA14\_09950 | NA | oxidoreductase | oxidoreductase activity, oxidation-reduction process |  | -26.01042 | repression |  |
| PA14\_10200 | NA | TonB-dependent receptor protein |  |  | -25.78131 | repression |  |
| PA14\_10240 | NA | branched-chain alpha-keto acid dehydrogenase subunit E2 |  |  | -26.49139 | no impact |  |
| PA14\_10360 | NA | hypothetical protein |  |  | -26.59829 | repression |  |
| PA14\_10370 | NA | hypothetical protein | oxidoreductase activity, flavin adenine dinucleotide binding, oxidation-reduction process, FAD binding, catalytic activity | Bile acid biosynthesis, Zeatin biosynthesis | -26.67096 | repression |  |
| PA14\_10490 | NA | hypothetical protein |  |  | -25.92770 | repression |  |
| PA14\_10500 | NA | cbb3-type cytochrome c oxidase subunit I | cytochrome-c oxidase activity, oxidation-reduction process, aerobic respiration, integral component of membrane, heme binding, plasma membrane respiratory chain complex IV | Metabolic pathways, Oxidative phosphorylation, Two-component system | -26.32564 | no impact |  |
| PA14\_10660 | NA | transcriptional regulator | regulation of transcription, DNA-templated, DNA-binding transcription factor activity, sequence-specific DNA binding, DNA binding |  | -26.16556 | repression |  |
| PA14\_10670 | aph | aminoglycoside 3'-phosphotransferase type IIB | ATP binding, phosphotransferase activity, alcohol group as acceptor, response to antibiotic |  | -26.00929 | no impact |  |
| PA14\_10840 | NA | dehydrogenase |  |  | -26.20237 | repression |  |
| PA14\_11050 | NA | hypothetical protein |  |  | -26.15485 | repression |  |
| PA14\_11060 | cupB1 | fimbrial subunit CupB1 | cell adhesion, pilus |  | -26.02667 | repression |  |
| PA14\_11080 | cupB3 | usher CupB3 | protein binding, pilus assembly, fimbrial usher porin activity, membrane |  | -25.96571 | repression |  |
| PA14\_11100 | cupB5 | adhesive protein CupB5 |  |  | -26.27332 | repression |  |
| PA14\_11120 | NA | response regulator | phosphorelay signal transduction system, regulation of transcription, DNA-templated, DNA binding | Two-component system | -26.17816 | repression |  |
| PA14\_11150 | NA | transcriptional regulator | DNA binding |  | -25.95858 | repression |  |
| PA14\_11170 | NA | hypothetical protein |  |  | -26.64455 | repression |  |
| PA14\_11180 | NA | transcriptional regulator | regulation of transcription, DNA-templated, DNA binding |  | -26.11965 | repression |  |
| PA14\_11230 | NA | hypothetical protein |  |  | -26.66498 | repression |  |
| PA14\_11420 | ribB | bifunctional 3,4-dihydroxy-2-butanone 4-phosphate synthase/GTP cyclohydrolase II-like protein | 3,4-dihydroxy-2-butanone-4-phosphate synthase activity, riboflavin biosynthetic process | Biosynthesis of secondary metabolites, Metabolic pathways, Riboflavin metabolism | -26.13107 | no impact |  |
| PA14\_11680 | NA | two-component regulator | phosphorelay signal transduction system, DNA binding, regulation of transcription, DNA-templated |  | -26.17867 | repression |  |
| PA14\_11845 | mpl | UDP-N-acetylmuramate:L-alanyl-gamma-D-glutamyl- meso-diaminopimelate ligase | ATP binding, biosynthetic process, ligase activity, peptidoglycan biosynthetic process, acid-amino acid ligase activity, cell wall organization, cellular response to antibiotic |  | -25.97372 | no impact |  |
| PA14\_12100 | dacC | D-ala-D-ala-carboxypeptidase | carboxypeptidase activity, proteolysis, serine-type D-Ala-D-Ala carboxypeptidase activity | Metabolic pathways, Peptidoglycan biosynthesis | -26.27169 | repression |  |
| PA14\_12130 | lis | lipoyl synthase | catalytic activity, iron-sulfur cluster binding, lipoate biosynthetic process, lipoate synthase activity, 4 iron, 4 sulfur cluster binding | Lipoic acid metabolism, Metabolic pathways | -25.92238 | no impact |  |
| PA14\_12230 | leuS | leucyl-tRNA synthetase | nucleotide binding, aminoacyl-tRNA ligase activity, ATP binding, tRNA aminoacylation for protein translation, leucine-tRNA ligase activity, leucyl-tRNA aminoacylation, aminoacyl-tRNA editing activity | Aminoacyl-tRNA biosynthesis, Aminoacyl-tRNA biosynthesis | -25.97629 | no impact |  |
| PA14\_12260 | NA | hypothetical protein |  |  | -25.85673 | no impact |  |
| PA14\_12300 | NA | hypothetical protein | flavin adenine dinucleotide binding |  | -26.32822 | repression |  |
| PA14\_12310 | NA | metalloprotease | metalloendopeptidase activity, rRNA processing |  | -26.38705 | repression |  |
| PA14\_12550 | NA | hypothetical protein | oxygen binding, heme binding |  | -25.97154 | repression |  |
| PA14\_12730 | NA | hypothetical protein | catalytic activity |  | -27.86752 | repression |  |
| PA14\_12780 | NA | two-component response regulator | regulation of transcription, DNA-templated, phosphorelay signal transduction system, DNA binding | Two-component system | -26.06717 | repression |  |
| PA14\_12810 | NA | two-component response regulator | phosphorelay signal transduction system |  | -26.85600 | repression |  |
| PA14\_12900 | NA | DNA binding protein | DNA binding |  | -26.06832 | repression |  |
| PA14\_13040 | cioB | CioB, cyanide insensitive terminal oxidase | membrane, oxidation-reduction process | Metabolic pathways, Oxidative phosphorylation, Two-component system | -26.15647 | repression |  |
| PA14\_13060 | NA | transcriptional regulator | DNA-binding transcription factor activity, regulation of transcription, DNA-templated, sequence-specific DNA binding, DNA binding |  | -26.33042 | repression |  |
| PA14\_13190 | NA | hypothetical protein | ATP binding |  | -26.04653 | repression |  |
| PA14\_13240 | moaD | molybdopterin converting factor, small subunit |  |  | -25.85719 | repression |  |
| PA14\_13320 | NA | hypothetical protein |  |  | -25.85143 | repression |  |
| PA14\_13350 | NA | hypothetical protein |  |  | -26.85535 | repression |  |
| PA14\_13360 | NA | hypothetical protein |  |  | -26.12598 | repression |  |
| PA14\_13370 | NA | hypothetical protein |  |  | -26.19700 | no impact |  |
| PA14\_13490 | NA | hypothetical protein | membrane, integral component of membrane |  | -25.82253 | repression |  |
| PA14\_13580 | NA | ABC transporter ATP-binding protein | ATP binding, membrane, glycine betaine transport, ATPase activity | ABC transporters | -26.13390 | repression |  |
| PA14\_13590 | NA | ABC transporter permease | membrane, transmembrane transport | ABC transporters | -25.95618 | repression |  |
| PA14\_13620 | nhaP | Na+/H+ antiporter NhaP | cation transport, solute:proton antiporter activity, integral component of membrane, transmembrane transport |  | -25.90977 | repression |  |
| PA14\_13690 | NA | methyltransferase | methyltransferase activity |  | -26.80318 | repression |  |
| PA14\_13850 | moaA | molybdenum cofactor biosynthesis protein A | catalytic activity, iron-sulfur cluster binding, Mo-molybdopterin cofactor biosynthetic process, metal ion binding, molybdopterin synthase complex, 4 iron, 4 sulfur cluster binding | Folate biosynthesis, Metabolic pathways, Sulfur relay system | -26.00217 | repression |  |
| PA14\_13870 | NA | hypothetical protein |  |  | -26.10598 | repression |  |
| PA14\_14100 | NA | amino acid-binding protein |  |  | -27.06885 | repression |  |
| PA14\_14520 | NA | hypothetical protein |  |  | -26.15200 | repression |  |
| PA14\_14650 | secF | preprotein translocase subunit SecF | intracellular protein transport, P-P-bond-hydrolysis-driven protein transmembrane transporter activity | Bacterial secretion system, Protein export | -25.82436 | repression |  |
| PA14\_14750 | NA | iron-binding protein IscA | structural molecule activity, iron-sulfur cluster binding, protein maturation by iron-sulfur cluster transfer, iron ion binding |  | -26.58702 | no impact |  |
| PA14\_14810 | NA | hypothetical protein | iron-sulfur cluster assembly |  | -25.97687 | repression |  |
| PA14\_14910 | NA | hypothetical protein | protein binding |  | -26.73598 | repression |  |
| PA14\_14975 | NA | hypothetical protein |  |  | -26.59725 | repression |  |
| PA14\_15070 | oprC | outer membrane copper receptor OprC |  |  | -26.21452 | repression |  |
| PA14\_15090 | NA | hypothetical protein |  |  | -26.43017 | repression |  |
| PA14\_15120 | NA | hypothetical protein |  |  | -25.97408 | repression |  |
| PA14\_15290 | NA | transcriptional regulator | DNA binding, regulation of transcription, DNA-templated |  | -27.18934 | repression |  |
| PA14\_15430 | NA | resolvase, essential for transposition | recombinase activity, DNA binding, DNA recombination |  | -25.95126 | repression |  |
| PA14\_15930 | NA | hemolysin | flavin adenine dinucleotide binding |  | -25.90461 | repression |  |
| PA14\_15990 | trmD | tRNA (guanine-N(1)-)-methyltransferase | tRNA processing, tRNA (guanine(37)-N(1))-methyltransferase activity |  | -26.28788 | repression |  |
| PA14\_16030 | NA | sodium/hydrogen antiporter | cation transport, solute:proton antiporter activity, integral component of membrane, transmembrane transport, potassium ion transport |  | -26.51285 | repression |  |
| PA14\_16130 | NA | hypothetical protein | CoA-transferase activity |  | -26.82606 | no impact |  |
| PA14\_16140 | NA | hypothetical protein |  |  | -27.32750 | repression |  |
| PA14\_16150 | NA | hypothetical protein |  |  | -26.41031 | repression |  |
| PA14\_16160 | NA | hypothetical protein |  |  | -27.14996 | repression |  |
| PA14\_16180 | NA | hypothetical protein |  |  | -26.45421 | repression |  |
| PA14\_16200 | NA | hypothetical protein |  |  | -26.89547 | repression |  |
| PA14\_16250 | lasB | elastase LasB | metalloendopeptidase activity, proteolysis, elastin biosynthetic process, elastin catabolic process | Cationic antimicrobial peptide (CAMP) resistance, Quorum sensing | -26.70078 | repression |  |
| PA14\_16320 | NA | peptidyl-prolyl cis-trans isomerase, FkbP-type | peptidyl-prolyl cis-trans isomerase activity |  | -26.42820 | no impact |  |
| PA14\_16410 | NA | MFS transporter | integral component of membrane, transmembrane transporter activity, transmembrane transport |  | -26.17919 | repression |  |
| PA14\_16450 | wspC | methyltransferase | protein binding, S-adenosylmethionine-dependent methyltransferase activity | Two-component system | -26.63684 | no impact |  |
| PA14\_16560 | NA | lipoprotein |  |  | -25.85995 | repression |  |
| PA14\_16600 | NA | alpha/beta hydrolase | proteolysis, peptidase activity |  | -25.81383 | repression |  |
| PA14\_16620 | NA | hypothetical protein |  |  | -26.12588 | repression |  |
| PA14\_16630 | NA | outer membrane protein, OmpA | cell outer membrane, integral component of membrane |  | -25.81715 | repression |  |
| PA14\_16640 | NA | lipoprotein |  |  | -26.92431 | repression |  |
| PA14\_16660 | NA | metal-transporting P-type ATPase | metal ion transport, metal ion binding, cation transport, integral component of membrane, ATPase-coupled cation transmembrane transporter activity, nucleotide binding |  | -25.80573 | no impact |  |
| PA14\_16780 | NA | hypothetical protein | iron ion binding, oxidoreductase activity, acting on paired donors, with incorporation or reduction of molecular oxygen, heme binding, oxidation-reduction process |  | -26.27789 | repression |  |
| PA14\_16820 | NA | efflux transmembrane protein | membrane, transmembrane transporter activity, transmembrane transport |  | -25.79636 | repression |  |
| PA14\_16870 | NA | ABC transporter ATP-binding protein | ATP binding, ATPase activity |  | -26.23051 | repression |  |
| PA14\_17000 | NA | hypothetical protein |  |  | -27.08974 | repression |  |
| PA14\_17010 | NA | Na(+)/H(+) exchanger protein | cation transport, solute:proton antiporter activity, integral component of membrane, transmembrane transport |  | -26.81511 | no impact |  |
| PA14\_17040 | glnD | PII uridylyl-transferase | nitrogen compound metabolic process, [protein-PII] uridylyltransferase activity, nucleotidyltransferase activity | Two-component system | -25.89944 | repression |  |
| PA14\_17060 | rpsB | 30S ribosomal protein S2 | structural constituent of ribosome, ribosome, translation, small ribosomal subunit | Ribosome | -25.95631 | no impact |  |
| PA14\_17100 | frr | ribosome recycling factor | translation |  | -26.66028 | no impact |  |
| PA14\_17110 | uppS | UDP pyrophosphate synthetase | transferase activity, transferring alkyl or aryl (other than methyl) groups | Biosynthesis of secondary metabolites, Terpenoid backbone biosynthesis, Terpenoid biosynthesis | -26.95000 | repression |  |
| PA14\_17290 | pyrG | CTP synthetase | CTP synthase activity, CTP biosynthetic process, pyrimidine nucleotide biosynthetic process | Metabolic pathways, Pyrimidine metabolism | -26.11832 | repression |  |
| PA14\_17380 | gfnR | glutathione-dependent formaldehyde neutralization regulator GfnR | DNA-binding transcription factor activity, regulation of transcription, DNA-templated |  | -25.92163 | no impact |  |
| PA14\_17470 | NA | hypothetical protein |  |  | -26.90654 | repression |  |
| PA14\_17530 | recA | recombinase A | DNA binding, ATP binding, DNA metabolic process, DNA-dependent ATPase activity, single-stranded DNA binding, DNA repair | Homologous recombination | -26.01173 | repression |  |
| PA14\_17550 | NA | hypothetical protein |  |  | -26.52388 | no impact |  |
| PA14\_17570 | NA | hypothetical protein |  |  | -26.19875 | repression |  |
| PA14\_17620 | potC | polyamine transport protein PotC | membrane, transmembrane transport | ABC transporters | -25.91694 | repression |  |
| PA14\_17700 | rpmE2 | 50S ribosomal protein L31 | structural constituent of ribosome, ribosome, translation | Ribosome | -26.80382 | repression |  |
| PA14\_17780 | NA | major facilitator transporter | integral component of membrane, transmembrane transporter activity, transmembrane transport |  | -25.86545 | repression |  |
| PA14\_17850 | NA | enoyl-CoA hydratase | catalytic activity |  | -25.96400 | repression |  |
| PA14\_17860 | NA | 3-hydroxyacyl-CoA dehydrogenase | oxidation-reduction process, 3-hydroxyacyl-CoA dehydrogenase activity, fatty acid metabolic process, oxidoreductase activity | (4Z,7Z,10Z,13Z,16Z)-docosapentaenoate biosynthesis (6-desaturase), (8<i>E</i>,10<i>E</i>)-dodeca-8,10-dienol biosynthesis, (R)- and (S)-3-hydroxybutanoate biosynthesis (engineered), 2-methylpropene degradation, 3-hydroxypropanoate/4-hydroxybutanate cycle, 4-coumarate degradation (aerobic), 4-coumarate degradation (anaerobic), 4-hydroxybenzoate biosynthesis III (plants), <i>Spodoptera littoralis</i> pheromone biosynthesis, alpha-Linolenic acid metabolism, Aminobenzoate degradation, androstenedione degradation, Benzoate degradation, Benzoate degradation, benzoyl-CoA degradation I (aerobic), beta-Alanine metabolism, Butanoate metabolism, Butanoate metabolism, Caprolactam degradation, Carbon fixation pathways in prokaryotes, cholesterol degradation to androstenedione I (cholesterol oxidase), cholesterol degradation to androstenedione II (cholesterol dehydrogenase), crotonate fermentation (to acetate and cyclohexane carboxylate), docosahexaenoate biosynthesis III (6-desaturase, mammals), fatty acid &beta;-oxidation II (peroxisome), Fatty acid degradation, Fatty acid elongation, fatty acid salvage, fermentation to 2-methylbutanoate, Geraniol degradation, glutaryl-CoA degradation, jasmonic acid biosynthesis, Limonene and pinene degradation, Lysine degradation, Metabolic pathways, methyl <i>tert</i>-butyl ether degradation, methyl ketone biosynthesis (engineered), Microbial metabolism in diverse environments, Phenylacetic acid aerobic catabolism, Phenylalanine metabolism, Phenylalanine metabolism, Propanoate metabolism, pyruvate fermentation to butanol I, pyruvate fermentation to butanol II (engineered), pyruvate fermentation to hexanol (engineered), Toluene degradation, Tryptophan metabolism, unsaturated, even numbered fatty acid &beta;-oxidation, Valine, leucine and isoleucine degradation | -25.84608 | repression |  |
| PA14\_17920 | glpM | membrane protein GlpM |  |  | -26.03977 | no impact |  |
| PA14\_18040 | NA | hypothetical protein |  |  | -26.50649 | no impact |  |
| PA14\_18140 | mmsB | 3-hydroxyisobutyrate dehydrogenase | oxidation-reduction process, oxidoreductase activity, NADP binding, NAD binding, 3-hydroxyisobutyrate dehydrogenase activity | Metabolic pathways, Valine, leucine and isoleucine degradation, Valine, leucine and isoleucine degradation | -26.46242 | repression |  |
| PA14\_18200 | NA | LysR family transcriptional regulator | DNA-binding transcription factor activity, regulation of transcription, DNA-templated |  | -27.15498 | repression |  |
| PA14\_18250 | NA | phosphotransferase system enzyme I | phosphoenolpyruvate-dependent sugar phosphotransferase system, transferase activity, transferring phosphorus-containing groups, phosphorylation, catalytic activity | Fructose and mannose metabolism, Metabolic pathways, Microbial metabolism in diverse environments, Phosphotransferase system (PTS) | -26.25190 | no impact |  |
| PA14\_18450 | algI | alginate o-acetyltransferase AlgI | alginic acid biosynthetic process |  | -25.90706 | no impact |  |
| PA14\_18565 | alg8 | alginate biosynthesis protein Alg8 | alginic acid biosynthetic process | Alginate biosynthesis, Fructose and mannose metabolism | -25.79661 | repression |  |
| PA14\_18580 | algD | GDP-mannose 6-dehydrogenase AlgD | oxidation-reduction process, oxidoreductase activity, acting on the CH-OH group of donors, NAD or NADP as acceptor, alginic acid biosynthetic process, NAD binding, GDP-mannose 6-dehydrogenase activity, biofilm formation | Amino sugar and nucleotide sugar metabolism, Fructose and mannose metabolism, Two-component system | -26.19408 | repression |  |
| PA14\_18620 | NA | hypothetical protein |  |  | -26.83607 | repression |  |
| PA14\_18760 | NA | RND efflux membrane fusion protein | membrane, transmembrane transporter activity, transmembrane transport |  | -26.27402 | repression |  |
| PA14\_18830 | NA | adenylosuccinate lyase | N6-(1,2-dicarboxyethyl)AMP AMP-lyase (fumarate-forming) activity, purine ribonucleotide biosynthetic process, catalytic activity | adenosine ribonucleotides <i>de novo</i> biosynthesis, Alanine, aspartate and glutamate metabolism, Alanine, aspartate and glutamate metabolism, Biosynthesis of antibiotics, Biosynthesis of secondary metabolites, inosine-5'-phosphate biosynthesis I, inosine-5'-phosphate biosynthesis II, inosine-5'-phosphate biosynthesis III, Metabolic pathways, Purine metabolism, Purine metabolism | -26.29362 | no impact |  |
| PA14\_18850 | NA | adenylosuccinate lyase | catalytic activity | adenosine ribonucleotides <i>de novo</i> biosynthesis, Alanine, aspartate and glutamate metabolism, Alanine, aspartate and glutamate metabolism, Biosynthesis of antibiotics, Biosynthesis of secondary metabolites, inosine-5'-phosphate biosynthesis I, inosine-5'-phosphate biosynthesis II, inosine-5'-phosphate biosynthesis III, Metabolic pathways, Purine metabolism, Purine metabolism | -27.01238 | repression |  |
| PA14\_18860 | NA | hypothetical protein |  |  | -25.80204 | repression |  |
| PA14\_18960 | tli5a | type VI secretion lipase immunity Tli5A |  |  | -27.15836 | repression |  |
| PA14\_18970 | pldA | phospholipase D | catalytic activity |  | -25.89773 | repression |  |
| PA14\_18985 | vgrG4b | VgrG4b |  |  | -25.94737 | repression |  |
| PA14\_19020 | tse3 | Tse3 |  |  | -26.51496 | repression |  |
| PA14\_19030 | NA | hypothetical protein |  |  | -26.85621 | repression |  |
| PA14\_19130 | rhlI | autoinducer synthesis protein RhlI | transferase activity | autoinducer AI-1 biosynthesis, Biofilm formation - Pseudomonas aeruginosa, Cysteine and methionine metabolism, Cysteine and methionine metabolism, Metabolic pathways, Quorum sensing, Two-component system | -26.37784 | repression |  |
| PA14\_19170 | NA | hypothetical protein |  |  | -25.84970 | no impact |  |
| PA14\_19230 | NA | hypothetical protein | membrane, transmembrane transport |  | -25.87870 | repression |  |
| PA14\_19360 | NA | GNAT family acetyltransferase | ATP binding, metal ion binding, N-acetyltransferase activity |  | -26.85104 | repression |  |
| PA14\_19370 | NA | asparagine synthetase | asparagine synthase (glutamine-hydrolyzing) activity, asparagine biosynthetic process | Alanine, aspartate and glutamate metabolism, Biosynthesis of secondary metabolites, Metabolic pathways | -26.61743 | repression |  |
| PA14\_19380 | NA | transcriptional regulator | DNA-binding transcription factor activity, regulation of transcription, DNA-templated |  | -27.06465 | repression |  |
| PA14\_19490 | lsfA | 1-Cys peroxiredoxin LsfA | oxidation-reduction process, cell redox homeostasis, antioxidant activity, oxidoreductase activity, peroxiredoxin activity, response to hydrogen peroxide, pathogenesis |  | -25.81317 | no impact |  |
| PA14\_19590 | NA | molybdopterin-binding protein | molybdate ion transport |  | -26.70072 | no impact |  |
| PA14\_19620 | folX | D-erythro-7,8-dihydroneopterin triphosphate 2'-epimerase | dihydroneopterin aldolase activity, folic acid-containing compound metabolic process, tetrahydrobiopterin biosynthetic process |  | -26.43002 | no impact |  |
| PA14\_19670 | NA | LysR family transcriptional regulator | DNA-binding transcription factor activity, regulation of transcription, DNA-templated |  | -27.29182 | repression |  |
| PA14\_19720 | NA | hypothetical protein |  |  | -26.26229 | repression |  |
| PA14\_19860 | NA | hypothetical protein |  |  | -26.01294 | repression |  |
| PA14\_19870 | ldh | leucine dehydrogenase | cellular amino acid metabolic process, oxidoreductase activity, oxidation-reduction process, oxidoreductase activity, acting on the CH-NH2 group of donors, NAD or NADP as acceptor | Biosynthesis of antibiotics, Biosynthesis of secondary metabolites, Metabolic pathways, Valine, leucine and isoleucine biosynthesis, Valine, leucine and isoleucine degradation | -25.99024 | no impact |  |
| PA14\_19910 | NA | pyruvate dehydrogenase E1 component, beta chain | catalytic activity | Biosynthesis of antibiotics, Biosynthesis of secondary metabolites, Carbon metabolism, Citrate cycle (TCA cycle), Glycolysis / Gluconeogenesis, Metabolic pathways, Microbial metabolism in diverse environments, Pyruvate metabolism | -26.16541 | repression |  |
| PA14\_19950 | NA | hypothetical protein |  |  | -25.82443 | no impact |  |
| PA14\_20010 | hasR | heme uptake outer membrane receptor HasR | outer membrane, transmembrane transporter activity, transmembrane transport, heme transporter activity, heme transport, heme binding |  | -26.86567 | no impact |  |
| PA14\_20120 | NA | hypothetical protein |  |  | -25.76839 | repression |  |
| PA14\_20130 | NA | LysR family transcriptional regulator | DNA-binding transcription factor activity, regulation of transcription, DNA-templated |  | -26.42456 | repression |  |
| PA14\_20190 | nosD | copper ABC transporter periplasmic substrate-binding protein |  |  | -26.16724 | repression |  |
| PA14\_20280 | NA | hypothetical protein |  |  | -25.88817 | no impact |  |
| PA14\_20440 | phnN | phosphonate transport ATP-binding protein | 5-phosphoribose 1-diphosphate biosynthetic process, ribose 1,5-bisphosphate phosphokinase activity | Pentose phosphate pathway | -26.32165 | no impact |  |
| PA14\_20460 | NA | hypothetical protein |  |  | -26.73825 | repression |  |
| PA14\_20470 | NA | hypothetical protein |  |  | -26.80276 | repression |  |
| PA14\_20560 | amiE | acylamide amidohydrolase | nitrogen compound metabolic process, amidase activity | Aminobenzoate degradation, Arginine and proline metabolism, Microbial metabolism in diverse environments, Phenylalanine metabolism, Styrene degradation, Tryptophan metabolism | -26.36225 | repression |  |
| PA14\_20580 | amiC | aliphatic amidase expression-regulating protein | amino acid transport, negative regulation of hydrolase activity, amide binding, regulation of cellular amide catabolic process | ABC transporters, Quorum sensing | -26.62535 | repression |  |
| PA14\_20670 | NA | glutamine synthetase | catalytic activity, glutamate-ammonia ligase activity, nitrogen compound metabolic process | Alanine, aspartate and glutamate metabolism, Alanine, aspartate and glutamate metabolism, ammonia assimilation cycle I, ammonia assimilation cycle II, Arginine biosynthesis, Arginine biosynthesis, Biosynthesis of amino acids, Glyoxylate and dicarboxylate metabolism, Glyoxylate and dicarboxylate metabolism, L-glutamine biosynthesis III, Metabolic pathways, Microbial metabolism in diverse environments, nitrate reduction II (assimilatory), nitrate reduction V (assimilatory), Nitrogen metabolism, Nitrogen metabolism, Two-component system | -25.86110 | repression |  |
| PA14\_20770 | NA | hypothetical protein |  |  | -25.93219 | no impact |  |
| PA14\_20800 | NA | histidine phosphotransfer domain-containing protein | phosphorelay signal transduction system, histidine phosphotransfer kinase activity, negative regulation of single-species biofilm formation, positive regulation of chemotaxis, regulation of cell motility | Biofilm formation - Pseudomonas aeruginosa, Two-component system | -26.42412 | repression |  |
| PA14\_20940 | NA | acyl carrier protein |  |  | -27.31959 | repression |  |
| PA14\_21020 | NA | non-ribosomal peptide synthetase | biosynthetic process, hydrolase activity, acting on ester bonds, phosphopantetheine binding, catalytic activity |  | -26.91402 | repression |  |
| PA14\_21030 | NA | ATP-dependent Clp protease proteolytic subunit | serine-type endopeptidase activity, proteolysis |  | -27.05512 | repression |  |
| PA14\_21210 | NA | hypothetical protein | integral component of membrane, catalytic activity, sulfuric ester hydrolase activity | Cationic antimicrobial peptide (CAMP) resistance | -26.12260 | repression |  |
| PA14\_21300 | NA | MFS transporte | integral component of plasma membrane, transmembrane transport |  | -25.82375 | repression |  |
| PA14\_21310 | phaJ1 | hypothetical protein | fatty acid synthase activity, fatty acid synthase complex, fatty acid biosynthetic process, oxidation-reduction process |  | -25.91942 | repression |  |
| PA14\_21450 | NA | hypothetical protein |  |  | -25.94737 | repression |  |
| PA14\_21530 | NA | ankyrin domain-containing protein | protein binding |  | -25.79932 | repression |  |
| PA14\_21580 | NA | hypothetical protein |  |  | -26.59585 | repression |  |
| PA14\_21600 | NA | hypothetical protein |  |  | -25.94506 | repression |  |
| PA14\_21610 | oprO | pyrophosphate-specific outer membrane porin OprO precursor |  |  | -26.83569 | repression |  |
| PA14\_21620 | oprP | phosphate-specific outer membrane porin OprP precursor |  |  | -26.48652 | repression |  |
| PA14\_21630 | NA | hypothetical protein |  |  | -26.15904 | repression |  |
| PA14\_21670 | NA | hypothetical protein |  |  | -25.88098 | no impact |  |
| PA14\_21900 | NA | HAD-superfamily hydrolase |  |  | -26.37002 | no impact |  |
| PA14\_21960 | NA | hypothetical protein |  |  | -25.82256 | repression |  |
| PA14\_21970 | NA | transcriptional regulator | DNA binding, regulation of transcription, DNA-templated, DNA-binding transcription factor activity |  | -26.32323 | repression |  |
| PA14\_21980 | NA | hypothetical protein | N-acetyltransferase activity |  | -25.82838 | repression |  |
| PA14\_22320 | NA | hypothetical protein |  |  | -26.31297 | repression |  |
| PA14\_22330 | NA | glycine betaine-binding protein | transmembrane transporter activity, ATP-binding cassette (ABC) transporter complex, transmembrane transport | ABC transporters | -26.57926 | repression |  |
| PA14\_22420 | NA | hypothetical protein |  |  | -26.74312 | repression |  |
| PA14\_22440 | NA | ABC transporter ATP-binding protein/permease | ATP binding, ATPase activity, integral component of membrane, ATPase-coupled transmembrane transporter activity, transmembrane transport | ABC transporters | -25.98806 | repression |  |
| PA14\_22470 | NA | LysR family transcriptional regulator | DNA-binding transcription factor activity, regulation of transcription, DNA-templated |  | -26.47356 | repression |  |
| PA14\_22570 | csaA | CsaA protein | tRNA binding |  | -26.01327 | repression |  |
| PA14\_22650 | NA | ABC transporter |  |  | -25.78261 | repression |  |
| PA14\_22710 | NA | hypothetical protein | oxidoreductase activity |  | -25.85693 | repression |  |
| PA14\_23050 | NA | hypothetical protein | catalytic activity, carbohydrate metabolic process, carbohydrate binding, isomerase activity |  | -26.04791 | repression |  |
| PA14\_23080 | pgl | 6-phosphogluconolactonase | carbohydrate metabolic process, pentose-phosphate shunt, 6-phosphogluconolactonase activity | Amino sugar and nucleotide sugar metabolism, Biosynthesis of antibiotics, Biosynthesis of secondary metabolites, Carbon metabolism, chitin degradation I (archaea), chitin derivatives degradation, Entner-Doudoroff pathway I, Metabolic pathways, Microbial metabolism in diverse environments, Pentose phosphate pathway, Pentose phosphate pathway, UDP-<i>N</i>-acetyl-D-galactosamine biosynthesis II | -25.86417 | repression |  |
| PA14\_23120 | NA | hypothetical protein |  |  | -26.39796 | no impact |  |
| PA14\_23260 | gyrA | DNA gyrase subunit A | DNA binding, DNA topoisomerase type II (ATP-hydrolyzing) activity, ATP binding, DNA topological change, chromosome, DNA topoisomerase activity, DNA metabolic process |  | -26.47014 | repression |  |
| PA14\_23280 | pheA | chorismate mutase | prephenate dehydratase activity, L-phenylalanine biosynthetic process, chorismate metabolic process, chorismate mutase activity, cytoplasm | Biosynthesis of amino acids, Biosynthesis of antibiotics, Biosynthesis of secondary metabolites, Metabolic pathways, Phenylalanine, tyrosine and tryptophan biosynthesis | -25.96976 | repression |  |
| PA14\_23340 | ihfB | integration host factor subunit beta | DNA binding, chromosome, DNA recombination, regulation of transcription, DNA-templated |  | -27.11791 | no impact |  |
| PA14\_23370 | orfK | UDP-N-acetylglucosamine 2-epimerase | UDP-N-acetylglucosamine 2-epimerase activity | Amino sugar and nucleotide sugar metabolism, Metabolic pathways | -27.07920 | no impact |  |
| PA14\_23450 | orfM | NAD dependent epimerase/dehydratase | catalytic activity, coenzyme binding |  | -25.88524 | no impact |  |
| PA14\_23470 | wbpM | nucleotide sugar epimerase/dehydratase WbpM |  |  | -27.51562 | repression |  |
| PA14\_23540 | act | transcriptional regulator | DNA-binding transcription factor activity, regulation of transcription, DNA-templated |  | -25.89344 | repression |  |
| PA14\_23630 | NA | hypothetical protein |  |  | -26.12185 | no impact |  |
| PA14\_23700 | NA | LysR family transcriptional regulator | DNA-binding transcription factor activity, regulation of transcription, DNA-templated |  | -26.38005 | repression |  |
| PA14\_23830 | fimV | pilus assembly protein | protein binding |  | -26.57988 | repression |  |
| PA14\_23890 | NA | hypothetical protein | peptidoglycan binding |  | -26.78843 | repression |  |
| PA14\_23920 | purF | amidophosphoribosyltransferase | amidophosphoribosyltransferase activity, purine nucleobase biosynthetic process, nucleoside metabolic process | Alanine, aspartate and glutamate metabolism, Biosynthesis of antibiotics, Biosynthesis of secondary metabolites, Metabolic pathways, Purine metabolism | -26.02310 | repression |  |
| PA14\_23930 | metZ | O-succinylhomoserine sulfhydrylase | transsulfuration, pyridoxal phosphate binding, homocysteine biosynthetic process, catalytic activity, methionine biosynthetic process | Cysteine and methionine metabolism, Metabolic pathways, Sulfur metabolism | -26.19706 | repression |  |
| PA14\_24040 | xcpU | general secretion pathway outer membrane protein H precursor | protein secretion by the type II secretion system, type II protein secretion system complex | Bacterial secretion system | -25.90969 | repression |  |
| PA14\_24070 | xcpX | general secretion pathway protein K | protein secretion, integral component of membrane, type II protein secretion system complex | Bacterial secretion system | -25.93777 | repression |  |
| PA14\_24210 | NA | hypothetical protein |  |  | -26.41236 | no impact |  |
| PA14\_24440 | NA | lipoprotein |  |  | -26.75498 | repression |  |
| PA14\_24490 | pelB | hypothetical protein | protein binding, single-species biofilm formation, single-species biofilm formation | Biofilm formation - Pseudomonas aeruginosa | -25.84737 | repression |  |
| PA14\_24580 | NA | hypothetical protein | DNA binding |  | -26.52792 | repression |  |
| PA14\_24590 | NA | hypothetical protein | DNA binding |  | -26.60951 | repression |  |
| PA14\_24700 | NA | hypothetical protein |  |  | -25.92705 | repression |  |
| PA14\_24840 | NA | transcriptional regulator | DNA binding |  | -26.79541 | repression |  |
| PA14\_24890 | mobA | molybdopterin-guanine dinucleotide biosynthesis protein MobA | catalytic activity, Mo-molybdopterin cofactor biosynthetic process |  | -26.07043 | no impact |  |
| PA14\_24940 | NA | hypothetical protein | flavin adenine dinucleotide binding, FAD binding, catalytic activity, oxidoreductase activity, oxidation-reduction process | Ether lipid metabolism, Metabolic pathways | -26.44050 | repression |  |
| PA14\_24970 | NA | lipid kinase | lipid kinase activity, metal ion binding, kinase activity, NAD+ kinase activity |  | -27.05651 | repression |  |
| PA14\_25030 | NA | hypothetical protein |  |  | -25.81730 | repression |  |
| PA14\_25040 | NA | hypothetical protein |  |  | -26.24908 | no impact |  |
| PA14\_25060 | NA | hypothetical protein |  |  | -26.56518 | repression |  |
| PA14\_25080 | fadB | multifunctional fatty acid oxidation complex subunit alpha | 3-hydroxyacyl-CoA dehydrogenase activity, dodecenoyl-CoA delta-isomerase activity, enoyl-CoA hydratase activity, 3-hydroxybutyryl-CoA epimerase activity, fatty acid catabolic process, fatty acid beta-oxidation multienzyme complex, oxidation-reduction process, catalytic activity, fatty acid metabolic process, oxidoreductase activity | Benzoate degradation, beta-Alanine metabolism, Biosynthesis of antibiotics, Biosynthesis of secondary metabolites, Biosynthesis of unsaturated fatty acids, Butanoate metabolism, Caprolactam degradation, Carbon metabolism, Fatty acid degradation, Fatty acid metabolism, Geraniol degradation, Limonene and pinene degradation, Lysine degradation, Metabolic pathways, Microbial metabolism in diverse environments, Propanoate metabolism, Tryptophan metabolism, Valine, leucine and isoleucine degradation | -26.35696 | repression |  |
| PA14\_25090 | fadA | 3-ketoacyl-CoA thiolase | transferase activity, transferring acyl groups other than amino-acyl groups, acetyl-CoA C-acyltransferase activity, cytoplasm, fatty acid metabolic process, lipid catabolic process, catalytic activity | (4Z,7Z,10Z,13Z,16Z)-docosapentaenoate biosynthesis (6-desaturase), (8<i>E</i>,10<i>E</i>)-dodeca-8,10-dienol biosynthesis, 10-<i>cis</i>-heptadecenoyl-CoA degradation (yeast), 10-<i>trans</i>-heptadecenoyl-CoA degradation (MFE-dependent, yeast), 10-<i>trans</i>-heptadecenoyl-CoA degradation (reductase-dependent, yeast), 4-ethylphenol degradation (anaerobic), 4-hydroxybenzoate biosynthesis III (plants), 4-oxopentanoate degradation, 9-<i>cis</i>, 11-<i>trans</i>-octadecadienoyl-CoA degradation (isomerase-dependent, yeast), alpha-Linolenic acid metabolism, alpha-Linolenic acid metabolism, androstenedione degradation, Benzoate degradation, Benzoate degradation, Biosynthesis of antibiotics, Biosynthesis of secondary metabolites, cholesterol degradation to androstenedione I (cholesterol oxidase), cholesterol degradation to androstenedione II (cholesterol dehydrogenase), docosahexaenoate biosynthesis III (6-desaturase, mammals), Ethylbenzene degradation, fatty acid &beta;-oxidation (peroxisome, yeast), fatty acid &beta;-oxidation II (peroxisome), Fatty acid degradation, Fatty acid degradation, Fatty acid elongation, Fatty acid metabolism, fatty acid salvage, fermentation to 2-methylbutanoate, Geraniol degradation, Geraniol degradation, jasmonic acid biosynthesis, Metabolic pathways, Microbial metabolism in diverse environments, pyruvate fermentation to hexanol (engineered), sitosterol degradation to androstenedione, Valine, leucine and isoleucine degradation, Valine, leucine and isoleucine degradation | -25.81502 | no impact |  |
| PA14\_25210 | NA | 5'-methylthioadenosine phosphorylase | S-methyl-5-thioadenosine phosphorylase activity, catalytic activity, nucleoside metabolic process, transferase activity, transferring pentosyl groups | Cysteine and methionine metabolism, Metabolic pathways | -25.89917 | no impact |  |
| PA14\_25270 | aroP1 | aromatic amino acid transport protein AroP1 | amino acid transport, integral component of membrane, transmembrane transport, membrane, transmembrane transporter activity |  | -26.17613 | repression |  |
| PA14\_25305 | nqrB | Na(+)-translocating NADH-quinone reductase subunit B | FMN binding, integral component of membrane, oxidoreductase activity, acting on NAD(P)H, quinone or similar compound as acceptor, respiratory electron transport chain, membrane, transmembrane transport |  | -26.26475 | repression |  |
| PA14\_25340 | nqrE | Na(+)-translocating NADH-quinone reductase subunit E | Gram-negative-bacterium-type cell wall, integral component of membrane, oxidoreductase activity, acting on NAD(P)H, quinone or similar compound as acceptor, respiratory electron transport chain, oxidation-reduction process, membrane |  | -26.42902 | repression |  |
| PA14\_25390 | sth | soluble pyridine nucleotide transhydrogenase | cell redox homeostasis, oxidation-reduction process, oxidoreductase activity, flavin adenine dinucleotide binding, electron transfer activity, NAD(P)+ transhydrogenase (B-specific) activity | Metabolic pathways, Nicotinate and nicotinamide metabolism | -26.72353 | no impact |  |
| PA14\_25420 | NA | hypothetical protein | cyclic-di-GMP binding |  | -26.54937 | no impact |  |
| PA14\_25480 | NA | competence protein | integral component of membrane, establishment of competence for transformation |  | -26.63862 | repression |  |
| PA14\_25520 | NA | hypothetical protein |  |  | -25.89708 | repression |  |
| PA14\_25830 | NA | hypothetical protein |  |  | -26.41059 | no impact |  |
| PA14\_25840 | NA | electron transfer flavoprotein-ubiquinone oxidoreductase |  |  | -25.80475 | repression |  |
| PA14\_26000 | NA | magnesium chelatase |  |  | -25.95272 | repression |  |
| PA14\_26140 | NA | transcriptional regulator | DNA binding |  | -27.06976 | no impact |  |
| PA14\_26165 | NA | hypothetical protein |  |  | -27.31012 | no impact |  |
| PA14\_26210 | hisP | histidine transport | ATP binding, ATPase activity, amino acid transmembrane transport, ATPase-coupled amino acid transmembrane transporter activity | ABC transporters | -26.58451 | no impact |  |
| PA14\_26240 | hisJ | periplasmic histidine-binding protein HisJ |  |  | -25.81831 | repression |  |
| PA14\_26510 | cobI | precorrin-2 C(20)-methyltransferase | S-adenosylmethionine-dependent methyltransferase activity, cobalamin biosynthetic process, precorrin-2 C20-methyltransferase activity, methyltransferase activity | Metabolic pathways, Porphyrin and chlorophyll metabolism | -26.13547 | repression |  |
| PA14\_26690 | NA | enoyl-CoA hydratase/isomerase | catalytic activity | Geraniol degradation | -26.08283 | repression |  |
| PA14\_26700 | NA | acyl-CoA dehydrogenase | oxidoreductase activity, acting on the CH-CH group of donors, oxidation-reduction process, flavin adenine dinucleotide binding, acyl-CoA dehydrogenase activity, citronellyl-CoA dehydrogenase activity, terpene catabolic process | Geraniol degradation | -26.38646 | repression |  |
| PA14\_26760 | NA | transcriptional regulator | DNA binding |  | -26.41044 | repression |  |
| PA14\_26850 | NA | hypothetical protein |  |  | -26.42383 | repression |  |
| PA14\_26880 | bvlR | BvlR | DNA-binding transcription factor activity, regulation of transcription, DNA-templated, pathogenesis |  | -27.42778 | repression |  |
| PA14\_26940 | NA | hypothetical protein |  |  | -26.56985 | no impact |  |
| PA14\_27370 | NA | ATP-dependent RNA helicase | nucleic acid binding, ATP binding, ribosomal large subunit assembly, RNA helicase activity, cellular response to cold | RNA degradation | -26.15910 | repression |  |
| PA14\_27400 | NA | LysR family transcriptional regulator | DNA-binding transcription factor activity, regulation of transcription, DNA-templated |  | -26.60955 | repression |  |
| PA14\_27510 | NA | methionine sulfoxide reductase B | peptide-methionine (R)-S-oxide reductase activity, oxidation-reduction process, pathogenesis, response to hypochlorite, cellular response to oxidative stress |  | -26.02742 | repression |  |
| PA14\_27730 | fadE | acyl-CoA dehydrogenase | oxidoreductase activity, acting on the CH-CH group of donors, oxidation-reduction process, flavin adenine dinucleotide binding, acyl-CoA dehydrogenase activity, fatty acid beta-oxidation using acyl-CoA dehydrogenase | Fatty acid degradation, Fatty acid metabolism, Metabolic pathways | -25.89754 | repression |  |
| PA14\_27755 | NA | glutathione S-transferase | protein binding | Glutathione metabolism | -25.75142 | repression |  |
| PA14\_27810 | NA | two-component response regulator | phosphorelay signal transduction system, DNA binding, regulation of transcription, DNA-templated | Two-component system | -25.83603 | repression |  |
| PA14\_27830 | NA | hypothetical protein | negative regulation of protein secretion, negative regulation of transcription, DNA-templated, stress response to copper ion |  | -25.96994 | repression |  |
| PA14\_27910 | NA | hypothetical protein |  |  | -26.02109 | no impact |  |
| PA14\_27990 | NA | sialidase |  |  | -26.40647 | no impact |  |
| PA14\_28010 | NA | hypothetical protein |  |  | -26.54012 | repression |  |
| PA14\_28070 | NA | hypothetical protein | protein binding |  | -26.60333 | repression |  |
| PA14\_28130 | NA | hypothetical protein | DNA binding |  | -26.71865 | repression |  |
| PA14\_28170 | NA | formate/nitrate transporter | membrane, transmembrane transporter activity, transmembrane transport |  | -26.27395 | no impact |  |
| PA14\_28290 | NA | hypothetical protein | hydrolase activity |  | -26.04588 | no impact |  |
| PA14\_28400 | NA | outer membrane OprD family porin | integral component of membrane |  | -26.52561 | no impact |  |
| PA14\_28410 | NA | hypothetical protein |  |  | -26.33321 | repression |  |
| PA14\_28440 | NA | hypothetical protein |  |  | -26.66162 | repression |  |
| PA14\_28450 | eco | ecotin | serine-type endopeptidase inhibitor activity |  | -25.82337 | repression |  |
| PA14\_28530 | NA | hypothetical protein |  |  | -26.32650 | repression |  |
| PA14\_28600 | NA | hypothetical protein |  |  | -25.98767 | repression |  |
| PA14\_28630 | NA | hydrolase |  |  | -26.31442 | repression |  |
| PA14\_28650 | thrS | threonyl-tRNA synthetase | aminoacyl-tRNA ligase activity, ATP binding, tRNA aminoacylation, nucleotide binding, tRNA aminoacylation for protein translation, threonine-tRNA ligase activity, cytoplasm, threonyl-tRNA aminoacylation | Aminoacyl-tRNA biosynthesis | -26.04016 | repression |  |
| PA14\_28660 | infC | translation initiation factor IF-3 | translation initiation factor activity, translational initiation |  | -25.88104 | repression |  |
| PA14\_28670 | rpmI | 50S ribosomal protein L35 | structural constituent of ribosome, ribosome, translation | Ribosome | -27.49188 | no impact |  |
| PA14\_28690 | pheS | phenylalanyl-tRNA synthetase subunit alpha | phenylalanine-tRNA ligase activity, nucleotide binding, tRNA binding, aminoacyl-tRNA ligase activity, ATP binding, tRNA aminoacylation, cytoplasm, phenylalanyl-tRNA aminoacylation | Aminoacyl-tRNA biosynthesis | -25.94277 | repression |  |
| PA14\_28720 | ihfA | integration host factor subunit alpha | DNA binding, DNA recombination, regulation of transcription, DNA-templated |  | -26.96559 | repression |  |
| PA14\_28910 | NA | radical activating enzyme | catalytic activity, iron-sulfur cluster binding |  | -25.89496 | repression |  |
| PA14\_28940 | NA | hypothetical protein |  |  | -26.66373 | repression |  |
| PA14\_28950 | NA | hypothetical protein | carbon-sulfur lyase activity |  | -26.88290 | repression |  |
| PA14\_28990 | NA | hypothetical protein |  |  | -25.92243 | repression |  |
| PA14\_29020 | cpo | chloroperoxidase |  |  | -26.83011 | repression |  |
| PA14\_29060 | NA | transcriptional regulator |  |  | -26.19856 | repression |  |
| PA14\_29150 | NA | hypothetical protein | carbon-sulfur lyase activity | formaldehyde oxidation II (glutathione-dependent), Methane metabolism | -25.79057 | repression |  |
| PA14\_29190 | NA | hypothetical protein |  |  | -25.91037 | no impact |  |
| PA14\_29260 | NA | transcriptional regulator | DNA-binding transcription factor activity, regulation of transcription, DNA-templated, sequence-specific DNA binding, DNA binding |  | -27.72237 | no impact |  |
| PA14\_29270 | NA | outer membrane lipoprotein |  |  | -26.14561 | repression |  |
| PA14\_29360 | pfeS | two-component sensor PfeS | signal transduction, integral component of membrane, phosphorelay sensor kinase activity, phosphorylation, transferase activity, transferring phosphorus-containing groups | Two-component system | -25.90853 | repression |  |
| PA14\_29400 | NA | hypothetical protein |  |  | -26.49574 | repression |  |
| PA14\_29520 | NA | type II secretion system protein |  |  | -26.37774 | repression |  |
| PA14\_29530 | NA | type II secretion system protein |  |  | -26.02191 | repression |  |
| PA14\_29560 | NA | hypothetical protein |  |  | -26.02736 | repression |  |
| PA14\_29570 | NA | hypothetical protein |  |  | -26.53862 | repression |  |
| PA14\_29650 | NA | hypothetical protein |  |  | -26.27926 | repression |  |
| PA14\_29720 | NA | hypothetical protein |  |  | -25.98739 | repression |  |
| PA14\_29730 | bqsR | two-component response regulator BqsR | DNA binding, regulation of transcription, DNA-templated, phosphorelay signal transduction system, cellular response to iron(II) ion |  | -25.93389 | repression |  |
| PA14\_29750 | NA | hypothetical protein |  |  | -26.74093 | repression |  |
| PA14\_29890 | nuoK | NADH dehydrogenase subunit K | oxidoreductase activity, acting on NAD(P)H, ATP synthesis coupled electron transport, oxidation-reduction process | Metabolic pathways, Oxidative phosphorylation | -26.05029 | repression |  |
| PA14\_30020 | nuoA | NADH dehydrogenase subunit A | oxidoreductase activity, acting on NAD(P)H, oxidation-reduction process, NADH dehydrogenase (ubiquinone) activity | Metabolic pathways, Oxidative phosphorylation | -25.78831 | no impact |  |
| PA14\_30190 | icd | isocitrate dehydrogenase | isocitrate dehydrogenase (NADP+) activity, tricarboxylic acid cycle, oxidation-reduction process, oxidoreductase activity, acting on the CH-OH group of donors, NAD or NADP as acceptor, magnesium ion binding, NAD binding | 2-Oxocarboxylic acid metabolism, Biosynthesis of amino acids, Biosynthesis of antibiotics, Biosynthesis of secondary metabolites, Carbon metabolism, Citrate cycle (TCA cycle), Glutathione metabolism, Metabolic pathways, Microbial metabolism in diverse environments | -25.94623 | repression |  |
| PA14\_30200 | cspD | cold-shock protein CspD | nucleic acid binding, cytoplasm, regulation of transcription, DNA-templated |  | -26.62991 | no impact |  |
| PA14\_30210 | clpS | ATP-dependent Clp protease adaptor protein ClpS | protein catabolic process, bacterial-type flagellum-dependent swarming motility, cellular response to antibiotic, single-species biofilm formation, single-species biofilm formation on inanimate substrate |  | -26.30038 | no impact |  |
| PA14\_30230 | clpA | ATP-dependent Clp protease, ATP-binding subunit ClpA | ATP binding, protein metabolic process, ATPase activity, protein unfolding |  | -25.89748 | repression |  |
| PA14\_30240 | infA | translation initiation factor IF-1 | translation initiation factor activity, translational initiation, RNA binding |  | -26.61809 | repression |  |
| PA14\_30340 | cysG | siroheme synthase | methyltransferase activity, oxidation-reduction process, porphyrin-containing compound biosynthetic process, uroporphyrin-III C-methyltransferase activity, cobalamin biosynthetic process, siroheme biosynthetic process, precorrin-2 dehydrogenase activity, sirohydrochlorin ferrochelatase activity, NAD binding | Biosynthesis of secondary metabolites, cob(II)yrinate <i>a,c</i>-diamide biosynthesis I (early cobalt insertion), factor 430 biosynthesis, Metabolic pathways, Porphyrin and chlorophyll metabolism, Porphyrin and chlorophyll metabolism, siroheme biosynthesis | -25.81183 | repression |  |
| PA14\_30350 | NA | hypothetical protein | glutathione transferase activity, protein binding |  | -26.02175 | repression |  |
| PA14\_30370 | NA | hypothetical protein |  |  | -25.82490 | no impact |  |
| PA14\_30560 | NA | hypothetical protein |  |  | -26.43421 | repression |  |
| PA14\_30630 | pqsH | FAD-dependent monooxygenase | FAD binding | Biofilm formation - Pseudomonas aeruginosa, Quorum sensing | -27.04271 | repression |  |
| PA14\_30750 | NA | tryptophan oxygenase | tryptophan 2,3-dioxygenase activity, tryptophan catabolic process to kynurenine, heme binding, metal ion binding | Metabolic pathways, Tryptophan metabolism, Tryptophan metabolism | -26.43842 | no impact |  |
| PA14\_30790 | NA | hypothetical protein | membrane, integral component of membrane |  | -26.53333 | repression |  |
| PA14\_30820 | NA | methyl-accepting chemotaxis transducer | signal transduction, integral component of membrane, membrane | Bacterial chemotaxis, Two-component system | -26.95724 | repression |  |
| PA14\_30840 | NA | signal transduction histidine kinase | phosphorylation, transferase activity, transferring phosphorus-containing groups, phosphorelay sensor kinase activity, signal transduction |  | -26.60344 | no impact |  |
| PA14\_31310 | NA | hypothetical protein |  |  | -26.80463 | repression |  |
| PA14\_31360 | NA | hypothetical protein |  |  | -27.07203 | repression |  |
| PA14\_31390 | NA | hypothetical protein |  |  | -25.85800 | repression |  |
| PA14\_31420 | NA | hypothetical protein |  |  | -26.07096 | repression |  |
| PA14\_31440 | NA | hypothetical protein |  |  | -26.69689 | repression |  |
| PA14\_31460 | NA | transporter | membrane |  | -26.52212 | repression |  |
| PA14\_31580 | NA | acyl-CoA dehydrogenase | oxidoreductase activity, acting on the CH-CH group of donors, oxidation-reduction process, flavin adenine dinucleotide binding | beta-Alanine metabolism, Biosynthesis of antibiotics, Biosynthesis of secondary metabolites, Carbon metabolism, Fatty acid degradation, Fatty acid metabolism, Metabolic pathways, Propanoate metabolism, Valine, leucine and isoleucine degradation | -27.33235 | repression |  |
| PA14\_31720 | NA | hypothetical protein |  |  | -26.93974 | repression |  |
| PA14\_31760 | NA | phosphatidate cytidylyltransferase |  |  | -26.37359 | repression |  |
| PA14\_31800 | NA | sodium:alanine symporter | sodium ion transport, alanine:sodium symporter activity, membrane, alanine transport |  | -26.38902 | repression |  |
| PA14\_31810 | tpx | thiol peroxidase | oxidoreductase activity, oxidoreductase activity, acting on peroxide as acceptor, oxidation-reduction process, cell redox homeostasis, thioredoxin peroxidase activity, antioxidant activity |  | -26.16785 | repression |  |
| PA14\_31870 | NA | RND efflux membrane fusion protein | membrane, transmembrane transporter activity, transmembrane transport | Two-component system | -25.79007 | repression |  |
| PA14\_31970 | czcC | CzcC family cobalt/zinc/cadmium efflux transporter outer membrane protein | efflux transmembrane transporter activity, transmembrane transport |  | -26.05385 | repression |  |
| PA14\_31990 | czcB | cobalt/zinc/cadmium efflux RND transporter, membrane fusion protein, CzcB famil | membrane, transmembrane transporter activity, transmembrane transport |  | -26.04165 | repression |  |
| PA14\_32100 | xylY | toluate 1,2-dioxygenase subunit beta | cellular aromatic compound metabolic process, oxidation-reduction process | Benzoate degradation, Degradation of aromatic compounds, Fluorobenzoate degradation, Metabolic pathways, Microbial metabolism in diverse environments, Xylene degradation | -26.08987 | repression |  |
| PA14\_32110 | xylZ | toluate 1,2-dioxygenase electron transfer component | oxidoreductase activity, oxidation-reduction process, electron transfer activity, iron-sulfur cluster binding, 2 iron, 2 sulfur cluster binding | Benzoate degradation, Degradation of aromatic compounds, Fluorobenzoate degradation, Metabolic pathways, Microbial metabolism in diverse environments, Xylene degradation | -25.79972 | repression |  |
| PA14\_32230 | catC | muconolactone delta-isomerase | cellular aromatic compound metabolic process | Benzoate degradation, Degradation of aromatic compounds, Metabolic pathways, Microbial metabolism in diverse environments | -25.78463 | no impact |  |
| PA14\_32270 | NA | porin | integral component of membrane |  | -26.79213 | repression |  |
| PA14\_32300 | NA | kinase | protein kinase activity, ATP binding, protein phosphorylation, protein binding |  | -26.05170 | repression |  |
| PA14\_32470 | NA | hypothetical protein |  |  | -26.18840 | no impact |  |
| PA14\_32490 | NA | hypothetical protein |  |  | -25.90426 | repression |  |
| PA14\_32630 | NA | cytochrome P450 | iron ion binding, oxidoreductase activity, acting on paired donors, with incorporation or reduction of molecular oxygen, heme binding, oxidation-reduction process |  | -25.80891 | repression |  |
| PA14\_32700 | NA | transcriptional regulator | DNA-binding transcription factor activity, regulation of transcription, DNA-templated |  | -25.95101 | repression |  |
| PA14\_32770 | NA | hypothetical protein |  |  | -27.02224 | repression |  |
| PA14\_32780 | NA | hypothetical protein | protein transport |  | -25.91193 | repression |  |
| PA14\_32790 | NA | hypothetical protein |  |  | -25.88133 | repression |  |
| PA14\_32880 | NA | hypothetical protein |  |  | -26.32028 | repression |  |
| PA14\_32905 | NA | hypothetical protein | iron ion binding, cytoplasm, iron ion transport, enterochelin esterase activity |  | -27.10917 | NA |  |
| PA14\_32970 | NA | transcriptional regulator | DNA-binding transcription factor activity, regulation of transcription, DNA-templated |  | -25.97577 | repression |  |
| PA14\_33040 | gcvT2 | glycine cleavage system protein T2 | protein binding, aminomethyltransferase activity, glycine catabolic process | Biosynthesis of antibiotics, Biosynthesis of secondary metabolites, Carbon metabolism, Glycine, serine and threonine metabolism, Glycine, serine and threonine metabolism, Glycine, serine and threonine metabolism;Nitrogen metabolism;One carbon pool by folate, Glyoxylate and dicarboxylate metabolism, Metabolic pathways, One carbon pool by folate, One carbon pool by folate | -25.82749 | repression |  |
| PA14\_33120 | NA | hypothetical protein |  |  | -26.32505 | repression |  |
| PA14\_33170 | NA | transcriptional regulator | DNA-binding transcription factor activity, regulation of transcription, DNA-templated |  | -25.90045 | repression |  |
| PA14\_33190 | NA | hypothetical protein | plasma membrane, transmembrane transporter activity, transmembrane transport |  | -26.25547 | repression |  |
| PA14\_33280 | pvdL | peptide synthase | phosphopantetheine binding, lipid biosynthetic process, catalytic activity | pyoverdine synthesis | -25.86004 | repression |  |
| PA14\_33410 | NA | porin | integral component of membrane |  | -26.63941 | repression |  |
| PA14\_33420 | NA | hydrolase | catalytic activity |  | -26.15758 | repression |  |
| PA14\_33520 | NA | thioesterase | biosynthetic process, hydrolase activity, acting on ester bonds |  | -25.94658 | repression |  |
| PA14\_33690 | pvdE | pyoverdine biosynthesis protein PvdE | ATP binding, integral component of membrane, ATPase-coupled transmembrane transporter activity, transmembrane transport, peptide transport, peptide transmembrane transporter activity, ATPase activity | ABC transporters | -25.81494 | repression |  |
| PA14\_33740 | pvdP | protein PvdP |  |  | -26.29546 | repression |  |
| PA14\_33760 | NA | ABC transporter ATP-binding protein/permease | ATP binding, ATPase activity, efflux transmembrane transporter activity, xenobiotic detoxification by transmembrane export across the plasma membrane, membrane | ABC transporters | -25.93252 | repression |  |
| PA14\_33800 | NA | RNA polymerase sigma factor | DNA-binding transcription factor activity, DNA-templated transcription, initiation, regulation of transcription, DNA-templated, DNA binding, sigma factor activity |  | -25.93390 | repression |  |
| PA14\_33810 | pvdA | L-ornithine N5-oxygenase |  |  | -26.19276 | repression |  |
| PA14\_33820 | pvdQ | penicillin acylase-related protein | hydrolase activity, acting on carbon-nitrogen (but not peptide) bonds, in linear amides, hydrolase activity, antibiotic biosynthetic process, N-acetyl-anhydromuramoyl-L-alanine amidase activity |  | -25.78850 | repression |  |
| PA14\_33830 | NA | hypothetical protein |  |  | -26.88351 | repression |  |
| PA14\_33880 | NA | hypothetical protein |  |  | -26.72797 | repression |  |
| PA14\_33920 | NA | transcriptional regulator | phosphorelay signal transduction system, regulation of transcription, DNA-templated, DNA binding |  | -27.47880 | repression |  |
| PA14\_33940 | tseF | TseF |  |  | -26.08340 | repression |  |
| PA14\_34020 | hsiF3 | HsiF3 |  |  | -26.18784 | repression |  |
| PA14\_34070 | hsiB3 | HsiB3 |  |  | -27.23878 | repression |  |
| PA14\_34100 | hsiJ3 | HsiJ3 |  |  | -26.52365 | no impact |  |
| PA14\_34110 | dotU3 | DotU3 |  |  | -26.65398 | repression |  |
| PA14\_34250 | NA | glycerophosphoryl diester phosphodiesterase | lipid metabolic process, phosphoric diester hydrolase activity | Glycerophospholipid metabolism | -26.65768 | repression |  |
| PA14\_34540 | NA | xenobiotic compound DszA family monooxygenase | monooxygenase activity, oxidoreductase activity, acting on paired donors, with incorporation or reduction of molecular oxygen, oxidation-reduction process |  | -26.58070 | repression |  |
| PA14\_34720 | NA | hypothetical protein |  |  | -26.23007 | repression |  |
| PA14\_34740 | NA | hypothetical protein |  |  | -26.13019 | repression |  |
| PA14\_34810 | NA | non-ribosomal peptide synthetase | catalytic activity |  | -25.85860 | repression |  |
| PA14\_34820 | NA | regulatory protein | oxidoreductase activity, oxidation-reduction process |  | -25.94038 | repression |  |
| PA14\_34900 | NA | oxidoreductase | oxidoreductase activity, oxidation-reduction process |  | -26.09934 | repression |  |
| PA14\_34940 | NA | hypothetical protein |  |  | -26.24789 | no impact |  |
| PA14\_35060 | NA | hypothetical protein |  |  | -26.46564 | no impact |  |
| PA14\_35170 | NA | redox-sensing activator of soxS | DNA binding, regulation of transcription, DNA-templated, response to oxidative stress, 2 iron, 2 sulfur cluster binding |  | -25.89750 | no impact |  |
| PA14\_35200 | NA | acetyltransferase | N-acetyltransferase activity |  | -26.35281 | repression |  |
| PA14\_35290 | gnd | gluconate dehydrogenase | oxidoreductase activity, acting on CH-OH group of donors, oxidation-reduction process, flavin adenine dinucleotide binding | Metabolic pathways, Microbial metabolism in diverse environments, Pentose phosphate pathway | -26.39052 | no impact |  |
| PA14\_35500 | bkdB | branched-chain alpha-keto acid dehydrogenase subunit E2 | transferase activity, transferring acyl groups | Biosynthesis of antibiotics, Biosynthesis of secondary metabolites, Metabolic pathways, Propanoate metabolism, Valine, leucine and isoleucine degradation | -26.23396 | repression |  |
| PA14\_35590 | pslM | FAD-binding dehydrogenase |  |  | -26.00246 | repression |  |
| PA14\_35640 | pslI | transferase |  |  | -26.08339 | NA |  |
| PA14\_35670 | pslG | glycosyl hydrolase | hydrolase activity, hydrolyzing O-glycosyl compounds, carbohydrate metabolic process | Biofilm formation - Pseudomonas aeruginosa | -26.27623 | repression |  |
| PA14\_36060 | NA | hypothetical protein | catalytic activity |  | -26.28597 | repression |  |
| PA14\_36120 | NA | MFS transporter | integral component of plasma membrane, transmembrane transport |  | -25.82691 | repression |  |
| PA14\_36230 | NA | amino acid ABC transporter permease | membrane, transmembrane transport, integral component of membrane, transmembrane transporter activity, ATP-binding cassette (ABC) transporter complex, nitrogen compound transport |  | -26.12596 | repression |  |
| PA14\_36250 | NA | hypothetical protein |  |  | -25.95537 | no impact |  |
| PA14\_36270 | NA | dehydrogenase | oxidoreductase activity, oxidation-reduction process, NADP binding, NAD binding |  | -26.51357 | repression |  |
| PA14\_36350 | NA | hypothetical protein |  |  | -26.13276 | repression |  |
| PA14\_36360 | NA | hypothetical protein | transmembrane transport |  | -26.50085 | repression |  |
| PA14\_36420 | NA | sensor/response regulator hybrid | phosphorylation, transferase activity, transferring phosphorus-containing groups, phosphorelay sensor kinase activity, signal transduction, phosphorelay signal transduction system |  | -26.82739 | repression |  |
| PA14\_36500 | NA | hypothetical protein |  |  | -25.94253 | repression |  |
| PA14\_36530 | NA | hypothetical protein |  |  | -25.77286 | repression |  |
| PA14\_36550 | NA | hypothetical protein |  |  | -25.91810 | no impact |  |
| PA14\_36570 | glgA | glycogen synthase | glycogen (starch) synthase activity | Biosynthesis of secondary metabolites, Metabolic pathways, Starch and sucrose metabolism | -26.89338 | repression |  |
| PA14\_36670 | NA | hypothetical protein |  |  | -26.57906 | no impact |  |
| PA14\_36810 | katE | hydroperoxidase II | catalase activity, response to oxidative stress, heme binding, oxidation-reduction process | Biosynthesis of antibiotics, Biosynthesis of secondary metabolites, Carbon metabolism, Glyoxylate and dicarboxylate metabolism, Glyoxylate and dicarboxylate metabolism, Tryptophan metabolism, Tryptophan metabolism | -26.88837 | repression |  |
| PA14\_36820 | NA | hypothetical protein |  |  | -25.98320 | no impact |  |
| PA14\_36850 | NA | hypothetical protein |  |  | -26.54302 | repression |  |
| PA14\_37080 | NA | hypothetical protein | transcription regulatory region DNA binding |  | -26.54359 | repression |  |
| PA14\_37090 | NA | aldehyde dehydrogenase | oxidoreductase activity, oxidation-reduction process, oxidoreductase activity, acting on the aldehyde or oxo group of donors, NAD or NADP as acceptor |  | -25.89539 | repression |  |
| PA14\_37120 | NA | LysR family transcriptional regulator | DNA-binding transcription factor activity, regulation of transcription, DNA-templated |  | -26.33448 | repression |  |
| PA14\_37140 | NA | LysR family transcriptional regulator | DNA-binding transcription factor activity, regulation of transcription, DNA-templated |  | -26.19753 | repression |  |
| PA14\_37200 | NA | hypothetical protein |  |  | -26.44452 | repression |  |
| PA14\_37350 | NA | hypothetical protein |  |  | -26.66673 | repression |  |
| PA14\_37410 | NA | hypothetical protein |  |  | -26.11195 | repression |  |
| PA14\_37420 | NA | transmembrane sensor protein |  |  | -26.46435 | no impact |  |
| PA14\_37440 | NA | MFS transporter | integral component of plasma membrane, transmembrane transport |  | -27.04293 | repression |  |
| PA14\_37510 | NA | hypothetical protein |  |  | -26.73276 | repression |  |
| PA14\_37560 | NA | asparagine synthetase, glutamine-hydrolysing | asparagine synthase (glutamine-hydrolyzing) activity, asparagine biosynthetic process | Alanine, aspartate and glutamate metabolism, Alanine, aspartate and glutamate metabolism, Biosynthesis of secondary metabolites, Metabolic pathways | -26.58029 | repression |  |
| PA14\_37580 | NA | leucine-responsive regulatory protein | DNA-binding transcription factor activity, regulation of transcription, DNA-templated, sequence-specific DNA binding |  | -25.82896 | no impact |  |
| PA14\_37730 | NA | TonB dependent receptor |  |  | -26.50438 | repression |  |
| PA14\_37810 | pcoB | copper resistance protein B | copper ion binding, cellular copper ion homeostasis, cell outer membrane |  | -26.34793 | repression |  |
| PA14\_37830 | NA | pyridoxal-phosphate dependent protein | catalytic activity | Metabolic pathways, Sulfur relay system, Thiamine metabolism, Thiamine metabolism, [2Fe-2S] iron-sulfur cluster biosynthesis | -26.09970 | repression |  |
| PA14\_37840 | sppD | ABC transporter ATP-binding protein, SppD | ATP binding, ATPase activity, nucleotide binding, peptide transport | ABC transporters | -25.76776 | no impact |  |
| PA14\_38110 | NA | serine/threonine transporter SstT | symporter activity, integral component of membrane, threonine transport, serine transport |  | -26.54428 | repression |  |
| PA14\_38190 | NA | hypothetical protein | methyltransferase activity |  | -26.61305 | repression |  |
| PA14\_38200 | NA | thiamine pyrophosphate protein | catalytic activity, thiamine pyrophosphate binding, magnesium ion binding | 2-Oxocarboxylic acid metabolism, Biosynthesis of amino acids, Biosynthesis of antibiotics, Biosynthesis of secondary metabolites, Butanoate metabolism, C5-Branched dibasic acid metabolism, Metabolic pathways, Pantothenate and CoA biosynthesis, Valine, leucine and isoleucine biosynthesis | -26.91223 | repression |  |
| PA14\_38210 | NA | hypothetical protein |  |  | -26.41769 | repression |  |
| PA14\_38270 | NA | hypothetical protein |  |  | -25.87066 | no impact |  |
| PA14\_38360 | NA | nucleotide sugar dehydrogenase | oxidoreductase activity, acting on the CH-OH group of donors, NAD or NADP as acceptor, NAD binding, oxidation-reduction process, polysaccharide biosynthetic process, UDP-glucose 6-dehydrogenase activity | Amino sugar and nucleotide sugar metabolism, Ascorbate and aldarate metabolism, Metabolic pathways, Pentose and glucuronate interconversions | -25.77618 | no impact |  |
| PA14\_38395 | NA | periplasmic multidrug efflux lipoprotein | membrane, transmembrane transporter activity, transmembrane transport | beta-Lactam resistance, Two-component system | -26.12439 | repression |  |
| PA14\_38490 | gnyL | hydroxymethylglutaryl-CoA lyase | hydroxymethylglutaryl-CoA lyase activity, catalytic activity | Butanoate metabolism, Geraniol degradation, Metabolic pathways, Synthesis and degradation of ketone bodies, Valine, leucine and isoleucine degradation | -26.06754 | no impact |  |
| PA14\_38560 | NA | MFS transporter | integral component of plasma membrane, transmembrane transport, integral component of membrane, transmembrane transporter activity |  | -26.45491 | repression |  |
| PA14\_38610 | NA | hypothetical protein |  |  | -25.79016 | repression |  |
| PA14\_38690 | NA | acetoacetyl-CoA synthetase | catalytic activity, lipid metabolic process, acetoacetate-CoA ligase activity | Butanoate metabolism, Valine, leucine and isoleucine degradation | -26.53530 | repression |  |
| PA14\_38800 | pqqC | pyrroloquinoline quinone biosynthesis protein PqqC | pyrroloquinoline quinone biosynthetic process, oxidation-reduction process |  | -26.95991 | no impact |  |
| PA14\_38825 | pqqA | coenzyme PQQ synthesis protein PqqA | pyrroloquinoline quinone biosynthetic process | Pyrroloquinoline quinone biosynthesis | -26.49150 | no impact |  |
| PA14\_38900 | NA | two-component response regulator | phosphorelay signal transduction system, regulation of transcription, DNA-templated, DNA binding |  | -26.12861 | no impact |  |
| PA14\_38990 | NA | hypothetical protein |  |  | -27.85489 | no impact |  |
| PA14\_39190 | bacA | UDP pyrophosphate phosphatase | membrane, dephosphorylation, undecaprenyl-diphosphatase activity | Peptidoglycan biosynthesis | -27.46641 | repression |  |
| PA14\_39260 | NA | hypothetical protein |  |  | -26.13349 | repression |  |
| PA14\_39280 | rbsK | ribokinase | ribokinase activity, D-ribose metabolic process, phosphotransferase activity, alcohol group as acceptor, kinase activity | Pentose phosphate pathway | -26.70863 | no impact |  |
| PA14\_39300 | rbsR | ribose operon repressor RbsR | DNA binding, regulation of transcription, DNA-templated |  | -26.26325 | repression |  |
| PA14\_39500 | NA | hypothetical protein |  |  | -26.09176 | repression |  |
| PA14\_39520 | NA | hydroxylase large subunit | oxidoreductase activity, oxidation-reduction process | Metabolic pathways, Microbial metabolism in diverse environments, Purine metabolism | -26.07114 | no impact |  |
| PA14\_39560 | NA | chemotaxis transducer | signal transduction, membrane, transmembrane signaling receptor activity, chemotaxis | Bacterial chemotaxis, Two-component system | -26.48256 | repression |  |
| PA14\_39590 | metE | 5- methyltetrahydropteroyltriglutamate/homocysteine S-methyltransferase | 5-methyltetrahydropteroyltriglutamate-homocysteine S-methyltransferase activity, zinc ion binding, methionine biosynthetic process, cellular amino acid biosynthetic process | Biosynthesis of amino acids, Biosynthesis of secondary metabolites, Cysteine and methionine metabolism, Metabolic pathways, Selenocompound metabolism | -25.75036 | no impact |  |
| PA14\_39750 | NA | amino acid permease | amino acid transmembrane transport, integral component of plasma membrane, aromatic amino acid transmembrane transporter activity, aromatic amino acid transport |  | -25.79305 | repression |  |
| PA14\_39890 | phzF2 | phenazine biosynthesis protein | catalytic activity, biosynthetic process | Phenazine biosynthesis, Quorum sensing | -26.44280 | NA |  |
| PA14\_39945 | phzC2 | phenazine biosynthesis protein PhzC | catalytic activity, 3-deoxy-7-phosphoheptulonate synthase activity, aromatic amino acid family biosynthetic process | Biosynthesis of amino acids, Biosynthesis of antibiotics, Biosynthesis of secondary metabolites, Metabolic pathways, Phenazine biosynthesis, Phenylalanine, tyrosine and tryptophan biosynthesis, Quorum sensing | -28.02415 | repression |  |
| PA14\_39970 | phzA2 | phenazine biosynthesis protein | antibiotic biosynthetic process, phenazine biosynthetic process | Phenazine biosynthesis, Quorum sensing | -25.97961 | repression |  |
| PA14\_39980 | qscR | transcriptional regulator | regulation of transcription, DNA-templated, DNA binding | Phenazine biosynthesis | -27.13635 | no impact |  |
| PA14\_40020 | NA | hypothetical protein | lipid metabolic process |  | -25.90425 | no impact |  |
| PA14\_40100 | NA | hypothetical protein |  |  | -25.78868 | repression |  |
| PA14\_40120 | polB | DNA polymerase II | nucleotide binding, nucleic acid binding, DNA-directed DNA polymerase activity, DNA binding |  | -26.07252 | repression |  |
| PA14\_40230 | NA | secretion protein |  |  | -26.89720 | repression |  |
| PA14\_40250 | NA | outer membrane protein | efflux transmembrane transporter activity, transmembrane transport |  | -27.39896 | no impact |  |
| PA14\_40300 | NA | hypothetical protein |  |  | -26.64647 | repression |  |
| PA14\_40310 | NA | acyl carrier protein | phosphopantetheine binding, fatty acid biosynthetic process |  | -26.39792 | repression |  |
| PA14\_40490 | NA | hypothetical protein |  |  | -26.34378 | repression |  |
| PA14\_40510 | ccoN-2 | cbb3-type cytochrome c oxidase subunit I | oxidation-reduction process, cytochrome-c oxidase activity, aerobic respiration, integral component of membrane, heme binding, plasma membrane respiratory chain complex IV | aerobic respiration I (cytochrome c), aerobic respiration II (cytochrome c) (yeast), arsenite oxidation I (respiratory), Fe(II) oxidation, Metabolic pathways, Oxidative phosphorylation, Oxidative phosphorylation, Respiratory chain; terminal step, Two-component system | -25.80670 | repression |  |
| PA14\_40660 | NA | hypothetical protein |  |  | -25.88398 | repression |  |
| PA14\_40780 | NA | hypothetical protein |  |  | -26.00768 | repression |  |
| PA14\_40790 | NA | transcriptional regulator | DNA binding |  | -25.86526 | repression |  |
| PA14\_40830 | NA | oxidoreductase | oxidation-reduction process, oxidoreductase activity |  | -25.84296 | no impact |  |
| PA14\_40840 | NA | periplasmic protease | proteolysis, peptidase activity, serine-type endopeptidase activity, plasma membrane |  | -26.35110 | repression |  |
| PA14\_40910 | NA | LysR family transcriptional regulatory protein | DNA-binding transcription factor activity, regulation of transcription, DNA-templated |  | -26.16122 | repression |  |
| PA14\_40930 | NA | hypothetical protein |  |  | -25.93181 | repression |  |
| PA14\_40960 | NA | pilin biosynthetic protein | phosphorelay signal transduction system |  | -25.87612 | no impact |  |
| PA14\_41010 | NA | amino acid permease | membrane, transmembrane transporter activity, transmembrane transport |  | -26.02811 | repression |  |
| PA14\_41060 | rnhA | ribonuclease H | nucleic acid binding, RNA-DNA hybrid ribonuclease activity | DNA replication | -26.31384 | no impact |  |
| PA14\_41170 | fabI | NADH-dependent enoyl-ACP reductase | enoyl-[acyl-carrier-protein] reductase (NADH) activity, fatty acid biosynthetic process, oxidation-reduction process, enoyl-[acyl-carrier-protein] reductase activity | (5Z)-dodecenoate biosynthesis II, 8-amino-7-oxononanoate biosynthesis I, <i>cis</i>-vaccenate biosynthesis, Biotin metabolism, Fatty acid biosynthesis, Fatty acid biosynthesis, Fatty acid metabolism, gondoate biosynthesis (anaerobic), Metabolic pathways, mycolate biosynthesis, oleate biosynthesis IV (anaerobic), palmitate biosynthesis II (bacteria and plants), palmitoleate biosynthesis I (from (5Z)-dodec-5-enoate), stearate biosynthesis II (bacteria and plants), superpathway of mycolate biosynthesis | -26.94666 | repression |  |
| PA14\_41190 | ppiD | peptidyl-prolyl cis-trans isomerase D | peptidyl-prolyl cis-trans isomerase activity |  | -26.08300 | no impact |  |
| PA14\_41480 | nasS | hypothetical protein |  |  | -25.88176 | repression |  |
| PA14\_41530 | nirB | assimilatory nitrite reductase large subunit | oxidoreductase activity, heme binding, iron-sulfur cluster binding, oxidation-reduction process, nitrite reductase [NAD(P)H] activity, nitrate assimilation, flavin adenine dinucleotide binding, NADP binding | assimilatory sulfate reduction III, Microbial metabolism in diverse environments, Nitrogen metabolism, Sulfur metabolism | -25.85741 | repression |  |
| PA14\_41540 | nirD | assimilatory nitrite reductase small subunit | nitrite reductase [NAD(P)H] activity, oxidation-reduction process, oxidoreductase activity, 2 iron, 2 sulfur cluster binding | Microbial metabolism in diverse environments, Nitrogen metabolism | -25.96966 | no impact |  |
| PA14\_41560 | NA | assimilatory nitrate reductase | oxidoreductase activity, oxidation-reduction process, molybdopterin cofactor binding, 4 iron, 4 sulfur cluster binding, electron transfer activity | Microbial metabolism in diverse environments, nitrate reduction IV (dissimilatory), Nitrogen metabolism, Nitrogen metabolism | -26.12420 | repression |  |
| PA14\_41570 | oprF | major porin and structural outer membrane porin OprF precursor | cell outer membrane, integral component of membrane, calcium ion binding, porin activity, outer membrane, adhesion of symbiont to host |  | -26.10800 | repression |  |
| PA14\_41575 | sigX | RNA polymerase sigma factor SigX | DNA-binding transcription factor activity, DNA-templated transcription, initiation, regulation of transcription, DNA-templated, sigma factor activity, positive regulation of cell growth |  | -26.54150 | repression |  |
| PA14\_41730 | NA | hypothetical protein | ATP binding, metal ion binding |  | -26.37460 | no impact |  |
| PA14\_41750 | NA | hypothetical protein |  |  | -26.06864 | no impact |  |
| PA14\_41780 | NA | hypothetical protein | ATP binding, proteolysis, peptidase activity, integral component of membrane |  | -26.27066 | repression |  |
| PA14\_41790 | NA | hypothetical protein |  |  | -26.30129 | repression |  |
| PA14\_41860 | NA | hypothetical protein |  |  | -25.86570 | no impact |  |
| PA14\_41880 | NA | universal stress protein |  |  | -27.67748 | repression |  |
| PA14\_41900 | panE | 2-dehydropantoate 2-reductase | oxidation-reduction process, oxidoreductase activity, 2-dehydropantoate 2-reductase activity, pantothenate biosynthetic process | Biosynthesis of secondary metabolites, Metabolic pathways, Pantothenate and CoA biosynthesis, Pantothenate and CoA biosynthesis, phosphopantothenate biosynthesis III (archaebacteria) | -27.02706 | repression |  |
| PA14\_41910 | NA | hypothetical protein |  |  | -25.95312 | repression |  |
| PA14\_41930 | NA | hypothetical protein |  |  | -26.25638 | no impact |  |
| PA14\_42020 | NA | hypothetical protein | protein binding, electron transfer activity, protein disulfide oxidoreductase activity, cell redox homeostasis |  | -27.02148 | repression |  |
| PA14\_42030 | NA | hypothetical protein |  |  | -25.96684 | repression |  |
| PA14\_42080 | NA | 3-hydroxyacyl-CoA dehydrogenase | 3-hydroxyacyl-CoA dehydrogenase activity, fatty acid metabolic process, oxidoreductase activity, oxidation-reduction process, catalytic activity | Benzoate degradation, beta-Alanine metabolism, Biosynthesis of antibiotics, Biosynthesis of secondary metabolites, Biosynthesis of unsaturated fatty acids, Butanoate metabolism, Caprolactam degradation, Carbon metabolism, Fatty acid degradation, Fatty acid metabolism, Geraniol degradation, Limonene and pinene degradation, Lysine degradation, Metabolic pathways, Microbial metabolism in diverse environments, Propanoate metabolism, Tryptophan metabolism, Valine, leucine and isoleucine degradation | -26.63659 | repression |  |
| PA14\_42090 | NA | acetyl-CoA acetyltransferase | transferase activity, transferring acyl groups other than amino-acyl groups, catalytic activity | Benzoate degradation, Biosynthesis of antibiotics, Biosynthesis of secondary metabolites, Butanoate metabolism, Carbon metabolism, Fatty acid degradation, Fatty acid metabolism, Glyoxylate and dicarboxylate metabolism, Lysine degradation, Metabolic pathways, Microbial metabolism in diverse environments, Propanoate metabolism, Pyruvate metabolism, Synthesis and degradation of ketone bodies, Terpenoid backbone biosynthesis, Tryptophan metabolism, Two-component system, Valine, leucine and isoleucine degradation | -25.91760 | no impact |  |
| PA14\_42100 | NA | hypothetical protein | integral component of membrane, transmembrane transport |  | -27.17437 | repression |  |
| PA14\_42130 | NA | hypothetical protein | threonine-type endopeptidase activity, proteasome core complex, proteolysis involved in cellular protein catabolic process |  | -27.53686 | repression |  |
| PA14\_42150 | NA | hypothetical protein |  |  | -26.21547 | repression |  |
| PA14\_42200 | NA | hypothetical protein |  |  | -25.86085 | repression |  |
| PA14\_42280 | pscI | type III export protein PscI | protein secretion, protein secretion by the type III secretion system |  | -26.06406 | repression |  |
| PA14\_42310 | pscF | type III export protein PscF | type III protein secretion system complex, pathogenesis, protein transport, protein secretion by the type III secretion system | Bacterial secretion system | -25.92634 | no impact |  |
| PA14\_42340 | pscD | type III export protein PscD | protein secretion by the type III secretion system |  | -26.56853 | repression |  |
| PA14\_42360 | pscB | type III export apparatus protein | cytoplasm, regulation of protein secretion, protein secretion by the type III secretion system |  | -26.67391 | no impact |  |
| PA14\_42380 | NA | hypothetical protein | negative regulation of protein secretion, negative regulation of transcription, DNA-templated, negative regulation of DNA binding, negative regulation of protein binding, cellular response to calcium ion |  | -25.94342 | repression |  |
| PA14\_42390 | exsA | transcriptional regulator ExsA | DNA binding, DNA-binding transcription factor activity, regulation of transcription, DNA-templated, sequence-specific DNA binding | Biofilm formation - Pseudomonas aeruginosa | -26.71337 | repression |  |
| PA14\_42450 | popB | translocator protein PopB | pathogenesis, chaperone binding, translocation of peptides or proteins into host |  | -26.42863 | no impact |  |
| PA14\_42530 | NA | type III secretion protein | protein secretion |  | -25.84286 | repression |  |
| PA14\_42540 | NA | protein in type III secretion | negative regulation of protein secretion |  | -26.40519 | repression |  |
| PA14\_42770 | NA | hypothetical protein |  |  | -26.91876 | repression |  |
| PA14\_42870 | NA | hypothetical protein |  |  | -26.13516 | repression |  |
| PA14\_42910 | dotU2 | DotU2 |  |  | -26.91400 | no impact |  |
| PA14\_42950 | fha2 | Fha2 | protein binding |  | -26.53355 | repression |  |
| PA14\_42980 | clpV2 | ClpV2 | ATP binding, protein metabolic process | Bacterial secretion system, Biofilm formation - Pseudomonas aeruginosa | -26.04647 | repression |  |
| PA14\_42990 | hsiH2 | HsiH2 |  |  | -27.10569 | repression |  |
| PA14\_43040 | hsiB2 | HsiB2 | protein secretion by the type VI secretion system | Biofilm formation - Pseudomonas aeruginosa | -26.45089 | repression |  |
| PA14\_43220 | NA | methyl-accepting chemotaxis transducer | signal transduction, membrane, integral component of membrane | Bacterial chemotaxis, Two-component system | -26.16862 | repression |  |
| PA14\_43270 | NA | tRNA 2-selenouridine synthase | transferase activity, transferring selenium-containing groups, tRNA 2-selenouridine synthase activity, tRNA seleno-modification |  | -25.83102 | no impact |  |
| PA14\_43405 | kdbF | potassium-transporting ATPase subunit F | plasma membrane, potassium transmembrane transporter activity, phosphorylative mechanism, regulation of ATPase activity | Two-component system | -26.11260 | repression |  |
| PA14\_43420 | NA | acyl-CoA dehydrogenase | oxidoreductase activity, acting on the CH-CH group of donors, oxidation-reduction process, acyl-CoA dehydrogenase activity, flavin adenine dinucleotide binding | Caprolactam degradation, Metabolic pathways, Microbial metabolism in diverse environments | -25.97633 | repression |  |
| PA14\_43460 | NA | 3-hydroxyacyl-CoA dehydrogenase | 3-hydroxyacyl-CoA dehydrogenase activity, fatty acid metabolic process, oxidoreductase activity, oxidation-reduction process | (4Z,7Z,10Z,13Z,16Z)-docosapentaenoate biosynthesis (6-desaturase), (8<i>E</i>,10<i>E</i>)-dodeca-8,10-dienol biosynthesis, (R)- and (S)-3-hydroxybutanoate biosynthesis (engineered), 2-methylpropene degradation, 3-hydroxypropanoate/4-hydroxybutanate cycle, 4-coumarate degradation (aerobic), 4-coumarate degradation (anaerobic), 4-hydroxybenzoate biosynthesis III (plants), <i>Spodoptera littoralis</i> pheromone biosynthesis, alpha-Linolenic acid metabolism, Aminobenzoate degradation, androstenedione degradation, Benzoate degradation, Benzoate degradation, benzoyl-CoA degradation I (aerobic), beta-Alanine metabolism, Butanoate metabolism, Butanoate metabolism, Caprolactam degradation, Carbon fixation pathways in prokaryotes, cholesterol degradation to androstenedione I (cholesterol oxidase), cholesterol degradation to androstenedione II (cholesterol dehydrogenase), crotonate fermentation (to acetate and cyclohexane carboxylate), docosahexaenoate biosynthesis III (6-desaturase, mammals), fatty acid &beta;-oxidation II (peroxisome), Fatty acid degradation, Fatty acid elongation, fatty acid salvage, fermentation to 2-methylbutanoate, Geraniol degradation, glutaryl-CoA degradation, jasmonic acid biosynthesis, Limonene and pinene degradation, Lysine degradation, Metabolic pathways, methyl <i>tert</i>-butyl ether degradation, methyl ketone biosynthesis (engineered), Microbial metabolism in diverse environments, Phenylalanine metabolism, Phenylalanine metabolism, Propanoate metabolism, pyruvate fermentation to butanol I, pyruvate fermentation to butanol II (engineered), pyruvate fermentation to hexanol (engineered), Toluene degradation, Tryptophan metabolism, unsaturated, even numbered fatty acid &beta;-oxidation, Valine, leucine and isoleucine degradation | -26.12471 | repression |  |
| PA14\_43760 | NA | hypothetical protein |  |  | -27.05185 | no impact |  |
| PA14\_43820 | NA | transcriptional regulator | regulation of transcription, DNA-templated, DNA-binding transcription factor activity, sequence-specific DNA binding, DNA binding |  | -26.41731 | repression |  |
| PA14\_43870 | NA | hypothetical protein |  |  | -26.11220 | repression |  |
| PA14\_43880 | NA | hypothetical protein |  |  | -25.90407 | repression |  |
| PA14\_43910 | NA | hypothetical protein |  |  | -25.90105 | repression |  |
| PA14\_43920 | braB | branched chain amino acid transporter | branched-chain amino acid transmembrane transporter activity, integral component of membrane, branched-chain amino acid transport |  | -26.73978 | repression |  |
| PA14\_44100 | NA | hypothetical protein |  |  | -26.59117 | repression |  |
| PA14\_44120 | NA | 3-hydroxyisobutyrate dehydrogenase | NADP binding, oxidoreductase activity, oxidation-reduction process, NAD binding |  | -25.83289 | repression |  |
| PA14\_44130 | NA | hypothetical protein | nucleic acid binding |  | -26.22370 | repression |  |
| PA14\_44150 | NA | hypothetical protein |  |  | -27.39948 | repression |  |
| PA14\_44160 | NA | ATP-NAD kinase | NAD+ kinase activity, NADP biosynthetic process | NAD/NADH phosphorylation and dephosphorylation, NAD/NADP-NADH/NADPH cytosolic interconversion (yeast), NAD/NADP-NADH/NADPH mitochondrial interconversion (yeast), Nicotinate and nicotinamide metabolism | -26.70352 | no impact |  |
| PA14\_44340 | NA | cbb3-type cytochrome c oxidase subunit I | cytochrome-c oxidase activity, plasma membrane respiratory chain complex IV, oxidation-reduction process, aerobic respiration, integral component of membrane, heme binding | Metabolic pathways, Oxidative phosphorylation, Two-component system | -25.78613 | repression |  |
| PA14\_44440 | NA | cation-transporting P-type ATPase | integral component of membrane, metal ion transport, metal ion binding, cation transport, ATPase-coupled cation transmembrane transporter activity, nucleotide binding |  | -26.17072 | repression |  |
| PA14\_44510 | NA | hypothetical protein |  |  | -26.10331 | repression |  |
| PA14\_44640 | NA | hypothetical protein |  |  | -25.95118 | repression |  |
| PA14\_44670 | zipA | cell division protein ZipA | cell septum assembly, integral component of membrane |  | -26.98139 | repression |  |
| PA14\_44690 | NA | GntR family transcriptional regulator | DNA-binding transcription factor activity, regulation of transcription, DNA-templated |  | -26.46880 | no impact |  |
| PA14\_44700 | alkB2 | alkane-1 monooxygenase | oxidoreductase activity, lipid metabolic process | Aliphatic compound catabolism, Caprolactam degradation, Fatty acid degradation | -26.94204 | repression |  |
| PA14\_44890 | hcpA | secreted protein Hcp |  |  | -27.62500 | NA |  |
| PA14\_44900 | NA | hypothetical protein |  |  | -27.15546 | no impact |  |
| PA14\_44910 | NA | hypothetical protein |  |  | -25.88364 | repression |  |
| PA14\_44920 | NA | hypothetical protein |  |  | -26.47918 | repression |  |
| PA14\_44930 | NA | hypothetical protein |  |  | -26.46739 | repression |  |
| PA14\_45130 | NA | transporter | integral component of membrane, ethanolamine transmembrane transporter activity, ethanolamine transport |  | -26.40485 | repression |  |
| PA14\_45250 | NA | transcriptional regulator | DNA binding, regulation of transcription, DNA-templated |  | -26.37498 | repression |  |
| PA14\_45470 | NA | hypothetical protein | protein binding | Glutathione metabolism | -25.95135 | repression |  |
| PA14\_45520 | NA | plasmid partitioning protein |  |  | -26.03872 | repression |  |
| PA14\_45560 | motC | flagellar motor protein | bacterial-type flagellum-dependent cell motility, bacterial-type flagellum-dependent swarming motility | Bacterial chemotaxis, Flagellar assembly, Two-component system | -26.58162 | repression |  |
| PA14\_45580 | NA | chemotaxis-specific methylesterase | phosphorelay response regulator activity, phosphorelay signal transduction system, cytoplasm, chemotaxis, protein-glutamate methylesterase activity | Bacterial chemotaxis, Two-component system | -25.96472 | repression |  |
| PA14\_45610 | cheZ | chemotaxis protein CheZ | catalytic activity, bacterial-type flagellum, regulation of chemotaxis, chemotaxis, bacterial-type flagellum-dependent swarming motility | Bacterial chemotaxis | -26.24730 | no impact |  |
| PA14\_45640 | fleN | flagellar synthesis regulator FleN | ATP binding |  | -27.02558 | repression |  |
| PA14\_45700 | NA | hypothetical protein |  |  | -25.89156 | repression |  |
| PA14\_45720 | flhB | flagellar biosynthesis protein FlhB | protein secretion, membrane, protein transport, integral component of membrane, bacterial-type flagellum assembly | Flagellar assembly | -26.68334 | no impact |  |
| PA14\_45790 | fliN | flagellar motor switch protein | chemotaxis, bacterial-type flagellum, membrane, bacterial-type flagellum-dependent cell motility, motor activity, bacterial-type flagellum basal body | Bacterial chemotaxis, Flagellar assembly | -26.25479 | repression |  |
| PA14\_45810 | fliL | flagellar basal body-associated protein FliL | chemotaxis, bacterial-type flagellum basal body, bacterial-type flagellum-dependent cell motility |  | -26.15769 | repression |  |
| PA14\_45880 | NA | two-component response regulator | phosphorelay signal transduction system, DNA binding, regulation of transcription, DNA-templated | Two-component system | -26.46480 | no impact |  |
| PA14\_45910 | NA | RND efflux membrane fusion protein | membrane, transmembrane transporter activity, transmembrane transport |  | -26.60681 | repression |  |
| PA14\_45950 | rsaL | regulatory protein RsaL | quorum sensing, regulation of transcription, DNA-templated, negative regulation of secondary metabolite biosynthetic process, negative regulation of elastin catabolic process, negative regulation of cytolysis in other organism, negative regulation of cell motility, positive regulation of single-species biofilm formation, DNA binding |  | -25.82502 | no impact |  |
| PA14\_45960 | lasR | transcriptional regulator LasR | regulation of transcription, DNA-templated, DNA binding | Biofilm formation - Pseudomonas aeruginosa, Quorum sensing | -26.54263 | repression |  |
| PA14\_46010 | NA | ABC transporter ATP-binding protein | ATP binding, ATPase activity |  | -26.20967 | repression |  |
| PA14\_46160 | NA | hypothetical protein |  |  | -25.86268 | repression |  |
| PA14\_46250 | NA | hypothetical protein | methyltransferase activity |  | -25.81420 | repression |  |
| PA14\_46290 | NA | TetR family transcriptional regulator | DNA binding |  | -26.06230 | no impact |  |
| PA14\_46340 | NA | hypothetical protein |  |  | -26.62954 | repression |  |
| PA14\_46370 | NA | two-component sensor | phosphorelay signal transduction system, phosphorelay sensor kinase activity, signal transduction, phosphorylation, transferase activity, transferring phosphorus-containing groups |  | -27.09095 | repression |  |
| PA14\_46420 | NA | short chain dehydrogenase | oxidoreductase activity |  | -25.83749 | repression |  |
| PA14\_46480 | NA | transcriptional regulator |  |  | -25.80681 | repression |  |
| PA14\_46490 | fabF2 | 3-oxoacyl-ACP synthase | catalytic activity, fatty acid biosynthetic process, transferase activity, transferring acyl groups other than amino-acyl groups | Biotin metabolism, Fatty acid biosynthesis, Fatty acid metabolism, Metabolic pathways | -25.82721 | repression |  |
| PA14\_46650 | NA | transmembrane sensor |  |  | -26.36611 | no impact |  |
| PA14\_46670 | NA | hypothetical protein |  |  | -25.95791 | repression |  |
| PA14\_46710 | NA | transcriptional regulator | DNA binding |  | -27.22564 | no impact |  |
| PA14\_46760 | NA | hypothetical protein |  |  | -26.12279 | repression |  |
| PA14\_46770 | NA | hypothetical protein |  |  | -26.54755 | repression |  |
| PA14\_46930 | NA | ABC transporter permease | membrane, transmembrane transport, integral component of membrane, transmembrane transporter activity, ATP-binding cassette (ABC) transporter complex, nitrogen compound transport | ABC transporters, Two-component system | -26.11625 | repression |  |
| PA14\_47010 | NA | hypothetical protein |  |  | -25.78725 | repression |  |
| PA14\_47040 | NA | TerC family protein | integral component of membrane, flavin adenine dinucleotide binding |  | -26.10806 | repression |  |
| PA14\_47090 | NA | protease | nucleic acid binding, hydrolase activity, metal ion binding, proteolysis, serine-type peptidase activity |  | -26.71298 | repression |  |
| PA14\_47140 | NA | TonB-dependent receptor | cell outer membrane, siderophore uptake transmembrane transporter activity, siderophore transport, signaling receptor activity |  | -25.81110 | repression |  |
| PA14\_47190 | cyoB | cytochrome o ubiquinol oxidase subunit I | oxidation-reduction process, cytochrome-c oxidase activity, aerobic respiration, integral component of membrane, heme binding, oxidoreductase activity, acting on diphenols and related substances as donors, oxygen as acceptor | Metabolic pathways, Oxidative phosphorylation | -26.40979 | repression |  |
| PA14\_47210 | cyoA | cytochrome o ubiquinol oxidase subunit II | cytochrome o ubiquinol oxidase activity, integral component of membrane, electron transport chain, oxidation-reduction process, cytochrome-c oxidase activity, copper ion binding, membrane, oxidoreductase activity, acting on diphenols and related substances as donors, oxygen as acceptor | Metabolic pathways, Oxidative phosphorylation | -26.01759 | repression |  |
| PA14\_47320 | NA | hypothetical protein |  |  | -26.33781 | repression |  |
| PA14\_47330 | NA | hypothetical protein |  |  | -26.17864 | repression |  |
| PA14\_47370 | NA | signal peptidase | proteolysis, serine-type peptidase activity, membrane, integral component of membrane | Protein export, Quorum sensing | -25.92900 | repression |  |
| PA14\_47390 | NA | transmembrane sensor |  |  | -26.44576 | repression |  |
| PA14\_47400 | NA | RNA polymerase ECF-subfamily sigma-70 factor | DNA-binding transcription factor activity, DNA-templated transcription, initiation, regulation of transcription, DNA-templated, DNA binding, sigma factor activity |  | -27.30230 | repression |  |
| PA14\_47500 | sseA | 3-mercaptopyruvate sulfurtransferase |  |  | -26.49995 | no impact |  |
| PA14\_47610 | NA | transcriptional regulator | DNA binding |  | -26.98664 | repression |  |
| PA14\_47650 | cobS | cobalamin synthase | cobalamin 5'-phosphate synthase activity, cobalamin biosynthetic process, adenosylcobinamide-GDP ribazoletransferase activity | 2-methyladeninyl adenosylcobamide biosynthesis from adenosylcobinamide-GDP, 4-methylphenyl adenosylcobamide biosynthesis from adenosylcobinamide-GDP, 5-hydroxybenzimidazolyl adenosylcobamide biosynthesis from adenosylcobinamide-GDP, 5-methoxy-6-methylbenzimidazolyl adenosylcobamide biosynthesis from adenosylcobinamide-GDP, 5-methoxybenzimidazolyl adenosylcobamide biosynthesis from adenosylcobinamide-GDP, 5-methylbenzimidazolyl adenosylcobamide biosynthesis from adenosylcobinamide-GDP, adeninyl adenosylcobamide biosynthesis from adenosylcobinamide-GDP, adenosylcobalamin biosynthesis from adenosylcobinamide-GDP I, adenosylcobalamin biosynthesis from adenosylcobinamide-GDP II, benzimidazolyl adenosylcobamide biosynthesis from adenosylcobinamide-GDP, Metabolic pathways, phenyl adenosylcobamide biosynthesis from adenosylcobinamide-GDP, Porphyrin and chlorophyll metabolism, Porphyrin and chlorophyll metabolism, superpathway of adenosylcobalamin salvage from cobinamide II | -25.77193 | repression |  |
| PA14\_47690 | cobQ | cobyric acid synthase | catalytic activity, cobalamin biosynthetic process | Metabolic pathways, Porphyrin and chlorophyll metabolism | -26.33093 | repression |  |
| PA14\_47720 | cobC | threonine-phosphate decarboxylase | catalytic activity, cobalamin biosynthetic process, biosynthetic process, pyridoxal phosphate binding | Metabolic pathways, Porphyrin and chlorophyll metabolism | -26.38774 | repression |  |
| PA14\_47760 | cobB | cobyrinic acid a,c-diamide synthase | cobalamin biosynthetic process, cobyrinic acid a,c-diamide synthase activity, catalytic activity | Metabolic pathways, Porphyrin and chlorophyll metabolism | -26.00685 | repression |  |
| PA14\_47790 | cobO | cob(I)yrinic acid a,c-diamide adenosyltransferase | ATP binding, cob(I)yrinic acid a,c-diamide adenosyltransferase activity, cobalamin biosynthetic process | Metabolic pathways, Porphyrin and chlorophyll metabolism | -26.43013 | no impact |  |
| PA14\_47960 | NA | ABC transporter ATP-binding protein | ATP binding, ATPase activity, amino acid transmembrane transport, ATPase-coupled amino acid transmembrane transporter activity |  | -26.42819 | no impact |  |
| PA14\_48115 | aprD | alkaline protease secretion protein AprD | ATP binding, integral component of membrane, ATPase-coupled transmembrane transporter activity, transmembrane transport, protein secretion by the type I secretion system, type I protein secretion system complex, ATPase activity | ABC transporters | -26.67465 | repression |  |
| PA14\_48140 | NA | hypothetical protein |  |  | -26.51556 | repression |  |
| PA14\_48280 | NA | multidrug resistance efflux pump |  |  | -25.76993 | repression |  |
| PA14\_48300 | NA | MFS transporter | integral component of membrane, transmembrane transporter activity, transmembrane transport, integral component of plasma membrane |  | -25.77971 | repression |  |
| PA14\_48330 | NA | hypothetical protein |  |  | -26.62131 | repression |  |
| PA14\_48830 | NA | transcriptional regulator | ATP binding, regulation of transcription, DNA-templated, transcription factor binding, DNA binding, sequence-specific DNA binding |  | -26.09298 | repression |  |
| PA14\_48850 | NA | amino acid permease | membrane, transmembrane transporter activity, transmembrane transport, amino acid transport, integral component of membrane |  | -25.88486 | repression |  |
| PA14\_48880 | NA | bacteriophage integrase | DNA binding, DNA recombination, DNA integration |  | -26.13767 | no impact |  |
| PA14\_48890 | NA | hypothetical protein |  |  | -26.69162 | no impact |  |
| PA14\_48920 | NA | bacteriophage protein |  |  | -25.80865 | repression |  |
| PA14\_48970 | NA | helix destabilizing protein of bacteriophage Pf1 |  |  | -25.89847 | repression |  |
| PA14\_49000 | NA | hypothetical protein |  |  | -25.95352 | repression |  |
| PA14\_49050 | NA | hypothetical protein |  |  | -26.07349 | repression |  |
| PA14\_49070 | NA | hypothetical protein | CoA-transferase activity |  | -26.06063 | repression |  |
| PA14\_49130 | dctA | C4-dicarboxylate transporter DctA | symporter activity, integral component of membrane, dicarboxylic acid transport | Two-component system | -26.00502 | repression |  |
| PA14\_49210 | napE | periplasmic nitrate reductase NapE |  |  | -26.51881 | no impact |  |
| PA14\_49280 | NA | transglycosylase |  |  | -26.03826 | repression |  |
| PA14\_49420 | NA | two-component sensor | signal transduction, integral component of membrane, phosphorylation, transferase activity, transferring phosphorus-containing groups, phosphorelay sensor kinase activity | Two-component system | -26.28780 | repression |  |
| PA14\_49440 | NA | two-component response regulator | phosphorelay signal transduction system, DNA binding, regulation of transcription, DNA-templated | Two-component system | -25.85214 | repression |  |
| PA14\_49500 | NA | hypothetical protein |  |  | -26.35732 | repression |  |
| PA14\_49700 | NA | transcriptional regulator | regulation of transcription, DNA-templated, DNA binding |  | -26.01341 | NA |  |
| PA14\_49740 | NA | hypothetical protein |  |  | -26.44220 | repression |  |
| PA14\_49760 | rhlC | rhamnosyltransferase 2 |  |  | -26.82896 | repression |  |
| PA14\_49810 | NA | hypothetical protein |  |  | -27.06762 | repression |  |
| PA14\_49840 | dgt | deoxyguanosinetriphosphate triphosphohydrolase | triphosphoric monoester hydrolase activity, magnesium ion binding, dGTP catabolic process, dGTPase activity | Purine metabolism, Purine metabolism, Purine metabolism | -27.08635 | no impact |  |
| PA14\_49850 | NA | hypothetical protein |  |  | -26.22860 | repression |  |
| PA14\_49870 | NA | peptide deformylase |  |  | -25.76081 | repression |  |
| PA14\_50020 | NA | hypothetical protein |  |  | -26.72329 | repression |  |
| PA14\_50080 | fliJ | flagellar biosynthesis chaperone | bacterial-type flagellum, bacterial-type flagellum-dependent cell motility, motor activity, chemotaxis, membrane | Flagella assembly , Flagellar assembly | -25.84592 | repression |  |
| PA14\_50240 | NA | hypothetical protein |  |  | -26.49918 | repression |  |
| PA14\_50330 | NA | hypothetical protein |  |  | -26.87031 | repression |  |
| PA14\_50560 | braG | branched-chain amino acid transport protein BraG | branched-chain amino acid transmembrane transporter activity, branched-chain amino acid transport, ATP binding, ATPase activity | ABC transporters, Quorum sensing | -26.00637 | no impact |  |
| PA14\_50570 | NA | hypothetical protein | protein binding |  | -26.90941 | repression |  |
| PA14\_50590 | NA | HSP90 family protein | ATP binding, protein folding, unfolded protein binding |  | -26.43048 | repression |  |
| PA14\_50600 | NA | transcriptional regulator | DNA-binding transcription factor activity, regulation of transcription, DNA-templated |  | -26.07004 | repression |  |
| PA14\_50620 | NA | hypothetical protein |  |  | -27.12309 | repression |  |
| PA14\_50650 | NA | hypothetical protein | N-acetyltransferase activity |  | -26.27532 | repression |  |
| PA14\_50830 | NA | hypothetical protein |  |  | -25.87298 | repression |  |
| PA14\_51000 | NA | hypothetical protein |  |  | -26.53643 | no impact |  |
| PA14\_51040 | NA | oxidoreductase | oxidoreductase activity, oxidation-reduction process |  | -26.23634 | repression |  |
| PA14\_51080 | NA | dioxygenase | catalytic activity, nitronate monooxygenase activity, oxidation-reduction process | Nitrogen metabolism | -26.43890 | repression |  |
| PA14\_51150 | mucK | cis,cis-muconate transporter MucK | integral component of plasma membrane, transmembrane transport, integral component of membrane, transmembrane transporter activity |  | -26.35531 | repression |  |
| PA14\_51205 | NA | transcriptional regulator | DNA binding, regulation of transcription, DNA-templated |  | -26.20300 | repression |  |
| PA14\_51340 | mvfR | transcriptional regulator MvfR | DNA-binding transcription factor activity, regulation of transcription, DNA-templated, positive regulation of lyase activity, positive regulation of multi-organism process, regulation of transmembrane transport, transcription regulatory region DNA binding, plasma membrane | Biofilm formation - Pseudomonas aeruginosa, Quorum sensing | -26.55724 | repression |  |
| PA14\_51360 | phnA | anthranilate synthase component I | biosynthetic process, anthranilate synthase activity, phenazine biosynthetic process | 4-hydroxy-2(1<i>H</i>)-quinolone biosynthesis, acridone alkaloid biosynthesis, Biofilm formation - Pseudomonas aeruginosa, Biosynthesis of amino acids, Biosynthesis of antibiotics, Biosynthesis of secondary metabolites, Metabolic pathways, Phenazine biosynthesis, Phenylalanine, tyrosine and tryptophan biosynthesis, Phenylalanine, tyrosine and tryptophan biosynthesis, Quorum sensing | -26.38567 | repression |  |
| PA14\_51390 | pqsD | 3-oxoacyl-ACP synthase | catalytic activity, 3-oxoacyl-[acyl-carrier-protein] synthase activity, fatty acid biosynthetic process | Biofilm formation - Pseudomonas aeruginosa, Fatty acid biosynthesis (path 1) , Quorum sensing | -26.08746 | repression |  |
| PA14\_51420 | pqsB | PqsB | catalytic activity, secondary metabolite biosynthetic process | Biofilm formation - Pseudomonas aeruginosa, Quorum sensing | -25.85330 | repression |  |
| PA14\_51440 | ogt | methylated-DNA--protein-cysteine methyltransferase | catalytic activity, DNA repair, methylated-DNA-[protein]-cysteine S-methyltransferase activity |  | -26.20872 | no impact |  |
| PA14\_51450 | cupC3 | usher CupC3 | pilus assembly, fimbrial usher porin activity, membrane, protein binding |  | -26.82031 | repression |  |
| PA14\_51480 | NA | hypothetical protein | phosphorelay signal transduction system |  | -26.32230 | repression |  |
| PA14\_51500 | NA | hypothetical protein |  |  | -25.94821 | repression |  |
| PA14\_51810 | NA | hypothetical protein |  |  | -26.17124 | repression |  |
| PA14\_51830 | NA | DNA-binding stress protein | oxidoreductase activity, oxidizing metal ions, oxidation-reduction process, cellular iron ion homeostasis, ferric iron binding |  | -26.76716 | repression |  |
| PA14\_51850 | NA | hypothetical protein |  |  | -26.71951 | repression |  |
| PA14\_51890 | NA | hypothetical protein |  |  | -26.08026 | no impact |  |
| PA14\_51920 | NA | acylphosphatase |  |  | -26.18392 | repression |  |
| PA14\_51930 | NA | thioredoxin | cell redox homeostasis, antioxidant activity, oxidoreductase activity, oxidation-reduction process |  | -26.26459 | no impact |  |
| PA14\_51940 | NA | hypothetical protein |  |  | -25.90736 | repression |  |
| PA14\_51960 | NA | ribonuclease |  |  | -26.14785 | repression |  |
| PA14\_52040 | purM | phosphoribosylaminoimidazole synthetase | phosphoribosylformylglycinamidine cyclo-ligase activity, 'de novo' IMP biosynthetic process | Biosynthesis of antibiotics, Biosynthesis of secondary metabolites, Metabolic pathways, Purine metabolism, Purine metabolism | -25.75832 | no impact |  |
| PA14\_52060 | NA | hypothetical protein |  |  | -26.18165 | repression |  |
| PA14\_52090 | NA | hypothetical protein |  |  | -26.71398 | repression |  |
| PA14\_52180 | relA | GTP pyrophosphokinase | guanosine tetraphosphate metabolic process | Purine metabolism | -25.77444 | repression |  |
| PA14\_52260 | NA | sensor/response regulator hybrid | signal transduction, integral component of membrane, phosphorelay signal transduction system, phosphorelay sensor kinase activity, phosphorylation, transferase activity, transferring phosphorus-containing groups, positive regulation of cell motility, positive regulation of secondary metabolite biosynthetic process | Biofilm formation - Pseudomonas aeruginosa, Two-component system | -26.99472 | repression |  |
| PA14\_52330 | NA | hypothetical protein |  |  | -27.00334 | repression |  |
| PA14\_52340 | NA | hypothetical protein |  |  | -26.31841 | no impact |  |
| PA14\_52380 | NA | cytochrome b561 | electron transfer activity, integral component of membrane, membrane, respiratory electron transport chain |  | -26.16503 | repression |  |
| PA14\_52400 | kup | potassium uptake protein Kup | potassium ion transmembrane transporter activity, membrane, potassium ion transmembrane transport |  | -26.41985 | no impact |  |
| PA14\_52460 | mgtE | Mg transporter MgtE | cation transport, cation transmembrane transporter activity, magnesium ion transmembrane transporter activity, magnesium ion transport, membrane |  | -25.76921 | NA |  |
| PA14\_52480 | NA | hypothetical protein |  |  | -26.33184 | repression |  |
| PA14\_52520 | NA | hypothetical protein |  |  | -26.09445 | repression |  |
| PA14\_52570 | rsmA | carbon storage regulator | RNA binding, regulation of carbohydrate metabolic process, mRNA catabolic process | Biofilm formation - Pseudomonas aeruginosa, Two-component system | -26.17486 | no impact |  |
| PA14\_52580 | lysC | aspartate kinase | aspartate kinase activity, cellular amino acid biosynthetic process, lysine biosynthetic process via diaminopimelate | 2-Oxocarboxylic acid metabolism, Biosynthesis of amino acids, Biosynthesis of antibiotics, Biosynthesis of secondary metabolites, Cysteine and methionine metabolism, Glycine, serine and threonine metabolism, Glycine, serine and threonine metabolism;Lysine biosynthesis , Lysine biosynthesis, Metabolic pathways, Microbial metabolism in diverse environments, Monobactam biosynthesis | -26.56291 | no impact |  |
| PA14\_52670 | astD | succinylglutamic semialdehyde dehydrogenase | oxidoreductase activity, oxidation-reduction process, arginine catabolic process, succinylglutamate-semialdehyde dehydrogenase activity, oxidoreductase activity, acting on the aldehyde or oxo group of donors, NAD or NADP as acceptor | Arginine and proline metabolism, Arginine and proline metabolism, Arginine and proline metabolism; Butanoate metabolism; Glutamate metabolism, Metabolic pathways | -25.95964 | repression |  |
| PA14\_52700 | aruF | arginine/ornithine succinyltransferase AI subunit | arginine catabolic process, arginine N-succinyltransferase activity | Arginine and proline metabolism, Metabolic pathways | -26.19102 | repression |  |
| PA14\_52730 | NA | hypothetical protein |  |  | -26.60771 | no impact |  |
| PA14\_52900 | NA | acyl-CoA dehydrogenase | oxidoreductase activity, acting on the CH-CH group of donors, flavin adenine dinucleotide binding, oxidation-reduction process, acyl-CoA dehydrogenase activity | beta-Alanine metabolism, Biosynthesis of antibiotics, Biosynthesis of secondary metabolites, Carbon metabolism, Fatty acid degradation, Fatty acid metabolism, Metabolic pathways, Propanoate metabolism, Valine, leucine and isoleucine degradation | -26.00531 | repression |  |
| PA14\_52920 | NA | transcriptional regulator | DNA-binding transcription factor activity, regulation of transcription, DNA-templated |  | -26.25181 | no impact |  |
| PA14\_52930 | NA | transcriptional regulator | DNA-binding transcription factor activity, regulation of transcription, DNA-templated |  | -26.06311 | repression |  |
| PA14\_53140 | NA | hypothetical protein | regulation of transcription, DNA-templated |  | -26.74931 | repression |  |
| PA14\_53360 | plcH | hemolytic phospholipase C | hydrolase activity, acting on ester bonds, phosphatidylcholine phospholipase C activity, phospholipase C activity, lipid catabolic process, catalytic activity | 2-arachidonoylglycerol biosynthesis, Biosynthesis of secondary metabolites, Ether lipid metabolism, Ether lipid metabolism, Glycerophospholipid metabolism, Glycerophospholipid metabolism, Inositol phosphate metabolism, Inositol phosphate metabolism, Metabolic pathways, plasmalogen biosynthesis, plasmalogen degradation, Quorum sensing | -26.03698 | repression |  |
| PA14\_53420 | NA | glutathione peroxidase | glutathione peroxidase activity, response to oxidative stress, oxidation-reduction process | Arachidonic acid metabolism, Glutathione metabolism | -26.35472 | no impact |  |
| PA14\_53550 | NA | transcriptional regulator | DNA binding |  | -26.60532 | no impact |  |
| PA14\_53620 | NA | hypothetical protein |  |  | -27.27390 | repression |  |
| PA14\_53650 | NA | hypothetical protein | hydrolase activity, acting on ester bonds |  | -26.35420 | no impact |  |
| PA14\_53660 | NA | hypothetical protein |  |  | -26.04535 | repression |  |
| PA14\_53720 | NA | transcriptional regulator | DNA-binding transcription factor activity, regulation of transcription, DNA-templated |  | -25.82646 | repression |  |
| PA14\_53770 | NA | hypothetical protein | transferase activity, L-lysine catabolic process to acetate, catalytic activity |  | -26.84553 | repression |  |
| PA14\_53890 | NA | hypothetical protein |  |  | -26.08598 | repression |  |
| PA14\_54150 | putP | sodium/proline symporter PutP | proline:sodium symporter activity, sodium ion transport, proline transport, integral component of membrane, sodium ion binding, membrane, transmembrane transporter activity, transmembrane transport, proline catabolic process to glutamate, L-proline transmembrane transporter activity |  | -26.90802 | repression |  |
| PA14\_54240 | NA | hypothetical protein |  |  | -25.95856 | repression |  |
| PA14\_54400 | mucC | positive regulator for alginate biosynthesis MucC | regulation of polysaccharide biosynthetic process |  | -26.24973 | no impact |  |
| PA14\_54430 | algU | RNA polymerase sigma factor AlgU | DNA binding, DNA-binding transcription factor activity, DNA-templated transcription, initiation, regulation of transcription, DNA-templated, sigma factor activity, negative regulation of bacterial-type flagellum-dependent cell motility, regulation of polysaccharide biosynthetic process |  | -26.12164 | no impact |  |
| PA14\_54450 | nadB | L-aspartate oxidase | oxidoreductase activity, oxidation-reduction process, L-aspartate oxidase activity, NAD biosynthetic process | Alanine, aspartate and glutamate metabolism, Metabolic pathways, Nicotinate and nicotinamide metabolism | -25.75042 | repression |  |
| PA14\_54510 | NA | two-component response regulator | phosphorelay signal transduction system, DNA binding, regulation of transcription, DNA-templated | Two-component System, Two-component system | -26.15022 | no impact |  |
| PA14\_54590 | ung | uracil-DNA glycosylase | uracil DNA N-glycosylase activity, DNA repair, base-excision repair, hydrolase activity, hydrolyzing N-glycosyl compounds | Base excision repair | -26.09117 | no impact |  |
| PA14\_54670 | NA | 3-hydroxyisobutyrate dehydrogenase | 3-hydroxyisobutyrate dehydrogenase activity, NAD binding, oxidation-reduction process, oxidoreductase activity, NADP binding | Valine, leucine and isoleucine degradation | -25.82852 | no impact |  |
| PA14\_54760 | NA | hypothetical protein |  |  | -25.90424 | repression |  |
| PA14\_55100 | NA | hypothetical protein |  |  | -27.04477 | no impact |  |
| PA14\_55170 | cat | chloramphenicol acetyltransferase |  |  | -26.91880 | repression |  |
| PA14\_55320 | NA | hypothetical protein |  |  | -26.00816 | repression |  |
| PA14\_55390 | NA | hypothetical protein |  |  | -26.31723 | repression |  |
| PA14\_55490 | hxcT | HxcT | type II protein secretion system complex, protein secretion by the type II secretion system | Bacterial secretion system | -26.47400 | repression |  |
| PA14\_55550 | NA | ECF subfamily RNA polymerase sigma-70 factor | DNA-binding transcription factor activity, DNA-templated transcription, initiation, regulation of transcription, DNA-templated, DNA binding, sigma factor activity |  | -27.36791 | repression |  |
| PA14\_55760 | NA | hypothetical protein |  |  | -26.25671 | no impact |  |
| PA14\_55770 | NA | phosphate transporter | inorganic phosphate transmembrane transporter activity, phosphate ion transport, membrane |  | -25.90061 | repression |  |
| PA14\_55790 | NA | hypothetical protein |  |  | -25.79495 | no impact |  |
| PA14\_55800 | NA | hypothetical protein | aspartic-type endopeptidase activity, membrane | Type II secretion system | -26.01556 | repression |  |
| PA14\_55820 | NA | hypothetical protein |  |  | -26.61864 | repression |  |
| PA14\_55850 | NA | pilus assembly protein | protein binding |  | -25.98150 | no impact |  |
| PA14\_55920 | NA | type II secretion system protein | protein secretion |  | -26.32353 | repression |  |
| PA14\_55930 | NA | pilus assembly protein |  |  | -26.42236 | repression |  |
| PA14\_56000 | pctA | chemotactic transducer PctA | transmembrane signaling receptor activity, chemotaxis, signal transduction, membrane, integral component of membrane | Bacterial chemotaxis, Two-component system | -26.89068 | repression |  |
| PA14\_56070 | mvaT | transcriptional regulator MvaT, P16 subunit |  |  | -26.29723 | no impact |  |
| PA14\_56380 | NA | hypothetical protein |  |  | -26.03489 | repression |  |
| PA14\_56430 | NA | transcriptional regulator | DNA binding, regulation of transcription, DNA-templated |  | -25.90179 | repression |  |
| PA14\_56510 | NA | hypothetical protein |  |  | -26.10623 | repression |  |
| PA14\_56600 | NA | hypothetical protein | fatty acid biosynthetic process, [acyl-carrier-protein] phosphodiesterase activity |  | -26.32993 | no impact |  |
| PA14\_56620 | NA | hypothetical protein | DNA-binding transcription factor activity, regulation of transcription, DNA-templated, regulation of single-species biofilm formation on inanimate substrate, transcription regulatory region DNA binding |  | -25.97000 | repression |  |
| PA14\_56720 | NA | oxidoreductase | catalytic activity, coenzyme binding |  | -26.96950 | repression |  |
| PA14\_56890 | NA | multidrug efflux protein | membrane, transmembrane transporter activity, transmembrane transport | beta-Lactam resistance, Cationic antimicrobial peptide (CAMP) resistance | -26.04431 | repression |  |
| PA14\_56940 | NA | two-component sensor | phosphorelay sensor kinase activity, signal transduction, integral component of membrane, phosphorylation, transferase activity, transferring phosphorus-containing groups |  | -26.02488 | repression |  |
| PA14\_57030 | fxsA | FxsA protein | membrane |  | -26.22241 | repression |  |
| PA14\_57040 | NA | hypothetical protein | cofactor binding |  | -26.11177 | repression |  |
| PA14\_57050 | fabG | 3-ketoacyl-ACP reductase |  |  | -26.37681 | no impact |  |
| PA14\_57080 | NA | hypothetical protein | catalytic activity, DNA repair |  | -27.28387 | repression |  |
| PA14\_57110 | NA | hypothetical protein | membrane, transmembrane transport |  | -27.19536 | repression |  |
| PA14\_57140 | NA | two-component response regulator | phosphorelay signal transduction system |  | -26.59798 | repression |  |
| PA14\_57180 | NA | hypothetical protein |  |  | -26.03844 | repression |  |
| PA14\_57240 | NA | hypothetical protein |  |  | -26.90621 | repression |  |
| PA14\_57275 | ftsZ | cell division protein FtsZ | GTP binding, GTPase activity |  | -26.37723 | repression |  |
| PA14\_57320 | ddl | D-alanine--D-alanine ligase | cytoplasm, D-alanine-D-alanine ligase activity, ATP binding, metal ion binding | D-Alanine metabolism, D-Alanine metabolism, Metabolic pathways, Peptidoglycan biosynthesis, Peptidoglycan biosynthesis, UDP-<i>N</i>-acetylmuramoyl-pentapeptide biosynthesis I (<i>meso</i>-diaminopimelate containing), UDP-<i>N</i>-acetylmuramoyl-pentapeptide biosynthesis II (lysine-containing), UDP-<i>N</i>-acetylmuramoyl-pentapeptide biosynthesis III (<i>meso</i>-diaminopimelate containing), Vancomycin resistance | -25.91337 | repression |  |
| PA14\_57340 | murG | UDPdiphospho-muramoylpentapeptide beta-N- acetylglucosaminyltransferase | transferase activity, transferring hexosyl groups, undecaprenyldiphospho-muramoylpentapeptide beta-N-acetylglucosaminyltransferase activity, carbohydrate metabolic process, lipid glycosylation | Metabolic pathways, Peptidoglycan biosynthesis, Vancomycin resistance | -26.11866 | repression |  |
| PA14\_57490 | NA | hypothetical protein | nucleic acid binding, nuclease activity |  | -26.95879 | repression |  |
| PA14\_57520 | sspB | ClpXP protease specificity-enhancing factor |  |  | -26.06874 | repression |  |
| PA14\_57540 | NA | cytochrome c1 | electron transfer activity, heme binding | Metabolic pathways, Oxidative phosphorylation, Two-component system | -26.20564 | repression |  |
| PA14\_57710 | cysN | bifunctional sulfate adenylyltransferase subunit 1/adenylylsulfate kinase | GTPase activity, GTP binding, sulfate assimilation, adenylylsulfate kinase activity, ATP binding, sulfur compound metabolic process, cellular response to sulfate starvation | Biosynthesis of antibiotics, Metabolic pathways, Microbial metabolism in diverse environments, Monobactam biosynthesis, Purine metabolism, Selenocompound metabolism, Sulfur metabolism | -26.27893 | repression |  |
| PA14\_57740 | NA | hypothetical protein |  |  | -26.00524 | repression |  |
| PA14\_57810 | murA | UDP-N-acetylglucosamine 1-carboxyvinyltransferase | transferase activity, transferring alkyl or aryl (other than methyl) groups, UDP-N-acetylglucosamine 1-carboxyvinyltransferase activity, UDP-N-acetylgalactosamine biosynthetic process, catalytic activity | Amino sugar and nucleotide sugar metabolism, Metabolic pathways, Peptidoglycan biosynthesis | -26.16469 | repression |  |
| PA14\_57830 | NA | hypothetical protein |  |  | -26.29156 | repression |  |
| PA14\_57880 | NA | ABC transporter ATP-binding protein | ATP binding, ATPase activity | ABC transporters | -27.20033 | repression |  |
| PA14\_58090 | NA | hypothetical protein |  |  | -25.79988 | no impact |  |
| PA14\_58100 | cafA | cytoplasmic axial filament protein | RNA binding, ribonuclease activity, RNA processing, nucleic acid binding |  | -26.50096 | no impact |  |
| PA14\_58300 | NA | two-component response regulator | phosphorelay signal transduction system, sequence-specific DNA binding, transcription regulatory region DNA binding | Two-component system | -25.99365 | repression |  |
| PA14\_58375 | mdpA | metallopeptidase MdpA | hydrolase activity, peptide catabolic process |  | -25.85669 | no impact |  |
| PA14\_58390 | dppA3 | dipeptide ABC transporter substrate-binding protein DppA3 | ATP-binding cassette (ABC) transporter complex, transmembrane transport, dipeptide transport, dipeptide transport, peptide binding | ABC transporters, Bacterial chemotaxis | -26.34292 | repression |  |
| PA14\_58570 | NA | outer membrane ferric siderophore receptor | cell outer membrane, siderophore uptake transmembrane transporter activity, siderophore transport, signaling receptor activity |  | -26.24695 | repression |  |
| PA14\_58800 | NA | hypothetical protein |  |  | -26.56649 | repression |  |
| PA14\_58890 | NA | hemolysin activation/secretion protein |  |  | -25.87944 | repression |  |
| PA14\_60190 | clpB | clpB protein | protein metabolic process, ATP binding, cytoplasm, response to heat, protein refolding |  | -26.39608 | repression |  |
| PA14\_60230 | comL | competence protein ComL | protein binding |  | -26.48308 | repression |  |
| PA14\_60410 | NA | hypothetical protein |  |  | -26.14572 | repression |  |
| PA14\_60490 | NA | cytochrome c | electron transfer activity, heme binding |  | -27.04017 | repression |  |
| PA14\_60520 | NA | hypothetical protein |  |  | -25.93051 | no impact |  |
| PA14\_60540 | NA | hypothetical protein |  |  | -27.86924 | no impact |  |
| PA14\_60590 | NA | hypothetical protein |  |  | -26.24540 | no impact |  |
| PA14\_60630 | NA | hypothetical protein | membrane |  | -26.73767 | repression |  |
| PA14\_60650 | NA | hypothetical protein | RNA processing, RNA ligase activity |  | -26.23403 | repression |  |
| PA14\_60660 | NA | nucleotidyltransferase |  |  | -26.86003 | no impact |  |
| PA14\_60670 | rtcA | RNA 3'-terminal-phosphate cyclase | RNA processing, RNA-3'-phosphate cyclase activity, catalytic activity |  | -26.70312 | repression |  |
| PA14\_60790 | NA | ABC transporter ATP-binding protein | ATP binding, ATPase activity |  | -27.09936 | no impact |  |
| PA14\_60810 | NA | transcriptional regulator NfxB | DNA binding |  | -26.05989 | NA |  |
| PA14\_60850 | mexC | multidrug efflux RND membrane fusion protein | membrane, transmembrane transporter activity, transmembrane transport |  | -25.93127 | no impact |  |
| PA14\_60930 | NA | hypothetical protein |  |  | -26.03655 | no impact |  |
| PA14\_60960 | NA | hypothetical protein |  |  | -26.96288 | repression |  |
| PA14\_60990 | radA | DNA repair protein RadA | damaged DNA binding, ATP binding, DNA repair, DNA binding, DNA-dependent ATPase activity |  | -25.85632 | no impact |  |
| PA14\_61050 | mscL | large-conductance mechanosensitive channel | mechanosensitive ion channel activity, integral component of membrane, ion transmembrane transport |  | -26.49816 | repression |  |
| PA14\_61190 | NA | hypothetical protein |  |  | -25.95880 | no impact |  |
| PA14\_61200 | NA | hypothetical protein |  |  | -25.92076 | repression |  |
| PA14\_61260 | NA | hypothetical protein |  |  | -25.97979 | repression |  |
| PA14\_61270 | NA | hypothetical protein | polyketide metabolic process |  | -25.92757 | no impact |  |
| PA14\_61450 | NA | hypothetical protein |  |  | -26.52770 | repression |  |
| PA14\_61480 | uraA | uracil permease | membrane, transmembrane transporter activity, transmembrane transport |  | -26.11963 | repression |  |
| PA14\_61590 | NA | hypothetical protein | catalytic activity, coenzyme binding |  | -26.56598 | no impact |  |
| PA14\_61680 | prmC | S-adenosylmethionine-dependent methyltransferase, PrmC | methyltransferase activity, nucleic acid binding, methylation, protein methylation, protein methyltransferase activity, S-adenosylmethionine-dependent methyltransferase activity |  | -26.09569 | repression |  |
| PA14\_61880 | rimI | peptide n-acetyltransferase RimI | N-terminal protein amino acid acetylation, acetyltransferase activity, N-acetyltransferase activity |  | -25.85148 | repression |  |
| PA14\_61920 | NA | hypothetical protein |  |  | -25.78273 | repression |  |
| PA14\_61950 | NA | hypothetical protein |  |  | -26.32781 | repression |  |
| PA14\_61980 | NA | hypothetical protein |  |  | -26.25335 | repression |  |
| PA14\_62100 | NA | sulfite oxidase subunit YedZ | integral component of plasma membrane, protein repair |  | -26.17245 | repression |  |
| PA14\_62120 | pssA | phosphatidylserine synthase | phospholipid biosynthetic process, membrane, phosphotransferase activity, for other substituted phosphate groups | Biosynthesis of secondary metabolites, Glycerophospholipid metabolism, Glycine, serine and threonine metabolism, Metabolic pathways | -26.15956 | no impact |  |
| PA14\_62180 | NA | hypothetical protein |  |  | -26.45452 | repression |  |
| PA14\_62360 | NA | Rieske family iron-sulfur cluster-binding protein | oxidoreductase activity, 2 iron, 2 sulfur cluster binding, oxidation-reduction process |  | -26.20957 | repression |  |
| PA14\_62370 | NA | hypothetical protein |  |  | -26.54000 | repression |  |
| PA14\_62380 | NA | hypothetical protein |  |  | -26.68019 | no impact |  |
| PA14\_62420 | NA | hypothetical protein |  |  | -26.09600 | no impact |  |
| PA14\_62440 | NA | transporter | membrane, transmembrane transporter activity, transmembrane transport, nucleobase transmembrane transporter activity |  | -26.18425 | repression |  |
| PA14\_62650 | NA | hypothetical protein |  |  | -26.43604 | repression |  |
| PA14\_62660 | NA | hypothetical protein |  |  | -26.32516 | repression |  |
| PA14\_62810 | secG | preprotein translocase subunit SecG | protein secretion, P-P-bond-hydrolysis-driven protein transmembrane transporter activity, integral component of membrane | Bacterial secretion system, Protein export, Quorum sensing | -26.54087 | no impact |  |
| PA14\_62840 | glmM | phosphoglucosamine mutase | carbohydrate metabolic process, intramolecular transferase activity, phosphotransferases, magnesium ion binding, organic substance metabolic process, phosphoglucosamine mutase activity, phosphomannomutase activity, phosphoglucomutase activity, peptidoglycan biosynthetic process | Amino sugar and nucleotide sugar metabolism, Biosynthesis of antibiotics, Metabolic pathways | -26.22505 | repression |  |
| PA14\_62860 | ftsH | cell division protein FtsH | metalloendopeptidase activity, ATP binding, proteolysis, membrane, zinc ion binding, integral component of membrane, cellular response to antibiotic |  | -26.31823 | repression |  |
| PA14\_62970 | dnaK | molecular chaperone DnaK | ATP binding, protein folding, unfolded protein binding | RNA degradation | -26.70614 | repression |  |
| PA14\_63010 | recN | DNA repair protein RecN | ATP binding, DNA repair, DNA recombination |  | -26.73269 | repression |  |
| PA14\_63070 | NA | GntR family transcriptional regulator | DNA-binding transcription factor activity, regulation of transcription, DNA-templated |  | -26.37493 | repression |  |
| PA14\_63150 | pmrA | two-component response regulator | phosphorelay signal transduction system, DNA binding, regulation of transcription, DNA-templated | Quorum sensing, Two-component system | -26.02368 | no impact |  |
| PA14\_63210 | NA | two-component response regulator | phosphorelay signal transduction system |  | -26.55538 | repression |  |
| PA14\_63430 | NA | hypothetical protein |  |  | -26.99795 | repression |  |
| PA14\_63470 | NA | methyltransferase | methyltransferase activity |  | -26.09983 | repression |  |
| PA14\_63480 | NA | amino acid permease | membrane, transmembrane transporter activity, transmembrane transport | Quorum sensing | -25.87824 | repression |  |
| PA14\_63850 | lpd3 | dihydrolipoamide dehydrogenase | oxidoreductase activity, oxidation-reduction process, electron transfer activity, cell redox homeostasis, flavin adenine dinucleotide binding, dihydrolipoyl dehydrogenase activity, oxidoreductase activity, acting on a sulfur group of donors, NAD(P) as acceptor | Biosynthesis of antibiotics, Biosynthesis of secondary metabolites, Carbon metabolism, Citrate cycle (TCA cycle), Glycine, serine and threonine metabolism, Glycolysis / Gluconeogenesis, Glyoxylate and dicarboxylate metabolism, Metabolic pathways, Microbial metabolism in diverse environments, Propanoate metabolism, Pyruvate metabolism, Valine, leucine and isoleucine degradation | -26.72344 | repression |  |
| PA14\_63990 | speA | arginine decarboxylase | catalytic activity, arginine catabolic process, spermidine biosynthetic process, arginine decarboxylase activity, putrescine biosynthetic process from arginine | Arginine and proline metabolism, Metabolic pathways | -26.41367 | repression |  |
| PA14\_64030 | NA | hypothetical protein |  |  | -26.75446 | repression |  |
| PA14\_64280 | NA | branched-chain amino acid ABC transporter permease | membrane, transmembrane transporter activity, transmembrane transport | ABC transporters | -26.31395 | repression |  |
| PA14\_64290 | NA | ABC transporter permease | membrane, transmembrane transporter activity, transmembrane transport | ABC transporters | -26.50258 | repression |  |
| PA14\_64310 | NA | ABC transporter ATP-binding protein | ATP binding, ATPase activity | ABC transporters | -25.89234 | repression |  |
| PA14\_64540 | NA | hypothetical protein | nitrate assimilation |  | -26.68686 | repression |  |
| PA14\_64690 | NA | transmembrane sensor |  |  | -26.02169 | no impact |  |
| PA14\_64790 | NA | MFS transporter | integral component of membrane, transmembrane transporter activity, transmembrane transport, integral component of plasma membrane |  | -25.86214 | repression |  |
| PA14\_64860 | NA | ABC transporter ATP-binding protein | ATP binding, ATPase activity, branched-chain amino acid transmembrane transporter activity, branched-chain amino acid transport | ABC transporters, Quorum sensing | -25.90276 | no impact |  |
| PA14\_64870 | NA | ABC transporter ATP-binding protein | ATP binding, ATPase activity | ABC transporters, Quorum sensing | -25.82008 | repression |  |
| PA14\_64890 | NA | branched chain amino acid ABC transporter permease | membrane, transmembrane transporter activity, transmembrane transport | ABC transporters, Quorum sensing | -26.07133 | repression |  |
| PA14\_64900 | NA | ABC transporter substrate-binding protein | amino acid transport | ABC transporters, Quorum sensing | -26.79778 | repression |  |
| PA14\_64930 | NA | hypothetical protein | hydrolase activity |  | -26.41814 | repression |  |
| PA14\_65040 | NA | hypothetical protein | membrane, transmembrane transport |  | -26.98324 | repression |  |
| PA14\_65150 | rplI | 50S ribosomal protein L9 | structural constituent of ribosome, ribosome, translation | Ribosome | -26.32846 | repression |  |
| PA14\_65230 | purA | adenylosuccinate synthetase | GTP binding, adenylosuccinate synthase activity, purine nucleotide biosynthetic process | Alanine, aspartate and glutamate metabolism, Metabolic pathways, Purine metabolism | -26.94784 | repression |  |
| PA14\_65250 | hisZ | ATP phosphoribosyltransferase | histidine biosynthetic process, cytoplasm | Biosynthesis of amino acids, Biosynthesis of secondary metabolites, Histidine metabolism, Metabolic pathways | -26.48545 | no impact |  |
| PA14\_65270 | hflC | protease subunit HflC | membrane, integral component of membrane, regulation of peptidase activity |  | -26.00770 | repression |  |
| PA14\_65300 | hflX | GTP-binding protein | GTP binding |  | -26.08092 | no impact |  |
| PA14\_65310 | hfq | RNA-binding protein Hfq | RNA binding, regulation of transcription, DNA-templated | Quorum sensing, RNA degradation | -26.44463 | repression |  |
| PA14\_65370 | amiB | N-acetylmuramoyl-L-alanine amidase | N-acetylmuramoyl-L-alanine amidase activity, peptidoglycan catabolic process | Cationic antimicrobial peptide (CAMP) resistance | -26.47295 | repression |  |
| PA14\_65390 | NA | hypothetical protein | ADP-dependent NAD(P)H-hydrate dehydratase activity | NADH repair | -27.08775 | repression |  |
| PA14\_65500 | psd | phosphatidylserine decarboxylase | phosphatidylserine decarboxylase activity, phospholipid biosynthetic process | Biosynthesis of secondary metabolites, Glycerophospholipid metabolism, Metabolic pathways | -25.78809 | repression |  |
| PA14\_65740 | thiC | thiamine biosynthesis protein ThiC | thiamine biosynthetic process, iron-sulfur cluster binding, carbon-carbon lyase activity | Metabolic pathways, Thiamine metabolism | -25.91016 | repression |  |
| PA14\_65795 | NA | hypothetical protein | magnesium ion binding, thiamine pyrophosphate binding, catalytic activity | Arginine and proline metabolism, Metabolic pathways | -26.33738 | no impact |  |
| PA14\_65820 | NA | acyl-CoA dehydrogenase | acyl-CoA dehydrogenase activity, oxidation-reduction process, oxidoreductase activity, acting on the CH-CH group of donors, flavin adenine dinucleotide binding |  | -26.37244 | no impact |  |
| PA14\_65900 | NA | TetR family transcriptional regulator | DNA binding |  | -26.25507 | repression |  |
| PA14\_65970 | NA | transcriptional regulator | DNA-binding transcription factor activity, regulation of transcription, DNA-templated |  | -26.51420 | repression |  |
| PA14\_66110 | NA | glycosyl transferase family protein |  |  | -26.56124 | repression |  |
| PA14\_66120 | NA | hypothetical protein |  |  | -26.01147 | repression |  |
| PA14\_66160 | NA | glycosyl transferase family protein |  |  | -26.22993 | repression |  |
| PA14\_66210 | NA | hypothetical protein | protein kinase activity, protein phosphorylation |  | -26.40993 | repression |  |
| PA14\_66250 | waaF | heptosyltransferase II | lipopolysaccharide biosynthetic process, transferase activity, transferring glycosyl groups | Lipopolysaccharide biosynthesis, Metabolic pathways | -26.36280 | repression |  |
| PA14\_66340 | NA | hypothetical protein | rRNA methyltransferase activity, rRNA base methylation |  | -25.91521 | no impact |  |
| PA14\_66620 | pilQ | type 4 fimbrial biogenesis outer membrane protein PilQ precursor | outer membrane, protein secretion |  | -26.43861 | repression |  |
| PA14\_66640 | pilO | type 4 fimbrial biogenesis protein PilO | type IV pilus-dependent motility, type IV pilus biogenesis |  | -26.45195 | repression |  |
| PA14\_66690 | NA | protease | metalloendopeptidase activity, proteolysis |  | -26.59438 | repression |  |
| PA14\_66840 | phaC2 | poly(3-hydroxyalkanoic acid) synthase 2 | poly-hydroxybutyrate biosynthetic process, transferase activity, transferring acyl groups | Butanoate metabolism | -26.08728 | repression |  |
| PA14\_66890 | NA | hypothetical protein |  |  | -26.07996 | no impact |  |
| PA14\_66970 | tatB | sec-independent translocase | membrane, protein transport by the Tat complex, protein transport | Bacterial secretion system, Protein export | -25.81061 | repression |  |
| PA14\_67110 | NA | prolyl aminopeptidase | proteolysis, peptidase activity, aminopeptidase activity, cytoplasm | Arginine and proline metabolism | -25.81514 | repression |  |
| PA14\_67170 | NA | LysR family transcriptional regulator | DNA-binding transcription factor activity, regulation of transcription, DNA-templated |  | -25.98541 | repression |  |
| PA14\_67300 | NA | ABC transporter substrate-binding protein | transmembrane transporter activity, ATP-binding cassette (ABC) transporter complex, transmembrane transport | ABC transporters | -26.70841 | repression |  |
| PA14\_67310 | NA | amino acid permease | amino acid transport, integral component of membrane, transmembrane transport, membrane, transmembrane transporter activity |  | -26.57136 | repression |  |
| PA14\_67530 | NA | hypothetical protein |  |  | -26.40825 | repression |  |
| PA14\_67540 | NA | hypothetical protein |  |  | -26.14696 | no impact |  |
| PA14\_67550 | NA | transcriptional regulator | DNA binding, regulation of transcription, DNA-templated |  | -25.80983 | repression |  |
| PA14\_67620 | NA | hypothetical protein |  |  | -26.00664 | repression |  |
| PA14\_67700 | NA | hypothetical protein |  |  | -25.92493 | repression |  |
| PA14\_67770 | pgm | phosphoglyceromutase | catalytic activity, phosphoglycerate mutase activity, cytoplasm, glucose catabolic process, manganese ion binding, metal ion binding | 1-butanol autotrophic biosynthesis (engineered), Biosynthesis of amino acids, Biosynthesis of antibiotics, Biosynthesis of secondary metabolites, Carbon metabolism, Entner-Doudoroff pathway III (semi-phosphorylative), ethylene biosynthesis V (engineered), gluconeogenesis II (<i>Methanobacterium thermoautotrophicum</i>), glycerol degradation to butanol, Glycine, serine and threonine metabolism, Glycine, serine and threonine metabolism, Glycolysis / Gluconeogenesis, Glycolysis / Gluconeogenesis, glycolysis II (from fructose 6-phosphate), glycolysis IV (plant cytosol), Metabolic pathways, Methane metabolism, Methane metabolism, Microbial metabolism in diverse environments, photosynthetic 3-hydroxybutanoate biosynthesis (engineered), Rubisco shunt, superpathway of glucose and xylose degradation | -26.11933 | repression |  |
| PA14\_67840 | NA | ABC-type amino acid transporter |  |  | -25.93803 | repression |  |
| PA14\_67990 | mutY | A/G-specific adenine glycosylase | base-excision repair, DNA binding, catalytic activity, DNA N-glycosylase activity, hydrolase activity, DNA repair | Base excision repair | -26.32967 | repression |  |
| PA14\_68040 | NA | short-chain dehydrogenase | oxidoreductase activity |  | -26.28830 | repression |  |
| PA14\_68110 | NA | transcriptional regulator | DNA-binding transcription factor activity, regulation of transcription, DNA-templated |  | -25.78841 | repression |  |
| PA14\_68200 | rmlA | glucose-1-phosphate thymidylyltransferase | glucose-1-phosphate thymidylyltransferase activity, extracellular polysaccharide biosynthetic process, biosynthetic process, nucleotidyltransferase activity | Acarbose and validamycin biosynthesis, Biosynthesis of antibiotics, Metabolic pathways, Polyketide sugar unit biosynthesis, Streptomycin biosynthesis | -26.04083 | repression |  |
| PA14\_68340 | arcB | ornithine carbamoyltransferase | cellular amino acid metabolic process, amino acid binding, carboxyl- or carbamoyltransferase activity, ornithine metabolic process, ornithine carbamoyltransferase activity, arginine deiminase pathway | Arginine and proline metabolism; Urea cycle and metabolism of amino groups, Arginine biosynthesis, Biosynthesis of amino acids, Biosynthesis of antibiotics, Biosynthesis of secondary metabolites, Metabolic pathways | -25.85066 | repression |  |
| PA14\_68380 | nudE | ADP-ribose diphosphatase NudE | hydrolase activity | Purine metabolism | -25.92373 | repression |  |
| PA14\_68400 | NA | LysM domain/BON superfamily protein |  |  | -27.58552 | repression |  |
| PA14\_68470 | NA | hypothetical protein | RNA binding, regulation of carbohydrate metabolic process, mRNA catabolic process |  | -26.85627 | repression |  |
| PA14\_68500 | NA | iron-containing alcohol dehydrogenase | oxidoreductase activity, metal ion binding, oxidation-reduction process |  | -25.87900 | repression |  |
| PA14\_68570 | NA | hypothetical protein |  |  | -26.37132 | repression |  |
| PA14\_68620 | NA | hypothetical protein |  |  | -26.61808 | repression |  |
| PA14\_68830 | NA | hypothetical protein |  |  | -26.66519 | repression |  |
| PA14\_68870 | gcvT | glycine cleavage system aminomethyltransferase T | aminomethyltransferase activity, glycine catabolic process, protein binding | Biosynthesis of antibiotics, Biosynthesis of secondary metabolites, Carbon metabolism, Glycine, serine and threonine metabolism, Glycine, serine and threonine metabolism, Glyoxylate and dicarboxylate metabolism, Metabolic pathways, One carbon pool by folate, One carbon pool by folate | -25.78242 | repression |  |
| PA14\_68920 | NA | LysR family transcriptional regulator | DNA-binding transcription factor activity, regulation of transcription, DNA-templated |  | -27.18480 | repression |  |
| PA14\_68955 | NA | 2-octaprenyl-3-methyl-6-methoxy-1,4-benzoquinol hydroxylase | ubiquinone biosynthetic process, oxidoreductase activity, acting on paired donors, with incorporation or reduction of molecular oxygen, NAD(P)H as one donor, and incorporation of one atom of oxygen, flavin adenine dinucleotide binding, oxidation-reduction process, FAD binding | Biosynthesis of secondary metabolites, Metabolic pathways, Ubiquinone and other terpenoid-quinone biosynthesis | -25.90195 | repression |  |
| PA14\_69090 | NA | hypothetical protein |  |  | -26.29489 | repression |  |
| PA14\_69140 | NA | CDP-6-deoxy-delta-3,4-glucoseen reductase | electron transfer activity, iron-sulfur cluster binding, oxidoreductase activity, oxidation-reduction process | Amino sugar and nucleotide sugar metabolism | -26.24278 | repression |  |
| PA14\_69170 | NA | O-antigen acetylase | transferase activity, transferring acyl groups other than amino-acyl groups |  | -26.66883 | repression |  |
| PA14\_69230 | ppk | polyphosphate kinase | polyphosphate biosynthetic process, polyphosphate kinase activity, polyphosphate kinase complex | Oxidative phosphorylation, Oxidative phosphorylation, RNA degradation | -26.01586 | no impact |  |
| PA14\_69320 | NA | integral membrane transport protein | integral component of membrane |  | -25.84492 | repression |  |
| PA14\_69330 | NA | hypothetical protein | membrane, transmembrane transport |  | -26.36819 | repression |  |
| PA14\_69390 | algQ | anti-RNA polymerase sigma 70 factor | positive regulation of secondary metabolite biosynthetic process, negative regulation of proteolysis, positive regulation of single-species biofilm formation, regulation of transcription, DNA-templated |  | -26.26350 | repression |  |
| PA14\_69620 | NA | hypothetical protein |  |  | -25.91965 | repression |  |
| PA14\_69670 | lysA | diaminopimelate decarboxylase | catalytic activity, lysine biosynthetic process via diaminopimelate, diaminopimelate decarboxylase activity | Biosynthesis of amino acids, Biosynthesis of antibiotics, Biosynthesis of secondary metabolites, Lysine biosynthesis, Metabolic pathways, Microbial metabolism in diverse environments | -26.64274 | repression |  |
| PA14\_69850 | NA | choline transporter | membrane, transmembrane transporter activity, nitrogen compound transport |  | -25.78194 | repression |  |
| PA14\_69870 | pchP | phosphorylcholine phosphatase |  |  | -27.14901 | repression |  |
| PA14\_70060 | NA | lipoprotein |  |  | -26.34428 | repression |  |
| PA14\_70070 | NA | hypothetical protein |  |  | -26.82594 | repression |  |
| PA14\_70190 | rpmB | 50S ribosomal protein L28 | structural constituent of ribosome, ribosome, translation | Ribosome | -25.96308 | no impact |  |
| PA14\_70230 | radC | DNA repair protein RadC |  |  | -26.88931 | repression |  |
| PA14\_70270 | algC | phosphomannomutase | carbohydrate metabolic process, intramolecular transferase activity, phosphotransferases, magnesium ion binding, organic substance metabolic process, pathogenesis, phosphoglucomutase activity, lipopolysaccharide core region biosynthetic process, phosphomannomutase activity, alginic acid biosynthetic process | Amino sugar and nucleotide sugar metabolism, Amino sugar and nucleotide sugar metabolism, Biosynthesis of antibiotics, Biosynthesis of secondary metabolites, CMP-legionaminate biosynthesis I, Fructose and mannose metabolism, Galactose metabolism, Glycolysis / Gluconeogenesis, Metabolic pathways, Microbial metabolism in diverse environments, Pentose phosphate pathway, Purine metabolism, Starch and sucrose metabolism, Streptomycin biosynthesis | -26.44743 | no impact |  |
| PA14\_70360 | NA | hypothetical protein |  |  | -26.70193 | repression |  |
| PA14\_70470 | spoT | guanosine-3',5'-bis(diphosphate) 3'-pyrophosphohydrolase | guanosine tetraphosphate metabolic process | Purine metabolism | -25.99657 | repression |  |
| PA14\_70490 | NA | lipoprotein |  |  | -26.94896 | no impact |  |
| PA14\_70600 | NA | HU family DNA-binding protein | DNA binding |  | -26.12252 | no impact |  |
| PA14\_70650 | NA | GlcG protein |  |  | -26.19697 | repression |  |
| PA14\_70760 | phoR | two-component sensor PhoR | phosphorelay sensor kinase activity, phosphorelay signal transduction system, protein histidine kinase activity, integral component of membrane, regulation of transcription, DNA-templated, signal transduction, phosphorylation, transferase activity, transferring phosphorus-containing groups, phosphoprotein phosphatase activity | Two-component system | -26.07836 | repression |  |
| PA14\_71060 | sdaB | L-serine dehydratase | L-serine ammonia-lyase activity, gluconeogenesis, 4 iron, 4 sulfur cluster binding | Biosynthesis of amino acids, Biosynthesis of antibiotics, Biosynthesis of secondary metabolites, Carbon metabolism, Cysteine and methionine metabolism, Cysteine and methionine metabolism, Glycine, serine and threonine metabolism, Glycine, serine and threonine metabolism, Metabolic pathways | -25.78747 | repression |  |
| PA14\_71160 | NA | hypothetical protein | choline transport, choline binding, periplasmic space, transmembrane transporter activity, ATP-binding cassette (ABC) transporter complex, transmembrane transport | ABC transporters | -26.08623 | repression |  |
| PA14\_71250 | NA | hypothetical protein | glycine betaine catabolic process, choline catabolic process |  | -26.07017 | repression |  |
| PA14\_71280 | NA | ferredoxin | iron-sulfur cluster binding, choline catabolic process, glycine betaine catabolic process |  | -25.76588 | repression |  |
| PA14\_71390 | NA | hypothetical protein |  |  | -26.50089 | no impact |  |
| PA14\_71410 | NA | ring hydroxylating dioxygenase, alpha-subunit | oxidoreductase activity, 2 iron, 2 sulfur cluster binding, oxidation-reduction process, iron ion binding, cellular metabolic process |  | -26.05708 | repression |  |
| PA14\_71430 | NA | hypothetical protein |  |  | -26.43930 | repression |  |
| PA14\_71460 | glyA1 | serine hydroxymethyltransferase | pyridoxal phosphate binding, glycine hydroxymethyltransferase activity, glycine biosynthetic process from serine, tetrahydrofolate interconversion, catalytic activity | Biosynthesis of amino acids, Biosynthesis of antibiotics, Biosynthesis of secondary metabolites, Carbon metabolism, Cyanoamino acid metabolism, Glycine, serine and threonine metabolism, Glyoxylate and dicarboxylate metabolism, Metabolic pathways, Methane metabolism, Microbial metabolism in diverse environments, One carbon pool by folate | -26.53998 | no impact |  |
| PA14\_71490 | soxD | sarcosine oxidase delta subunit | sarcosine oxidase activity, tetrahydrofolate metabolic process, sarcosine catabolic process | Glycine, serine and threonine metabolism, Metabolic pathways | -26.21340 | repression |  |
| PA14\_71510 | soxG | sarcosine oxidase gamma subunit | protein binding, sarcosine oxidase activity, sarcosine catabolic process | Glycine, serine and threonine metabolism, Metabolic pathways | -26.72305 | repression |  |
| PA14\_71590 | NA | hypothetical protein | integral component of membrane |  | -26.73632 | repression |  |
| PA14\_71620 | purE | phosphoribosylaminoimidazole carboxylase catalytic subunit | 'de novo' IMP biosynthetic process | Biosynthesis of antibiotics, Biosynthesis of secondary metabolites, Metabolic pathways, Purine metabolism | -25.92860 | no impact |  |
| PA14\_71630 | adhA | alcohol dehydrogenase | oxidoreductase activity, zinc ion binding, oxidation-reduction process | Biosynthesis of antibiotics, Biosynthesis of secondary metabolites, Chloroalkane and chloroalkene degradation, Degradation of aromatic compounds, Fatty acid degradation, Glycolysis / Gluconeogenesis, Metabolic pathways, Microbial metabolism in diverse environments, Naphthalene degradation, Tyrosine metabolism | -25.94997 | repression |  |
| PA14\_71900 | NA | hypothetical protein |  |  | -25.87426 | repression |  |
| PA14\_71990 | gmd | GDP-mannose 4,6-dehydratase | GDP-mannose 4,6-dehydratase activity, GDP-mannose metabolic process | Amino sugar and nucleotide sugar metabolism, Fructose and mannose metabolism, Metabolic pathways | -26.43427 | no impact |  |
| PA14\_72010 | NA | glycosyltransferase | lipopolysaccharide biosynthetic process |  | -26.07097 | repression |  |
| PA14\_72210 | NA | hypothetical protein | RNA processing, RNA ligase activity | beta-Lactam resistance | -25.81286 | repression |  |
| PA14\_72340 | gltP | glutamate/aspartate:proton symporter | symporter activity, integral component of membrane, dicarboxylic acid transport |  | -25.91665 | repression |  |
| PA14\_72370 | NA | hypothetical protein | plasma membrane |  | -26.08570 | repression |  |
| PA14\_72380 | algB | two-component response regulator AlgB | sequence-specific DNA binding, DNA binding, phosphorelay signal transduction system, ATP binding, regulation of transcription, DNA-templated, transcription factor binding, alginic acid biosynthetic process, positive regulation of transcription, DNA-templated, negative regulation of single-species biofilm formation on inanimate substrate, transcription regulatory region DNA binding, phosphorelay response regulator activity, phosphorelay sensor kinase activity, positive regulation of proteolysis, positive regulation of secondary metabolite biosynthetic process, positive regulation of cell motility | Two-component system | -27.21684 | repression |  |
| PA14\_72520 | NA | hypothetical protein |  |  | -25.89468 | no impact |  |
| PA14\_72580 | znuC | zinc transporter | ATP binding, ATPase activity, plasma membrane, zinc ion transport, ATPase-coupled zinc transmembrane transporter activity, response to zinc ion | ABC transporters | -26.03917 | repression |  |
| PA14\_72740 | NA | two-component sensor | phosphorelay sensor kinase activity, signal transduction, phosphorylation, transferase activity, transferring phosphorus-containing groups | Two-component system | -26.43810 | no impact |  |
| PA14\_72760 | NA | beta-lactamase | penicillin binding, beta-lactam antibiotic catabolic process, beta-lactamase activity | beta-Lactam resistance | -26.58347 | repression |  |
| PA14\_72770 | NA | hypothetical protein |  |  | -26.09289 | repression |  |
| PA14\_72810 | NA | hypothetical protein |  |  | -25.93243 | no impact |  |
| PA14\_72960 | NA | MFS dicarboxylate transporter | integral component of membrane, transmembrane transporter activity, transmembrane transport, integral component of plasma membrane |  | -25.83824 | repression |  |
| PA14\_73150 | NA | hypothetical protein |  |  | -26.92076 | repression |  |
| PA14\_73200 | NA | hypothetical protein |  |  | -27.04344 | repression |  |
| PA14\_73350 | soj | chromosome partitioning protein Soj |  |  | -26.12023 | repression |  |
